# The perfect condition for the rising of superbugs: person-to-person contagion and antibiotic use are the key factors responsible for the positive correlation between antibiotic resistance gene diversity and virulence gene diversity in human metagenomes

**DOI:** 10.1101/2020.04.25.061853

**Authors:** Célia P. F. Domingues, João S. Rebelo, Teresa Nogueira, Joël Pothier, Francisca Monteiro, Francisco Dionisio

**Author notes:** These authors have contributed equally to this work (Célia P. F. Domingues, João S. Rebelo). **Correspondence:** Francisco Dionisio.

## Abstract

This study aims to understand the cause of the recent observation that humans with a higher diversity of virulence genes in their metagenomes tend to be precisely those with higher diversity of antibiotic-resistance genes. We simulated the transferring of virulence and antibiotic-resistance genes in a community of interacting people where some take antibiotics. The diversities of the two genes types became positively correlated whenever the contagion probability between two people was higher than the probability of losing resistant genes. However, no such positive correlations arise if no one takes antibiotics. This finding holds even under changes of several simulations’ parameters, such as the relative or total diversity of virulence and resistance genes, the contagion probability between individuals, the loss rate of resistance genes, or the social network type. Because the loss rate of resistance genes may be shallow, we conclude that the contagion between people and antibiotic usage is the leading cause of establishing the positive correlation mentioned above. Therefore, antibiotic use and something as prosaic as the contagion between people may facilitate the emergence of virulent and multi-resistant bacteria in people’s metagenomes with a high diversity of both gene types. These superbugs may then circulate in the community.

## 2. Introduction

Since the 1940s, antibiotics have been used in health contexts in medicine and veterinary and as growth promoters in livestock and agriculture (Castanon, 2007). As an incredible example of Darwinian selection, bacteria worldwide have gradually become resistant to several antibiotics. Such spread of resistance has had terrible consequences. For example, there were about 875 500 disability-adjusted life-years and more than 33 000 deaths in European Economic Area due to antibiotic resistance in 2015 (Cassini et al., 2019).

Bacterial communities are often very complex, eventually comprising both pathogenic and non-pathogenic bacteria. The human microbiome, defined as the set of microorganisms that colonize humans (body’s surfaces and biofluids, including tissues such as skin, mucosa, and, most importantly, the gastrointestinal tract) comprises about 3.8◻×◻10^13^ bacterial cells (Sender et al., 2016), spanning thousands of taxa.

Virulence factors are proteins that help bacteria in colonizing a host or biome. These traits are easily spread in bacterial populations or microbiomes by horizontal gene transfer, which can potentially convert mutualistic or commensal bacteria into pathogens able to progress into new tissues, triggering an infectious disease. We recently found a positive correlation between antibiotic resistance genes’ diversity and virulence genes’ diversity across human gut microbiomes (Escudeiro et al., 2019). Could this positive correlation result from administering antibiotics in sick people due to bacterial infections, eventually selecting bacteria encoding virulence and resistance determinants simultaneously? This hypothesis is unlikely to be adequate because, even when the objective of taking antibiotics is to kill or inhibit the growth of pathogenic bacteria, many non-pathogenic (mutualistic or commensal) strains and species are undoubtedly affected. Therefore, an explanation for the positive correlation mentioned above is still missing.

Both virulence and resistance genes present in commensal bacteria and pathogenic bacteria spread between people’s metagenomes. This dissemination may contribute to the accumulation of virulence and resistance genes in some people when themselves or their contacts take antibiotics. Meanwhile, pathogenic bacteria’s presence triggers the administration of antibiotics. Therefore, contagion (the dissemination of bacteria and their genes) between people should play a role in keeping the correlation between resistance and virulence genes’ diversity. Microbes’ transmission from mother to child is already well documented (Blaser and Falkow, 2009; Nayfach et al., 2016; Ferretti et al., 2018; Yassour et al., 2018; Nogueira et al., 2019). A recent study highlighted that the oral and gut microbiomes of people belonging to the same household share similarities in bacterial strains, regardless of these people’s genetic relationship (Brito et al., 2019). These studies suggest that bacteria in human microbiomes can have a shared exposure or result from person to person transfer on the social network. This suggestion is supported by a study that showed that social interactions shape the chimpanzee’s microbiomes (Moeller et al., 2016).

This work aims to find the key factors leading to the positive correlation between the diversity of virulence and antibiotic resistance genes observed across human metagenomes (Escudeiro et al., 2019). To this end, we simulated the transfer of bacterial pathogens and antibiotic resistance and virulence genes in a human-to-human transmission network. We show that a positive correlation between the diversity of antibiotic resistance coding genes and those coding for virulence emerges whenever the contagion rate between individuals is higher than the probability that metagenomes lose resistant genes, independently of all the other parameters of the simulations. This simple rule explains the positive correlation between virulence genes’ diversity and antibiotic resistance genes’ diversity.

## 3. Methods

### 2.1 Building the human network

We simulated a network where each node represents a person or, more precisely, a person’s metagenome. To simplify language, sometimes we use the words person or people, meaning a person’s metagenome or people’s metagenomes, respectively. The edges represent possible transmission avenues of microorganisms.

We built the social contact network following the Watts and Strogatz method (Watts and Strogatz, 1998). In a regular network, each node links to the *n* nearest nodes. In non-regular networks, each node’s link has a certain probability *p* of being reconnected to another randomly chosen node. The parameter *p* represents the probability of each connection to be modified. We defined the network type by the value assigned to the parameter *p* (for example, a regular network when *p* = 0, whereas *p* = 1 results in a random network). Unless noted, we performed simulations with *p* = 0.5.

### 2.2 The metagenome, pathogenic bacteria, and antibiotic administration

The model considers the transmission of bacterial pathogens (capable of causing infections), as well as virulence and antibiotic resistance genes, between people. These non-housekeeping genes are present in the metagenome. We focused on the presence or absence of genes encoding different functions, irrespectively of its copy-number in the metagenome. In the simulations, each gene represents a gene family (with similar functions). We divided resistance genes into groups, each group having the same number of families. Each group represents genes associated with resistance to an antibiotic. Of note, we did not consider resistance to multiple drugs in our simulations. Therefore, there will be as many groups as there are antibiotics accounted for in the simulations. We define the diversity of a specific gene kind as the number of genes of that type present in a human metagenome.

To simulate the migration of bacteria from individuals outside the network or the contagion from sources such as food or contaminated water, we inserted five different bacterial pathogenic species into random individuals per cycle. To simplify language, we assume that pathogenic bacteria belong to different species, but, in reality, some of them may constitute different strains of the same species. In this model, the only difference between species is the antibiotic to which they are susceptible, as explained below.

Individuals infected by pathogenic bacteria feel sick and take an antibiotic. The antibiotic administered is specific for the bacteria that caused the illness. The antibiotic selects cells carrying resistance genes by eliminating the remaining susceptible bacteria. In this work, we assume that all families of resistance genes are present in all metagenomes, but in two different possible states: in some metagenomes, they are present in low copy number, so they are not transmissible to other individuals in the network; in other metagenomes, the copy number of resistance genes is high due to the selective pressure of antibiotics to which they were previously submitted. In the latter case, resistance genes are transmissible from person to person.

Moreover, upon antibiotic consumption, the following events can occur: (i) elimination of a pathogenic bacterial species; (ii) selection in the metagenome of resistance genes belonging to the same group of resistance to an antibiotic, which means their copy number gets so high that they become transferable; (iii) loss of resistance genes associated with other antibiotics with a given probability (becoming non-transferable but still present in minute copy number); (iv) virulence genes disappear from the metagenome with a given probability.

Several processes lead to gene loss. Genes are lost because of the selective pressure by antibiotics and because we assume that resistance determinants impose a fitness cost (in the absence of antibiotics). To include this cost in the simulations, we consider that each metagenome may lose specific resistance genes according to a “loss rate” (with this process, these genes become non-transferable).

### 2.3 Algorithm of the program

Each simulation is composed of several cycles. In each cycle, we considered all procedures described in the pseudocode (Table 1; see also the flowchart in Fig. 1). We performed exploratory simulations to parameterize our model. We fixed a set of parameters as default (Table 2). The main steps of the program in each cycle are:

i. Transfer of pathogenic bacteria, virulence and resistance genes between people (i.e., between linked nodes), according to specific contagion probabilities of pathogens and virulence and resistance genes of the metagenome. With this process, the diversity of genes present in the recipient metagenome increases.
ii. To look for people infected by at least one pathogenic bacterial species. These people take antibiotics (chosen according to the pathogen). The antibiotic eliminates the pathogenic species and selects the resistance genes associated with the antibiotic used. According to a certain probability, the antibiotic also eliminates virulence genes and resistance genes unrelated to the administered antibiotic. Finally, the metagenome loses a few more resistance genes not associated with the antibiotic, according to the loss rate probability. The cause of this loss is the fitness cost of resistance genes.
iii. The metagenomes of people that did not take an antibiotic in this cycle lose resistance genes according to the loss rate probability. This loss is a consequence of the fitness cost imposed by resistance genes on their hosts, which is not happening with virulence genes.
iv. Insert the five bacterial pathogenic species in five individuals randomly chosen from the population.

**Table 1.**
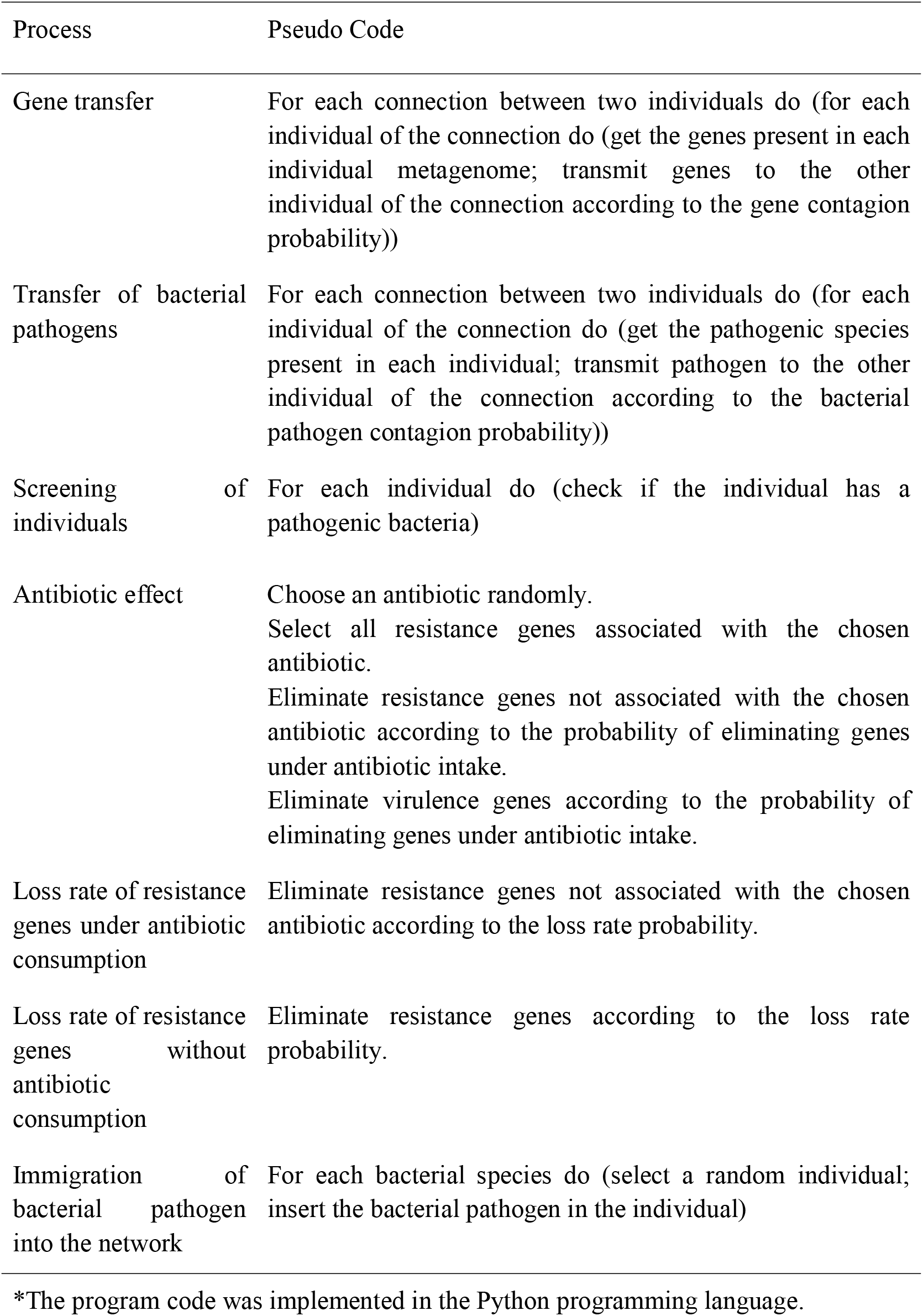
Pseudocode of the program*.

| Process | Pseudo Code |
| --- | --- |
| Gene transfer | For each connection between two individuals do (for each individual of the connection do (get the genes present in each individual metagenome; transmit genes to the other individual of the connection according to the gene contagion probability)) |
| Transfer of bacterial pathogens | For each connection between two individuals do (for each individual of the connection do (get the pathogenic species present in each individual; transmit pathogen to the other individual of the connection according to the bacterial pathogen contagion probability)) |
| Screening of individuals | For each individual do (check if the individual has a pathogenic bacteria) |
| Antibiotic effect | Choose an antibiotic randomly.<br>Select all resistance genes associated with the chosen antibiotic.<br>Eliminate resistance genes not associated with the chosen antibiotic according to the probability of eliminating genes under antibiotic intake.<br>Eliminate virulence genes according to the probability of eliminating genes under antibiotic intake. |
| Loss rate of resistance genes under antibiotic consumption | Eliminate resistance genes not associated with the chosen antibiotic according to the loss rate probability. |
| Loss rate of resistance genes without antibiotic consumption | Eliminate resistance genes according to the loss rate probability. |
| Immigration of bacterial pathogen into the network | For each bacterial species do (select a random individual; insert the bacterial pathogen in the individual) |

\*The program code was implemented in the Python programming language.

**Table 2.** Parameters and default values used in simulations.

| Parameters | Default values | Changing values |
| --- | --- | --- |
| Rewiring connectivity probability $p$ | 0.5 | 0 or 1 |
| Number of individuals | 1000 | 3000 |
| Number of virulence genes | 100 | 200, 400 |
| Number of resistance genes | 100 | 200, 400 |
| Number of pathogenic bacterial species | 5 | NA |
| Number of antibiotics | 5 | NA |
| Gene contagion probability | 0.005, 0.01 | 0.0005, 0.0025, 0.015, 0.02 |
| Bacterial pathogen contagion probability | 0.15 | 0.05, 0.1, 0.2, 0.25 |
| Probability of eliminating genes under antibiotic intake | 0.7 | 0.3, 0.5 |
| The loss rate of resistance genes | 0, 0.005, 0.01, 0.015, 0.02, 0.025, 0.03 | NA |

**Figure 1.**
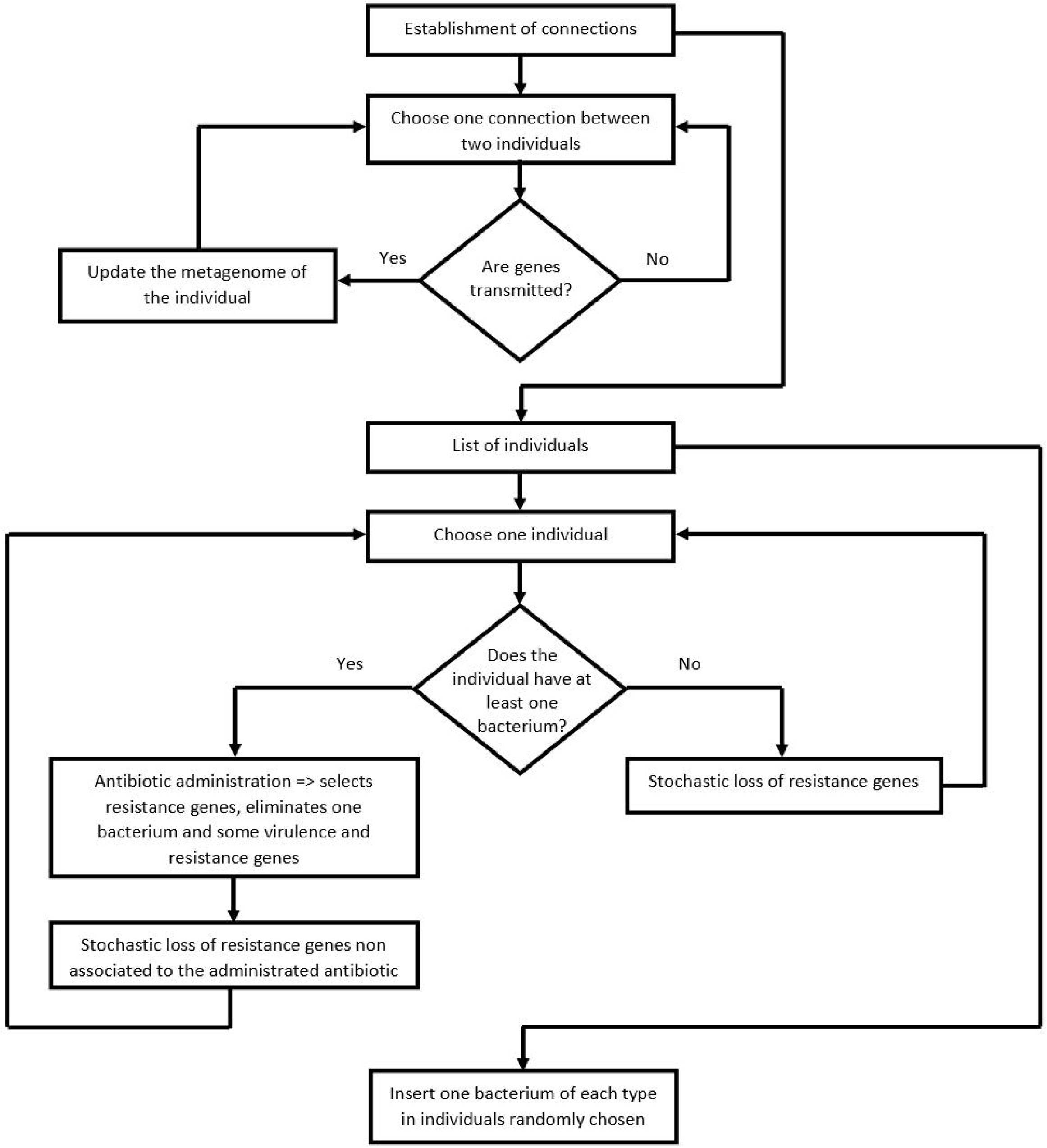
Flowchart of the program. After the network’s construction, the program performs several cycles where, eventually, there is gene transfer between nodes (people). Some individuals get sick and take antibiotics. Some genes are lost due to antibiotic pressure or the fitness cost imposed by resistance genes.

### 2.4 Statistical analysis

We considered that *Y* (diversity of resistance genes) correlates with *X* (diversity of virulence genes), according to:

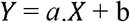

In this equation, the parameter *a* represents the linear regression slope, while *b* represents the point at which the line crosses the y-axis.

Given the complexity of human interactions, it is paramount to simplify the computer simulations. A simplified model allows us to comprehend the effect of specific factors in our simulations, which would otherwise be extremely difficult to detect. As these simplifications do not allow us to make quantitative inferences, we make qualitative analyses. The focus is always on the correlation or linear regression slope signal between the diversity of virulence and antibiotic resistance genes and whether the correlation is significantly different from zero. Accordingly, the null hypothesis is that there is no correlation between antibiotic resistance genes’ diversity and virulence genes’ diversity. The alternative hypothesis is that there is a correlation between antibiotic resistance genes’ diversity and virulence genes. We define *α* = 1×10^−6^, rejecting the null hypothesis if P-value < *α*.

We performed the data analyses described above, and the Student’s t-tests (see Supplementary Information) with R – version 3.5.1 (R Core Team, 2015).

## 4. Results

### 4.1 The number of diseases and the probability of contagion

This work aims to understand the positive correlation between antibiotic resistance genes’ diversity and virulence genes in metagenomes across human populations observed by Escudeiro et al. (2019). As explained in the Methods section, we assumed that people establish a fixed network of contacts between them and that there is the transmission of pathogenic bacteria along with antibiotic-resistance and virulence genes between connected people. In the simulations, five different pathogenic bacteria, belonging to distinct species, circulate between linked people. When pathogenic bacteria infect an individual, that person takes an antibiotic. The antibiotic eliminates only the pathogenic species associated with the administered antibiotic, even if more than one species infects that individual. The antibiotic also removes a certain percentage of virulence and resistance genes.

In principle, the bacterial pathogen contagion probability parameter could have any value in the simulations. Given the importance of this parameter, we must calibrate its value according to the model’s other conditions. We assumed that individuals are not affected by more than two infectious diseases at the same time. Therefore, we started this study searching for the parameters that led individuals to have a maximum of two pathogenic bacterial species or strain simultaneously at a given cycle.

We performed simulations with different bacterial pathogen contagion probabilities, and counted the number of pathogenic bacterial species that each individual has per cycle. As we can see in Table 3, when the bacteria pathogen contagion probability is 0.2, some individuals in a specific cycle (out of two million possibilities) became infected by three pathogenic bacteria. For this reason, we settled the bacterial pathogen contagion probability to be less than 0.2 in our simulations.

**Table 3.** Number of pathogenic species according to the bacterial pathogen contagion probability.

| Bacterial pathogen contagion probability | Number of pathogenic species (in 2 000 000 possibilities) |  |  |  |  |  |
| --- | --- | --- | --- | --- | --- | --- |
|  | 0 | 1 | 2 | 3 | 4 | 5 |
| 0.05 | 1987473 | 12496 | 31 | 0 | 0 | 0 |
| 0.1 | 1982852 | 17094 | 54 | 0 | 0 | 0 |
| 0.15 | 1973053 | 26763 | 184 | 0 | 0 | 0 |
| 0.2 | 1940458 | 58759 | 779 | 4 | 0 | 0 |
| 0.25 | 104967 | 262575 | 527204 | 705479 | 399253 | 522 |

In each cycle of the simulation, we introduced five pathogenic bacterial species into the population. Then, we counted the total number of pathogenic species present in the population. If this number is equal to five, then the only pathogenic species in the population are those that were inserted (simulating immigration into the network), which means that, before the insertion, all pathogenic bacterial species had disappeared in that cycle. As it is unrealistic that all bacterial species disappear simultaneously, we looked for a contagion value below 0.2 that minimizes the number of times that all bacteria disappear at the same time. As shown in Table 4, the number of times that pathogenic species disappear increases with a bacterial pathogen contagion probability of 0.1 or less. Therefore, we defined that this probability is 0.15 in the simulations.

**Table 4.** Simultaneous extinction of all pathogenic bacterial species according to the bacterial pathogen contagion probability.

| Bacterial pathogen contagion probability | Number of times that all pathogenic bacterial species disappeared (in 2 000 possibilities) |
| --- | --- |
| 0.05 | 570 |
| 0.1 | 70 |
| 0.15 | 2 |

### 4.2 Calibration of the contagion probability

As previously explained, individuals take antibiotics whenever pathogenic bacteria infect them. However, antibiotics remove other bacteria present in the microbiome carrying antibiotic-resistance and virulence genes, beyond pathogenic bacteria. Therefore, it is essential to calibrate the probability of passing these genes by avoiding their population’s loss. These genes disappeared from the community when the number of eliminated genes was higher than the number of genes passed between individuals.

To better understand the impact of the gene contagion probability parameter, we then studied the simpler case: only antibiotics can eliminate genes, and there is no fitness cost for harboring resistance genes (hence, loss rate = 0).

As we can see in Fig. 2, when the gene contagion probability was less than 0.005 (Figs 2A and 2B), virulence genes disappeared from the population. On the other hand, when the contagion probability of genes was higher than 0.01 (Figs 2E and 2F), several individuals had the maximum diversity of genes in their metagenome, which does not correspond to the observation in (Escudeiro et al., 2019). Following our results, we assumed that the gene contagion probability must be 0.005 or 0.01 (Figs 2C and 2D).

**Figure 2.**
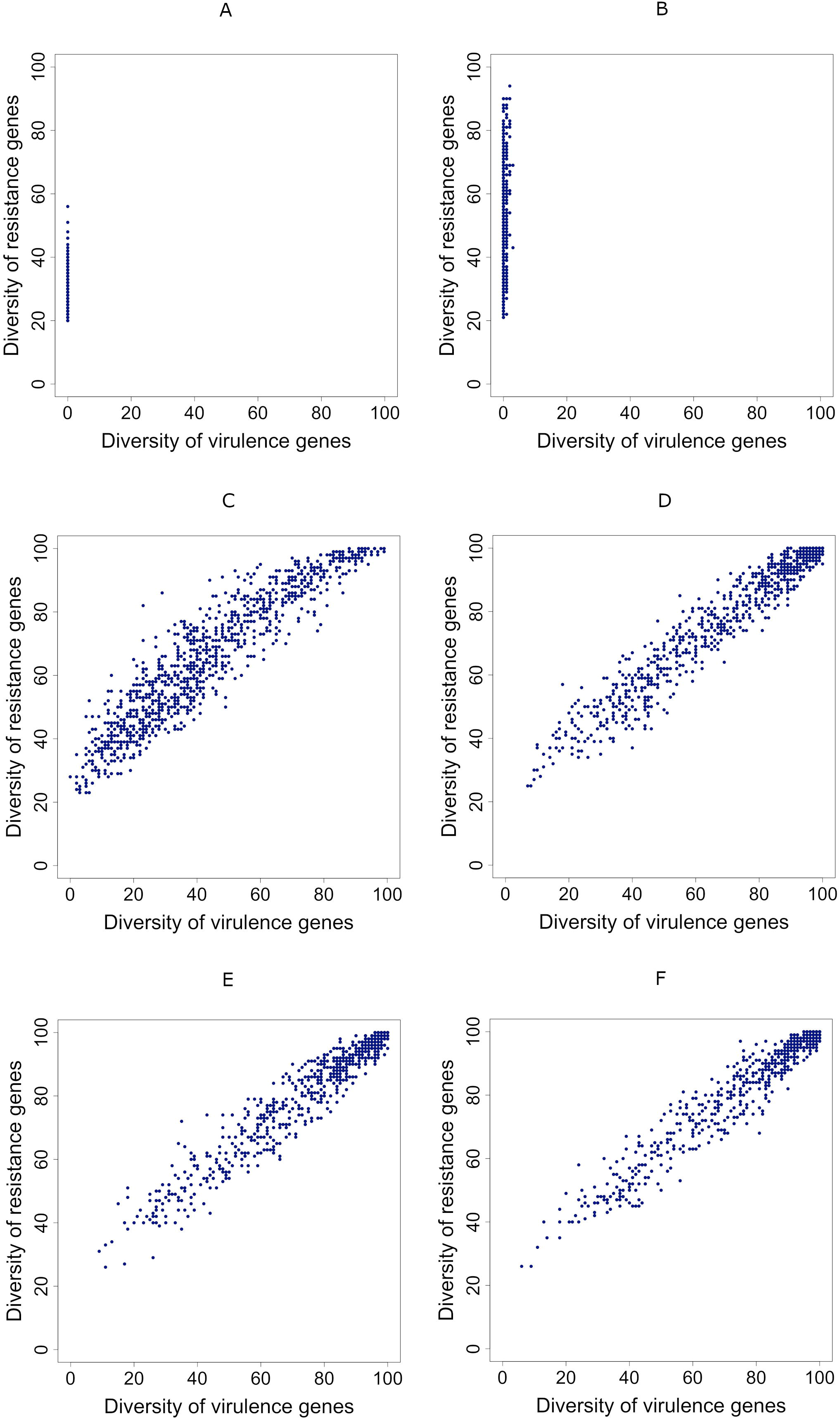
Effect of the gene transmission probability. A to F: the relationship between the diversity of resistance genes (vertical axes) and the diversity of virulence genes (horizontal axes). Each dot represents the case of an individual metagenome. A and B: disappearance of the diversity of virulence genes; C and D: positive correlation between the diversity of resistance genes and the diversity of virulence genes; E and F: positive correlation between the diversity of resistance genes and the diversity of virulence, with many individuals having a high diversity of the two gene types. Parameters as follows. In all cases, we set resistance genes loss rate = 0. In A, when the gene contagion probability is low (0.0005), virulence genes disappeared from the network. In B, gene contagion probability = 0.0025 (R = 0.309, slope = 11.00, p-value = 1.47×10^−23^); In C, gene contagion probability = 0.005 (R = 0.934, slope = 0.798, p-value = ~0); In D, gene contagion probability = 0.01 (R = 0.973, slope = 0.757, p-value = ~0); In E, gene contagion probability = 0.015 (R = 0.972, slope = 0.754, p-value = ~0); In F, gene contagion probability = 0.02 (R = 0.976, slope = 0.751, p-value = ~0).

### 4.3 Correlation between diversities is positive if gene contagion probability is higher than the resistance gene loss rate

We studied the correlation between virulence genes’ diversity and the diversity of resistance genes effect for different combinations of gene contagion probability and resistance gene loss rate. For that, we fixed all the other parameters (see Table 2). Fig. 3 shows that if the gene contagion probability is higher, the same or only slightly lower than the loss rate, the correlation between the diversity of virulence genes and the diversity of resistance genes is positive (Supp. Table 1, Fig. 3).

**Figure 3.**
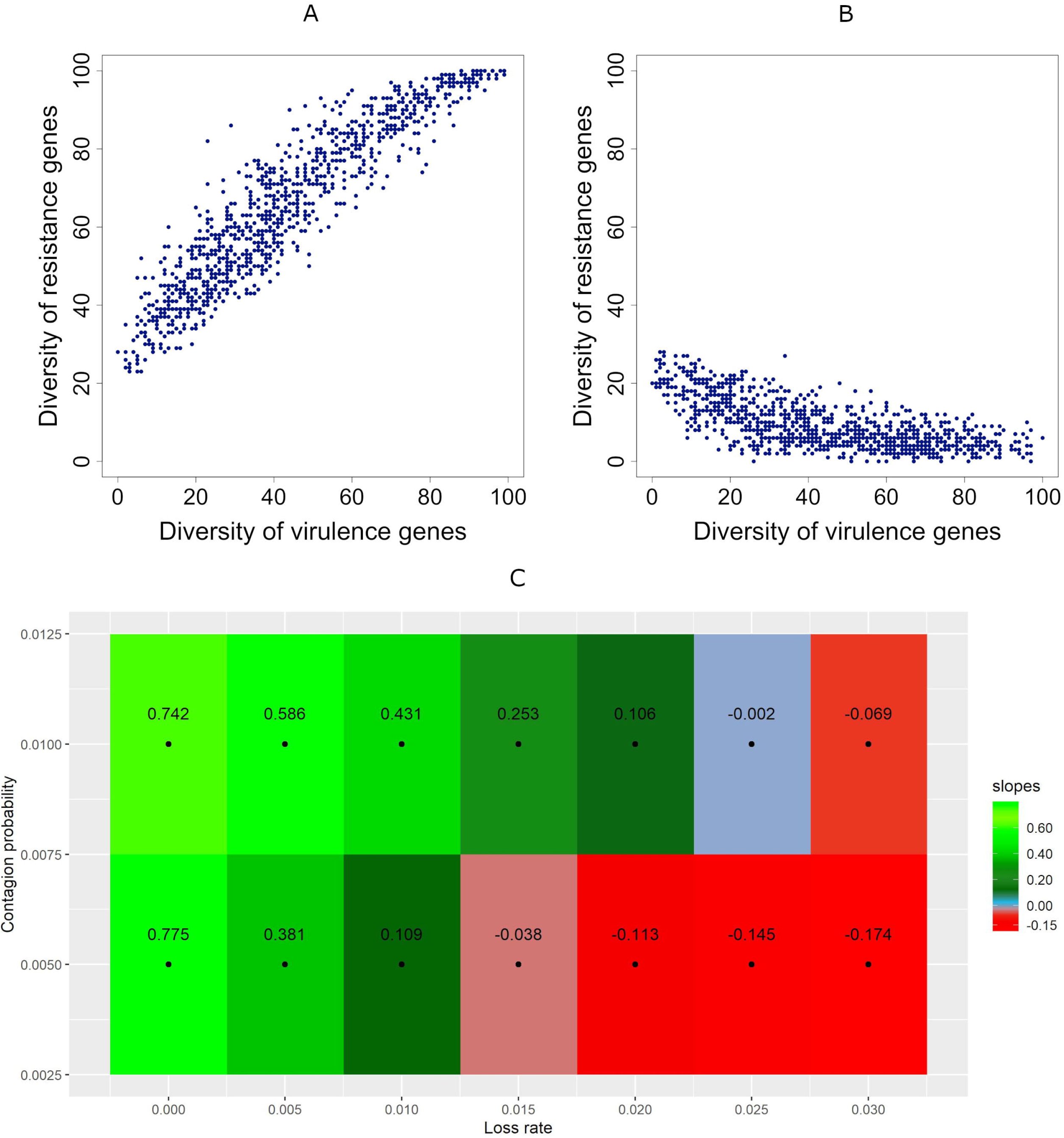
Effect of the relative values of the gene contagion probability and the resistance genes loss rate. A and B: the relationship between the diversity of virulence genes (horizontal axes) and the diversity of resistance genes (vertical axes). Each dot represents the case of an individual metagenome. In both A and B, the gene contagion probability = 0.005. A: resistance genes loss rate = 0, which is lower than the gene contagion probability, resulting in a positive slope; (R = 0.929, slope = 0.775, p-value ~ 0). B: resistance genes loss rate = 0.03, which higher than the gene contagion probability, resulting in a negative slope; (R = −0.682, slope = −0.174, p-value = 1.19×10^−137^). C: Slope of the regression between the diversity of virulence and resistance genes according to the loss rate (horizontal axes) and the gene contagion probability (vertical axes). Green: positive slopes; Red: negative slopes; Blue: the slope is not significantly different from zero (p-value ≥ 1×10^−6^).

### 4.4 Correlations maintain signal even when people take antibiotics randomly

Until now, we have studied the correlations when people take antibiotics because bacterial pathogens infected them through their contacts in the network. Here we examine what happens if individuals take antibiotics at random, not because pathogens infected them. We chose these individuals randomly from the population in each cycle. In the previous simulations, there were 13/1000 individuals, on average, taking antibiotics in each cycle. Thus, in this section, we considered that the probability that a random individual takes antibiotics is 0.013. At the end of simulations, we obtained the same correlations’ signals when assuming that people take antibiotics randomly or because pathogens infected them through their contacts in the network (compare Supp. Table 1 and Fig. 3C with Supp. Table 2.1 and Supp. Fig. 2.1 respectively). In other words, whatever are the reasons for taking antibiotics, the correlation between diversities is positive if gene contagion probability is higher than the resistance gene loss rate.

### 4.5 Taking antibiotics is crucial for a positive correlation between virulence and resistance genes’ diversity

In the previous sections, we showed that if the gene contagion probability is higher than the loss rate, the outcome is a positive correlation between virulence and resistance genes’ diversity. Here we show that taking antibiotics is crucial for this outcome (Supp. Fig. 3.1).

If no one takes antibiotics, there is no counter-selective pressure on commensal bacteria encoding virulence genes. The result is that virulence genes’ diversity gets the maximum possible value in everyone’s metagenome in the community (in Supp. Fig. 3.1 A, B and C, all the dots converge to the right). If the loss rate is null (if there is no fitness cost of resistance), all metagenomes accumulate every possible virulence and resistance gene families, so their diversity attains the maximum achievable value (in Supp. Fig. 3.1 A, all the dots congregate to a single point at the top right corner). If the loss rate is low, there is some diversity of resistance genes in the population (in Supp. Fig. 3.1 B, all the dots distribute in a vertical line on the right side). Finally, if the loss rate is high, more resistance genes are lost than those that accumulate through contagion, so all metagenomes lose all virulence genes (see Suppl. Fig 3.1 C, where all the dots congregate to a single point at the bottom right corner).

### 4.6 Positive correlations are robust under changes in the main simulated system’s properties

We have seen that the positive correlation between virulence and resistance genes’ diversity is positive if the gene contagion probability is higher than the loss rate (Fig. 3C). We then analyzed the robustness of this result. The next five subsections show the impact of changing the simulations’ parameters or changing the network itself. We studied the following parameters: population size, the ratio between virulence genes and antibiotic resistance genes, the elimination probability under antibiotic intake, the proportion of the population harboring antibiotic-resistance genes in their metagenome, and the network type.

#### 4.6.1 Population size has no impact on the correlations’ signal

Due to computer power constraints, we had to assume that the population has just a thousand people. Therefore, it is essential to understand whether population size impacts the correlations’ signals. We performed simulations with a population size of 3000 individuals, instead of 1000 individuals, for the 14 conditions shown in Fig. 3C. Although there were significant differences between the slopes in three cases, we didn’t observe a change of the correlation’s signal from the cases where the population size was 1000 individuals (Supp. Table 4.1 and Supp. Fig. 4.1). An increase in the population size leads to a rise in the number of intermediaries between two distant individuals. Therefore, for virulence genes and antibiotic resistance genes to be transferred between these two faraway individuals, more contacts are needed and, consequently, more time is required to achieve a stable correlation.

#### 4.6.2 The ratios between virulence and antibiotic resistance genes diversities have no impact on correlations’ signal

In all the other sections, we considered that virulence and resistance genes have the same total diversity. Here, we studied the effect of assuming that the diversity of virulence genes is different from that of resistance genes for the same 14 conditions of gene contagion probability and loss rate studied in the previous section. For that, we performed simulations similar to the previous ones, but with the following ratios between virulence and antibiotic resistance genes: 1:2, 1:4, 2:1, 4:1. Although there were significant differences between the slopes in 48 out of 56 cases, we didn’t observe a change of the correlation’s signal (Supp. Tables 5.1 to 5.4 and Supp. Figs 5.1 to 5.4).

#### 4.6.3 The correlation’s signal is robust under changes in the gene elimination probability when people take antibiotics

When an individual takes an antibiotic, virulence genes and resistance genes are eliminated from the metagenome with a probability of 0.7 (except for resistance genes corresponding to the antibiotic used, which are selected, not eliminated). In this section, we analyzed the impact of using other elimination probabilities when an individual takes an antibiotic. For that, we performed simulations similar to the previous ones, for the same 14 conditions of gene contagion probability and loss rate, but where the probability of eliminating genes under antibiotic intake is 0.3 and 0.5 for all gene types (instead of 0.7). In 19 out of 28 cases, the slopes were not significantly different from those obtained with a probability of 0.7 (Supp. Tables 6.1 to 6.2). The slopes were different in the other nine cases, but the signal remained the same (Supp. Tables 6.1 to 6.2 and Supp. Figs 6.1 and 6.2).

We also checked the impact of setting the probability of eliminating antibiotic resistance genes different from that of eliminating virulence genes. Although the slopes were significantly different in 51 out of 84 tested cases, the slopes’ signal remained the same (Supp. Tables 7.1 to 7.6 and Supp. Figs 7.1 to 7.6). Overall, these results show that the slope’s signal is robust under changes in the elimination probability.

#### 4.6.4 The initial proportion of metagenomes containing antibiotic-resistance genes has no impact on correlations’ signal

In the previous sections, we considered that every individual carries all the antibiotic resistance genes in two alternative states at the beginning of the simulation. Either resistance genes were present at low copy numbers (hence being unable to be transmitted to other people) or at high copy numbers because they previously selected by antibiotic exposure (thus transmitting to other people). In this section, we study the effect of considering that, initially, only 10% of the metagenomes contain antibiotic-resistance genes. With this parameter changed, the simulations take more time to stabilize because 90% of the population receives resistance genes only through contagion. We performed simulations similar to the ones shown in Figure 3, but with 5000 cycles. The final slopes are not significantly different from the case where all metagenomes initially contain antibiotic-resistance genes (Supp. Table 8.1 and Supp. Fig 8.1).

#### 4.6.5 The network type has no impact on correlations’ signal

The simulations leading to Fig. 3 were performed in a network with a rewiring probability of *p* = 0.5 (see Methods). We then performed similar simulations but in a regular (p = 0) and in a random (p = 1) networks. This parameter did not change the correlation signals (see Suppl. Tables 9.1 and 9.2). However, the time needed (number of cycles) to reach a stable distribution was lower for higher values of p (Supp. Fig. 9.3)

This section 3.5 shows that the simulated system’s main parameters have no impact on the correlation’s signal between the virulence and resistance genes diversities.

## 5. Discussion

Antibiotics affect hundreds of commensal and mutualist bacterial strains and species, even if their target is bacterial pathogens. Moreover, healthy animals often take antibiotics, given the properties of these drugs as growth-promoters. With these two processes, antibiotic-sensitive bacteria are counter-selected, raising the frequency of antibiotic resistance cells in metagenomes. Meanwhile, metagenomes, both from sick and healthy people, harbor virulence genes. This paper aimed to understand why there is a positive correlation between the diversity of virulence and antibiotic-resistance genes among human populations’ microbiomes (Escudeiro et al., 2019).

Our simulations’ main result is that a positive correlation emerges if the contagion probability is higher than the loss rate of antibiotic-resistance genes. We can understand this result in the following way.

In the absence of infection by bacterial pathogens, people do not take antibiotics (in that particular cycle), and thus, the diversity of virulence genes increase through contagion with other people. However, two opposing forces play a role in resistance genes of the microbiomes of people not taking antibiotics. Contagion from other people in the network makes the diversity of resistance genes to increase, whereas gene loss decreases it. At the end of a cycle, the diversity of resistance genes increases exclusively if the effect of contagion is higher than that of gene loss. The gene loss is just the consequence of the fitness cost imposed by resistance determinants (chromosomal mutations or genes) in competition with susceptible cells. However, the contagion effect has two main contributors: the contagion probability and the number of connections (which depends on the network type and varies from person to person in non-regular networks). Figs. 3C and the corresponding figures in Supplementary File (Suppl. Figs. 4.1, 5.1 – 5.4, 6.1, 6.2, 7.1 – 7.6, 8.1, 9.1 and 9.2) show that if the contagion rate is higher than the loss rate, a positive correlation emerges between the diversity of antibiotic resistance genes and virulence genes.

At first, one might expect to see a negative correlation whenever the contagion probability is lower than the loss rate, but that is not always the case. Indeed, when the contagion probability is only slightly lower than the loss rate, the correlation is positive.

For example, if the contagion probability is 0.005 and the loss rate is 0.01, the correlation is still positive (Fig. 3A and 3C, Supp. Table 1). The reason for these counter-intuitive cases is that, in each cycle, one individual contacts with four other individuals, and during each of these contacts they share bacteria from its microbiomes. In turn, each individual can only be medicated with antibiotics once (at the end of a cycle). That implies that the rate of loss of resistance genes applies only once in a cycle. Therefore, the impact of the contagion rate is higher than the loss rate of resistance genes.

Importantly, our conclusion that a positive correlation emerges if the contagion probability is higher than the loss rate of antibiotic-resistance genes is robust under changes of the population size (Supp. Tables 4.1), the ratio between virulence and antibiotic resistance genes (Supp. Tables 5.1 to 5.4), the elimination probability under antibiotic intake (Supp. Tables 6.1 to 7.6), or the network type (Supp. Tables 9.1 and 9.2).

We assumed that, by default, resistance determinants are already present in little amounts in all metagenomes because they are a part of the natural bacterial lifestyle, and human beings have used massive quantities of antibiotics since the 1940s. What is the impact of this assumption? As shown in Supp. Tables 8.1, if we assumed that, initially, only 10% of the metagenomes contain antibiotic-resistance genes, the final correlations between the diversity of resistance genes and the diversity of virulence genes are the same as in the default case. The only difference is that more cycles are needed to stabilize the correlation.

The contagion probability between people and the loss rate of antibiotic-resistance genes are the two critical parameters of our main result, so it is relevant to know their actual values. Human microbiomes’ interest strongly increased in recent years, yet we still do not know how much is the contagion probability of non-housekeeping genes. For example, we know that human microbiomes are more similar among humans living together, irrespective of the genetic relatedness, suggesting that transmission is a critical factor of the microbiome constitution (Rothschild et al., 2018).

Sarowska and colleagues recently reviewed the fate of the so-called extraintestinal pathogenic *Escherichia coli* (ExPEC), which are facultative pathogens of the normal human intestinal microbiome. ExPEC pathogenicity relies on many virulence genes, and pathogenicity islands, or mobile genetic elements (such as plasmids) encoding some of them. One of the authors’ conclusions is precisely the difficulty in assigning ExPEC transmission to people due to the delay between ExPEC colonization and infection: ExPEC cell can live in human intestines for months or even years before starting an infection (Sarowska et al., 2019). The same problem applies to the transmission rate of antibiotic-resistance genes: there is very little data on transmission rates between people (Andersson and Hughes, 2017).

We have seen that the relationship between the contagion rate and loss rate is paramount to understand the positive correlation between resistance and virulence genes diversity. So, we now discuss how much is the loss rate of resistance determinants in human metagenomes. Several longitudinal studies have shown that antibiotic-resistance genes often remain tens of days, sometimes months, in human gut microbiomes (Horcajada et al., 2002; Lautenbach et al., 2006; O’Fallon et al., 2009; Rogers et al., 2012). While still harboring resistance genes, people most probably contact with several other people. Yet, it is still unclear what is the relationship between contagion and loss rates.

As explained in the methods section, the loss of antibiotic resistance results from the fitness cost of resistance determinants on bacterial cells (compared to otherwise isogenic susceptible cells). Several studies have shown that resistance determinants, here broadly comprising resistance mutations and resistance genes encoded in the chromosome or plasmids, impose a fitness cost on their hosts (giving the sensitive strains a growth advantage) (Andersson and Levin, 1999). However, several mechanisms decrease or even eliminate it. First, compensatory mutations, which mask the deleterious effects of resistance mutations, have been observed in several studies (Levin et al., 1997; Schrag et al., 1997; Bjorkman et al., 2000; Maisnier-Patin and Andersson, 2004; Nilsson et al., 2006). Second, resistance mutations can even be beneficial in specific resistance genetic backgrounds(Trindade et al., 2009). Third, while resistance plasmids often impose a fitness cost to their hosts, it has also been observed that plasmid and/or cells need just a few hundreds of bacterial generations to adapt to each other (Bouma and Lenski, 1988; Modi and Adams, 1991; Dahlberg and Chao, 2003; Dionisio et al., 2005; Harrison et al., 2015). Fourth, plasmids sometimes increase the fitness of bacteria that already harbor a resistance mutation (Silva et al., 2011); likewise, some resistance mutations increase the fitness of plasmid bearing cells (Silva et al., 2011). The same may happen with two plasmids: one of them compensating for the fitness-cost of the other (Silva et al., 2011; San Millan et al., 2014). Fifth, plasmids may interact with other plasmids to facilitate their transfer (Gama et al., 2017c, 2017a, 2017b, 2018). Sixth, a few works suggested that plasmids appear costly because their fitness effect is often measured a long time after its isolation from nature (Lau et al., 2013; Gama et al., 2018).

Together, these six factors suggest that the fitness cost of resistance determinants is often very low or null, allowing the permanence of resistance determinants in microbiomes for long periods. This stability of resistance determinants implies that their loss rate, the probability that a metagenome loses a particular resistance gene or mutation, is undoubtedly lower than the contagion probability. Therefore, antibiotic consumption and contagion between people lead to a positive correlation between the diversity of resistance genes and virulence genes.

## 6. Concluding remarks

The simple fact that people contaminate between themselves, and antibiotic use, is chief to explain the positive correlation between antibiotic resistance gene diversity and virulence gene diversity across human metagenomes. This result is robust and general because we made very few assumptions. This result also has worrying health implications: people with a higher diversity of resistance genes in their metagenomes have a higher diversity of virulence genes. Such co-presence may potentiate the appearance of plasmids or bacteria encoding virulence and resistance genes simultaneously. Meanwhile, the current restrictive measures due to the COVID-19 pandemic may weaken this correlation between the diversity of resistance genes and antibiotics and virulence factors due to a decrease in the contagion rate (Domingues et al., 2020).

## 7. Conflict of Interest

The authors declare that the research was conducted in the absence of any commercial or financial relationships that could be construed as a potential conflict of interest.

## 8. Author Contributions

CD, JR, TN, and FD conceived the study and designed the simulations. CD and JR wrote the computer program; CD, JR, TN, JP, and FD analyzed the data. CD, JR, TN, and FD wrote the first draft of the manuscript, with contributions of JP and FM. All authors contributed to manuscript revision, read, and approved the submitted version.

## 9. Funding

Teresa Nogueira was supported by contract ALG-01-0145-FEDER-028824 and Francisca Monteiro acknowledges FCT-Fundação para a Ciência e a Tecnologia, I.P., for her Postdoc fellowship grant SFRH/BPD/123504/2016. FCT supports cE3c by contract UIDP/00329/2020.

## 10. Acknowledgments

This manuscript has been released as a pre-print at BioRxiv, (Domingues et al.).

## Notes

### Competing Interest Statement

The authors have declared no competing interest.

### Summary of Updates

We changed what triggers antibiotic administration.

## References

Andersson, D. I., and Hughes, D. (2017). Selection and Transmission of Antibiotic-Resistant Bacteria. Microbiol Spectr 5. doi:10.1128/microbiolspec.MTBP-0013-2016.

Andersson, D. I., and Levin, B. R. (1999). The biological cost of antibiotic resistance. Current opinion in microbiology 2, 489–493. doi:10.1016/S1369-5274(99)00005-3.

Bjorkman, J., Nagaev, I., Berg, O. G., Hughes, D., and Andersson, D. I. (2000). Effects of environment on compensatory mutations to ameliorate costs of antibiotic resistance. Science 287, 1479–1482. doi:10.1126/science.287.5457.1479.

Blaser, M. J., and Falkow, S. (2009). What are the consequences of the disappearing human microbiota? Nature reviews. Microbiology 7, 887–894. doi:10.1038/nrmicro2245.

Bouma, J. E., and Lenski, R. E. (1988). Evolution of a Bacteria Plasmid Association. Nature 335, 351–352. doi:10.1038/335351a0.

Cassini, A., Hogberg, L. D., Plachouras, D., Quattrocchi, A., Hoxha, A., Simonsen, G. S., et al. (2019). Attributable deaths and disability-adjusted life-years caused by infections with antibiotic-resistant bacteria in the EU and the European Economic Area in 2015: a population-level modelling analysis. The Lancet. Infectious diseases 19, 56–66. doi:10.1016/S1473-3099(18)30605-4.

Castanon, J. I. R. (2007). History of the use of antibiotic as growth promoters in European poultry feeds. Poultry Sci 86, 2466–2471. doi:10.3382/ps.2007-00249.

Dahlberg, C., and Chao, L. (2003). Amelioration of the cost of conjugative plasmid carriage in Eschericha coli K12. Genetics 165, 1641–1649.

Dionisio, F., Conceicao, I. C., Marques, A. C., Fernandes, L., and Gordo, I. (2005). The evolution of a conjugative plasmid and its ability to increase bacterial fitness. Biology Letters 1, 250–2. doi:10.1098/rsbl.2004.0275.

Domingues, C. P. F., Rebelo, J. S., Dionisio, F., Botelho, A., and Nogueira, T. (2020). The Social Distancing Imposed To Contain COVID-19 Can Affect Our Microbiome: a Double-Edged Sword in Human Health. mSphere 5, e00716–20, /msphere/5/5/mSphere716-20.atom. doi:10.1128/mSphere.00716-20.

Escudeiro, P., Pothier, J., Dionisio, F., and Nogueira, T. (2019). Antibiotic Resistance Gene Diversity and Virulence Gene Diversity Are Correlated in Human Gut and Environmental Microbiomes. mSphere 4. doi:10.1128/mSphere.00135-19.

Ferretti, P., Pasolli, E., Tett, A., Asnicar, F., Gorfer, V., Fedi, S., et al. (2018). Mother-to-Infant Microbial Transmission from Different Body Sites Shapes the Developing Infant Gut Microbiome. Cell Host Microbe 24, 133–+.

Gama, J. A., Zilhao, R., and Dionisio, F. (2017a). Conjugation efficiency depends on intra and intercellular interactions between distinct plasmids: Plasmids promote the immigration of other plasmids but repress co-colonizing plasmids. Plasmid 93, 6–16. doi:10.1016/j.plasmid.2017.08.003.

Gama, J. A., Zilhao, R., and Dionisio, F. (2017b). Co-resident plasmids travel together. Plasmid 93, 24–29. doi:10.1016/j.plasmid.2017.08.004.

Gama, J. A., Zilhao, R., and Dionisio, F. (2017c). Multiple plasmid interference - Pledging allegiance to my enemy’s enemy. Plasmid 93, 17–23. doi:10.1016/j.plasmid.2017.08.002.

Gama, J. A., Zilhao, R., and Dionisio, F. (2018). Impact of plasmid interactions with the chromosome and other plasmids on the spread of antibiotic resistance. Plasmid. doi:10.1016/j.plasmid.2018.09.009.

Harrison, E., Guymer, D., Spiers, A. J., Paterson, S., and Brockhurst, M. A. (2015). Parallel Compensatory Evolution Stabilizes Plasmids across the Parasitism-Mutualism Continuum. Current biology◻: CB 25, 2034–2039. doi:10.1016/j.cub.2015.06.024.

Horcajada, J. P., Vila, J., Moreno-Martinez, A., Ruiz, J., Martinez, J. A., Sanchez, M., et al. (2002). Molecular epidemiology and evolution of resistance to quinolones in Escherichia coli after prolonged administration of ciprofloxacin in patients with prostatitis. J Antimicrob Chemoth 49, 55–59. doi:10.1093/jac/49.1.55.

Lau, B. T. C., Malkus, P., and Paulsson, J. (2013). New quantitative methods for measuring plasmid loss rates reveal unexpected stability. Plasmid 70, 353–361. doi:10.1016/j.plasmid.2013.07.007.

Lautenbach, E., Tolomeo, P., Mao, X. Q., Fishman, N. O., Metlay, J. P., Bilker, W. B., et al. (2006). Duration of outpatient fecal colonization due to Escherichia coli isolates with decreased susceptibility to fluoroquinolones: Longitudinal study of patients recently discharged from the hospital. Antimicrobial agents and chemotherapy 50, 3939–3943. doi:10.1128/Aac.00503-06.

Levin, B. R., Lipsitch, M., Perrot, V., Schrag, S., Antia, R., Simonsen, L., et al. (1997). The population genetics of antibiotic resistance. Clinical infectious diseases◻: an official publication of the Infectious Diseases Society of America 24, S9–S16. doi:DOI 10.1093/clinids/24.Supplement_1.S9.

Maisnier-Patin, S., and Andersson, D. I. (2004). Adaptation to the deleterious effects of antimicrobial drug resistance mutations by compensatory evolution. Research in microbiology 155, 360–369. doi:10.1016/j.resmic.2004.01.019.

Modi, R. I., and Adams, J. (1991). Coevolution in Bacterial-Plasmid Populations. Evolution; international journal of organic evolution 45, 656–667. doi:DOI 10.1111/j.1558-5646.1991.tb04336.x.

Moeller, A. H., Foerster, S., Wilson, M. L., Pusey, A. E., Hahn, B. H., and Ochman, H. (2016). Social behavior shapes the chimpanzee pan-microbiome. Sci Adv 2. doi:10.1126/sciadv.1500997.

Nayfach, S., Rodriguez-Mueller, B., Garud, N., and Pollard, K. S. (2016). An integrated metagenomics pipeline for strain profiling reveals novel patterns of bacterial transmission and biogeography. Genome research 26, 1612–1625. doi:10.1101/gr.201863.115.

Nilsson, A. I., Zorzet, A., Kanth, A., Dahlstrom, S., Berg, O. G., and Andersson, D. I. (2006). Reducing the fitness cost of antibiotic resistance by amplification of initiator tRNA genes. Proceedings of the National Academy of Sciences of the United States of America 103, 6976–6981. doi:10.1073/pnas.0602171103.

Nogueira, T., David, P. H. C., and Pothier, J. (2019). Antibiotics as both friends and foes of the human gut microbiome: The microbial community approach. Drug Develop Res 80, 86–97. doi:10.1002/ddr.21466.

O’Fallon, E., Gautam, S., and D’Agata, E. M. C. (2009). Colonization with Multidrug-Resistant Gram-Negative Bacteria: Prolonged Duration and Frequent Cocolonization. Clinical infectious diseases◻: an official publication of the Infectious Diseases Society of America 48, 1375–1381. doi:10.1086/598194.

R Core Team (2015). R: A Language and Environment for Statistical Computing. R Foundation for Statistical Computing, Vienna, Austria.

Rogers, B. A., Kennedy, K. J., Sidjabat, H. E., Jones, M., Collignon, P., and Paterson, D. L. (2012). Prolonged carriage of resistant E-coli by returned travellers: clonality, risk factors and bacterial characteristics. Eur J Clin Microbiol 31, 2413–2420. doi:10.1007/s10096-012-1584-z.

Rothschild, D., Weissbrod, O., Barkan, E., Kurilshikov, A., Korem, T., Zeevi, D., et al. (2018). Environment dominates over host genetics in shaping human gut microbiota. Nature 555, 210–+. doi:10.1038/nature25973.

San Millan, A., Heilbron, K., and MacLean, R. C. (2014). Positive epistasis between co-infecting plasmids promotes plasmid survival in bacterial populations. The ISME journal 8, 601–612. doi:10.1038/ismej.2013.182.

Sarowska, J., Futoma-Koloch, B., Jama-Kmiecik, A., Frej-Madrzak, M., Ksiazczyk, M., Bugla-Ploskonska, G., et al. (2019). Virulence factors, prevalence and potential transmission of extraintestinal pathogenic Escherichia coli isolated from different sources: recent reports. Gut Pathog 11. doi:10.1186/s13099-019-0290-0.

Schrag, S. J., Perrot, V., and Levin, B. R. (1997). Adaptation to the fitness costs of antibiotic resistance in Escherichia coli. P Roy Soc B-Biol Sci 264, 1287–1291. doi:DOI 10.1098/rspb.1997.0178.

Sender, R., Fuchs, S., and Milo, R. (2016). Revised Estimates for the Number of Human and Bacteria Cells in the Body. PLoS biology 14. doi:10.1371/journal.pbio.1002533.

Silva, R. F., Mendonça, S. C., Carvalho, L. M., Reis, A. M., Gordo, I., Trindade, S., et al. (2011). Pervasive sign epistasis between conjugative plasmids and drug-resistance chromosomal mutations. PLoS genetics 7, e1002181.

Trindade, S., Sousa, A., Xavier, K. B., Dionisio, F., Ferreira, M. G., and Gordo, I. (2009). Positive epistasis drives the acquisition of multidrug resistance. PLoS genetics 5, e1000578.

Watts, D. J., and Strogatz, S. H. (1998). Collective dynamics of “small-world” networks. Nature 393, 440–442. doi:DOI 10.1038/30918.

Yassour, M., Jason, E., Hogstrom, L. J., Arthur, T. D., Tripathi, S., Siljander, H., et al. (2018). Strain-Level Analysis of Mother-to-Child Bacterial Transmission during the First Few Months of Life. Cell Host Microbe 24, 146–+. doi:10.1016/j.chom.2018.06.007.

