## Supplemental Information for "The perfect condition for the rising of superbugs: person-to-person contagion and antibiotic use are the key factors responsible for the positive correlation between antibiotic resistance gene diversity and virulence gene diversity in human metagenomes"

***Correspondence:**

Francisco Dionisio

1- **Results of Fig. 3C**

| **Supp. Table 1 –** Parameters used for Fig. 3C and the results of the simulations. | | | |  |
| --- | --- | --- | --- | --- |
| **Contagion probability (%)** | **Resistance genes loss rate (%)** | **Correlation (r) between resistance genes and virulence genes** | **Slope of the regression between resistance genes and virulence genes** | **P-value of the slope** |
| **0.5** | 0 | 0.929 | 0.775 | 0 |
|  | 0.5 | 0.825 | 0.381 | 5.28x10^-250^ |
|  | 1 | 0.414 | 0.109 | 1.24x10^-42^ |
|  | 1.5 | -0.17 | -0.038 | 6.76x10^-8^ |
|  | 2 | -0.472 | -0.113 | 1.29x10^-56^ |
|  | 2.5 | -0.586 | -0.145 | 4.10x10^-93^ |
|  | 3 | -0.682 | -0.174 | 1.19x10^-137^ |
| **1** | 0 | 0.966 | 0.742 | 0 |
|  | 0.5 | 0.94 | 0.586 | 0 |
|  | 1 | 0.848 | 0.431 | 8.35x10^-277^ |
|  | 1.5 | 0.671 | 0.253 | 8.77x10^-132^ |
|  | 2 | 0.362 | 0.106 | 2.95x10^-32^ |
|  | 2.5 | -0.01 | -0.002 | 7.47x10^-1^ |
|  | 3 | -0.319 | -0.069 | 3.69x10^-25^ |

**2 – Correlations maintain signal even when people take antibiotics randomly**

| **Supp. Table 2.1 –** The impact of considering random consumption of antibiotics on the correlation between virulence and resistance genes. | | | | | |
| --- | --- | --- | --- | --- | --- |
| **Contagion probability (%)** | **Resistance genes loss rate (%)** | **Slope considering random antibiotic consumption** | **Slope considering antibiotic consumption due to pathogenic bacteria** | **P-value of T-test for differences of slopes** | **Change in the slope signal?** |
| **0.5** | 0 | 0.762 | 0.775 | 3.41x10^-1^ | No |
|  | 0.5 | 0.423 | 0.381 | 2.31x10^-4^ | No |
|  | 1 | 0.173 | 0.109 | 1.75x10^-8^ | No |
|  | 1.5 | -0.009 | -0.038 | 3.55x10^-3^ | No |
|  | 2 | -0.082 | -0.113 | 1.50x10^-3^ | No |
|  | 2.5 | -0.147 | -0.145 | 8.57x10^-1^ | No |
|  | 3 | -0.149 | -0.174 | 3.06x10^-3^ | No |
| **1** | 0 | 0.742 | 0.742 | 9.82x10^-1^ | No |
|  | 0.5 | 0.588 | 0.586 | 8.97x10^-1^ | No |
|  | 1 | 0.440 | 0.431 | 4.88x10^-1^ | No |
|  | 1.5 | 0.282 | 0.253 | 2.93x10^-2^ | No |
|  | 2 | 0.120 | 0.106 | 2.68x10^-1^ | No |
|  | 2.5 | 0.038 | -0.002 | 4.54x10^-4^ | Yes |
|  | 3 | -0.036 | -0.069 | 8.98x10^-4^ | No |

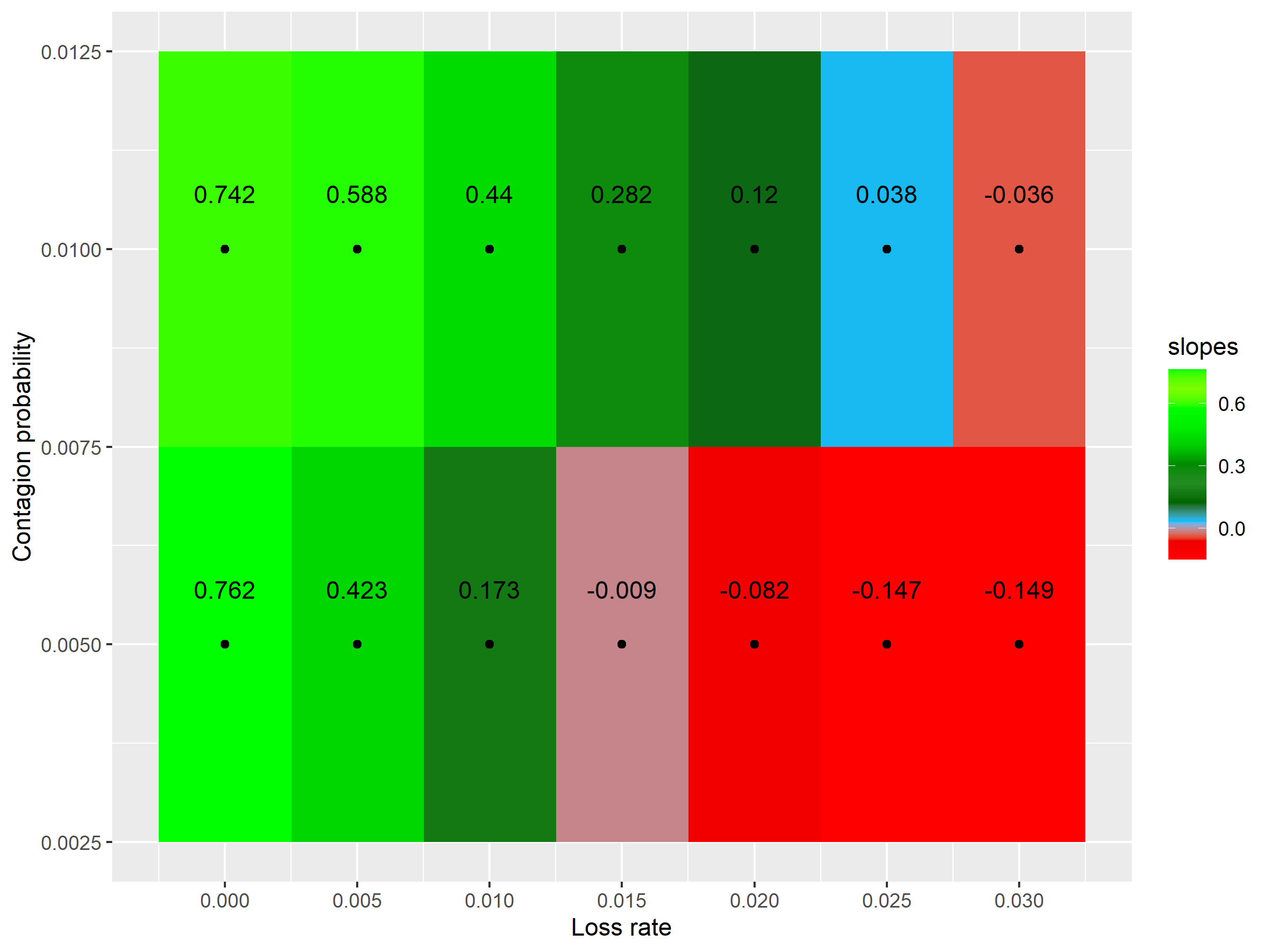

**Supp. Fig. 2.1 - Effect of considering random consumption of antibiotics.** Slope of the regression between the diversity of virulence genes and resistance genes according to the contagion probability (vertical axis) and the loss rate (horizontal axis). Green: positive slopes; and red: negative slopes.

**3 – Taking antibiotics is crucial for a positive correlation between virulence and resistance genes’ diversity**

A) **Resistance genes loss rate = 0**

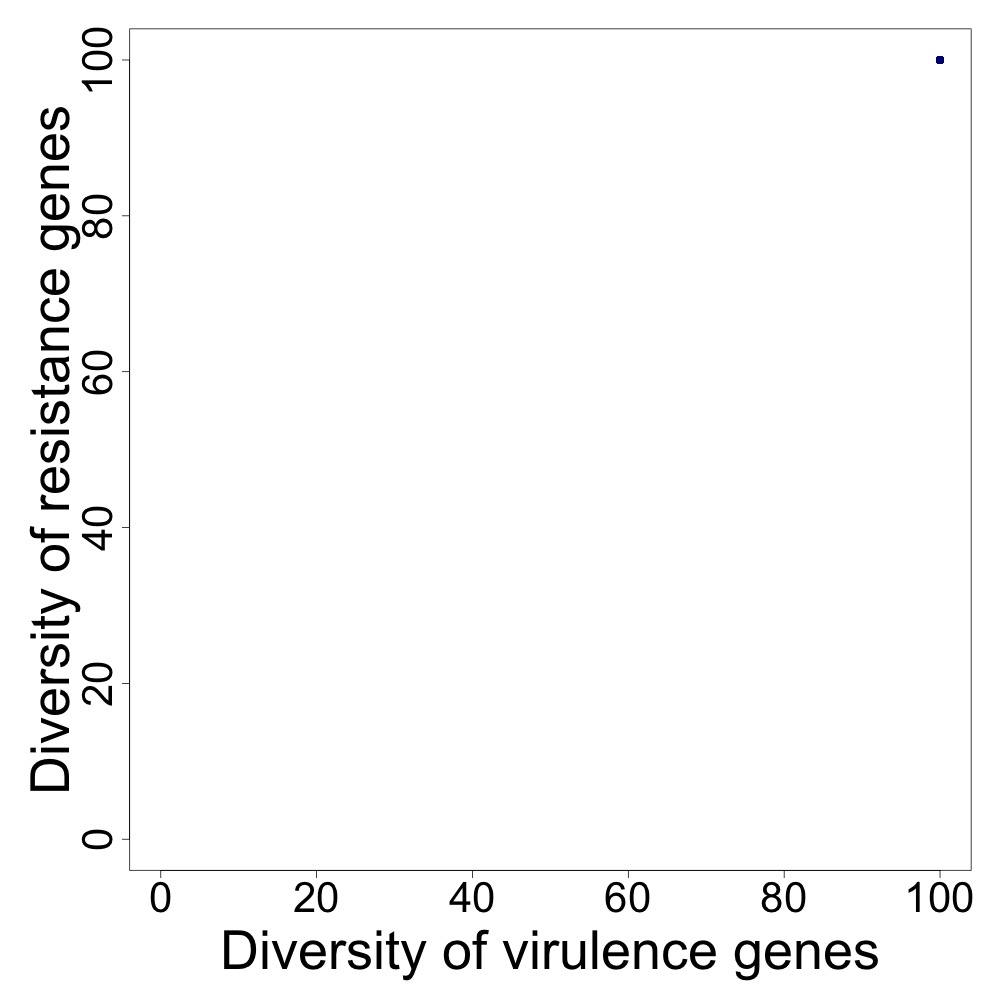

B) **Resistance genes loss rate = 0.005**

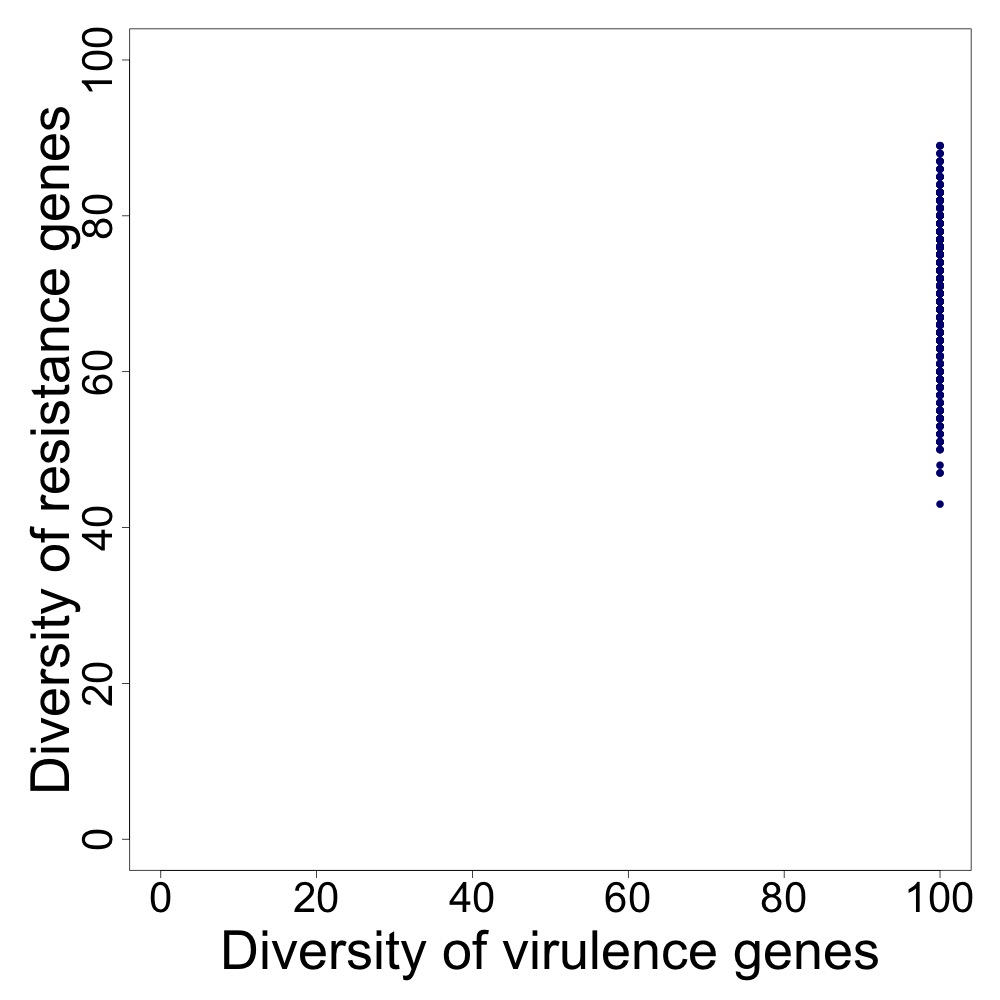

C) **Resistance genes loss rate = 0.03**

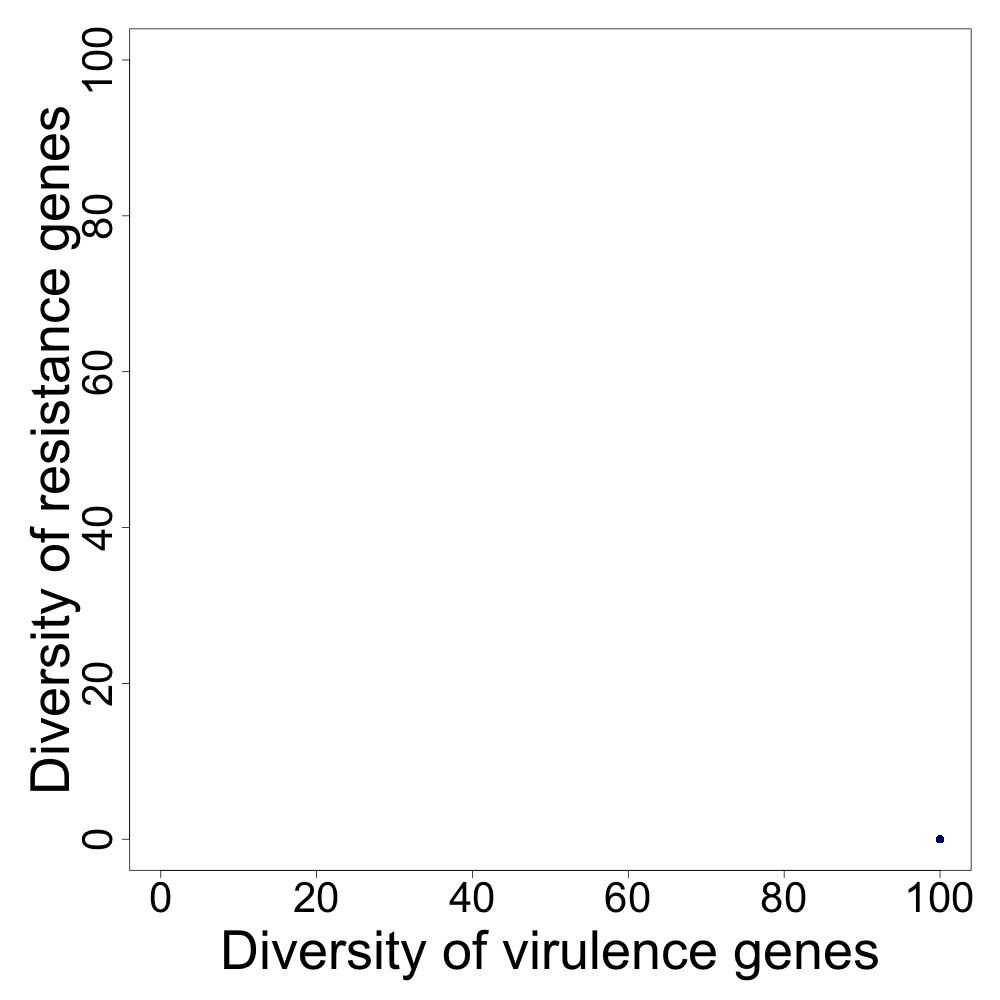

**Figure 3.1 - Effect of considering no consumption of antibiotics.** A to C: the relationship between the diversity of resistance genes (vertical axes) and the diversity of virulence genes (horizontal axes). Each dot represents the case of an individual metagenome. A: accumulation of the diversity of virulence genes and resistance genes. B: . C: disappearence of the diversity of resistance genes; Parameters as follows. In A, B and C the gene contagion probability = 0.005.

**4 - Population size has no impact on the correlation’s signal**

| **Supp. Table 4.1 –** The impact of considering 3000 individuals on the correlation between virulence and resistance genes. | | | | | |
| --- | --- | --- | --- | --- | --- |
| **Contagion probability (%)** | **Resistance genes loss rate (%)** | **Slope considering 3000 individuals** | **Slope considering 1000 individuals** | **P-value of T-test for differences of slopes** | **Change in the slope signal?** |
| **0.5** | 0 | 0.757 | 0.775 | 8.93x10^-2^ | No |
|  | 0.5 | 0.453 | 0.381 | 4.19x10^-13^ | No |
|  | 1 | 0.137 | 0.109 | 4.83x10^-3^ | No |
|  | 1.5 | -0.078 | -0.038 | 1.24x10^-6^ | No |
|  | 2 | -0.147 | -0.113 | 3.13x10^-6^ | No |
|  | 2.5 | -0.166 | -0.145 | 1.97x10^-3^ | No |
|  | 3 | -0.160 | -0.174 | 2.91x10^-2^ | No |
| **1** | 0 | 0.769 | 0.742 | 1.03x10^-4^ | No |
|  | 0.5 | 0.640 | 0.586 | 2.24x10^-9^ | No |
|  | 1 | 0.484 | 0.431 | 1.12x10^-5^ | No |
|  | 1.5 | 0.326 | 0.253 | 2.55x10^-8^ | No |
|  | 2 | 0.152 | 0.106 | 2.07x10^-4^ | No |
|  | 2.5 | 0.007 | -0.002 | 3.53x10^-1^ | Yes |
|  | 3 | -0.106 | -0.069 | 1.35x10^-5^ | No |

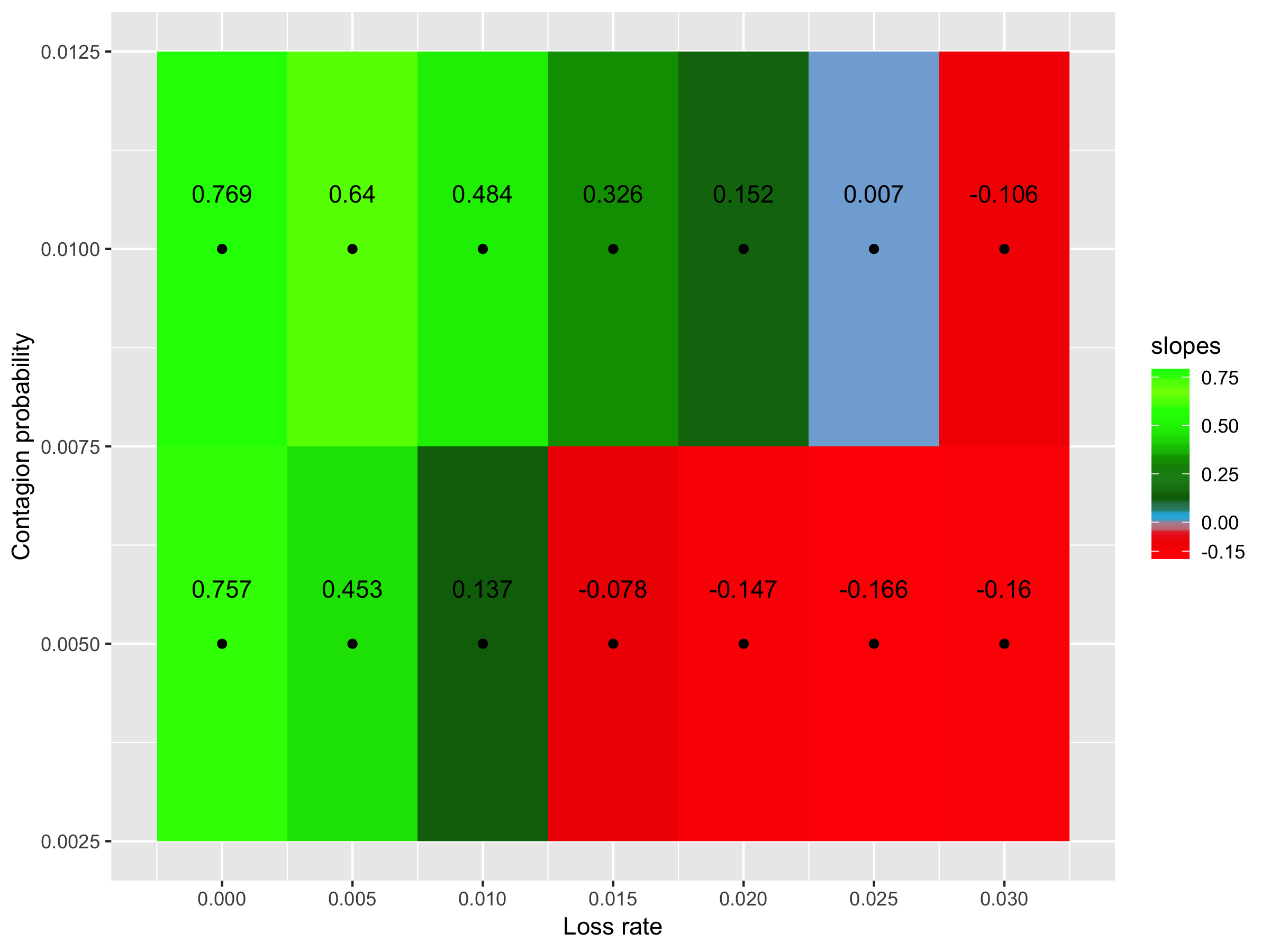

**Supp. Fig. 4.1 - Effect of considering bigger populations (3000 individuals).** Slope of the regression between the diversity of virulence genes and resistance genes according to the contagion probability (vertical axis) and the loss rate (horizontal axis). Green: positive slopes; and red: negative slopes.

**5 - The ratios between virulence and antibiotic resistance genes diversities have no impact on correlation’s signal**

| **Supp. Table 5.1 –** The impact of considering a ratio of 1 virulence gene for 2 resistance genes on the correlation between virulence and resistance genes. | | | | | |
| --- | --- | --- | --- | --- | --- |
| **Contagion probability (%)** | **Resistance genes loss rate (%)** | **Slope considering a ratio of 1:2** | **Slope considering a ratio of 1:1** | **P-value of T-test for differences of slopes** | **Change in the slope signal?** |
| **0.5** | 0 | 1.591 | 0.775 | 1.015073 x10^-285^ | No |
|  | 0.5 | 0.836 | 0.381 | 5.669288 x10^-138^ | No |
|  | 1 | 0.235 | 0.109 | 1.553558 x10^-15^ | No |
|  | 1.5 | -0.094 | -0.038 | 1.068848 x10^-4^ | No |
|  | 2 | -0.213 | -0.113 | 5.657481 x10^-13^ | No |
|  | 2.5 | -0.291 | -0.145 | 3.064856 x10^-29^ | No |
|  | 3 | -0.317 | -0.174 | 1.175099 x10^-28^ | No |
| **1** | 0 | 1.515 | 0.742 | 0 | No |
|  | 0.5 | 1.177 | 0.586 | 6.56x10^-288^ | No |
|  | 1 | 0.862 | 0.431 | 6.60x10^-123^ | No |
|  | 1.5 | 0.548 | 0.253 | 6.89x10^-51^ | No |
|  | 2 | 0.257 | 0.106 | 3.33x10^-17^ | No |
|  | 2.5 | 0.039 | -0.002 | 6.35x10^-3^ | Yes |
|  | 3 | -0.128 | -0.069 | 4.25x10^-5^ | No |

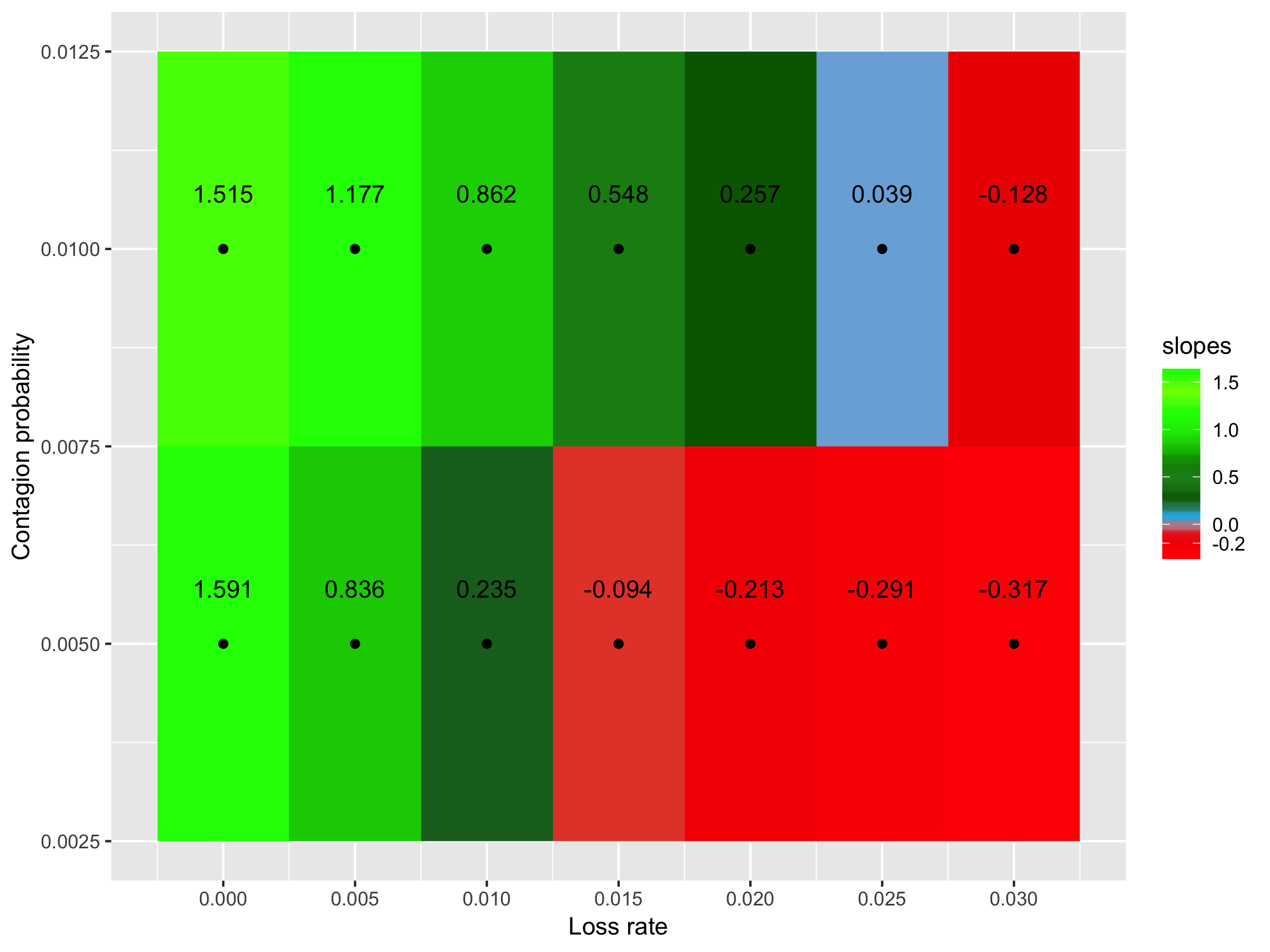

**Supp. Fig. 5.1 - Effect of considering a ratio of 1 virulence gene for 2 resistance genes.** Slope of the regression between the diversity of virulence genes and resistance genes according to the contagion probability (vertical axis) and the loss rate (horizontal axis). Green: positive slopes; and red: negative slopes.

| **Supp. Table 5.2 –** The impact of considering a ratio of 1 virulence gene for 4 resistance genes on the correlation between virulence and resistance genes. | | | | | |
| --- | --- | --- | --- | --- | --- |
| **Contagion probability (%)** | **Resistance genes loss rate (%)** | **Slope considering a ratio of 1:4** | **Slope considering a ratio of 1:1** | **P-value of T-test for differences of slopes** | **Change in the slope signal?** |
| **0.5** | 0 | 3.115 | 0.775 | 0 | No |
|  | 0.5 | 1.592 | 0.381 | 3.68x10^-275^ | No |
|  | 1 | 0.530 | 0.109 | 3.36x10^-50^ | No |
|  | 1.5 | -0.165 | -0.038 | 6.52x10^-8^ | No |
|  | 2 | -0.484 | -0.113 | 7.66x10^-56^ | No |
|  | 2.5 | -0.571 | -0.145 | 1.13x10^-76^ | No |
|  | 3 | -0.665 | -0.174 | 2.05x10^-92^ | No |
| **1** | 0 | 3.012 | 0.742 | 0 | No |
|  | 0.5 | 2.376 | 0.586 | 0 | No |
|  | 1 | 1.684 | 0.431 | 1.25x10^-269^ | No |
|  | 1.5 | 1.086 | 0.253 | 9.29x10^-128^ | No |
|  | 2 | 0.454 | 0.106 | 2.54x10^-27^ | No |
|  | 2.5 | 0.088 | -0.002 | 1.02x10^-3^ | Yes |
|  | 3 | -0.255 | -0.069 | 4.10x10^-18^ | No |

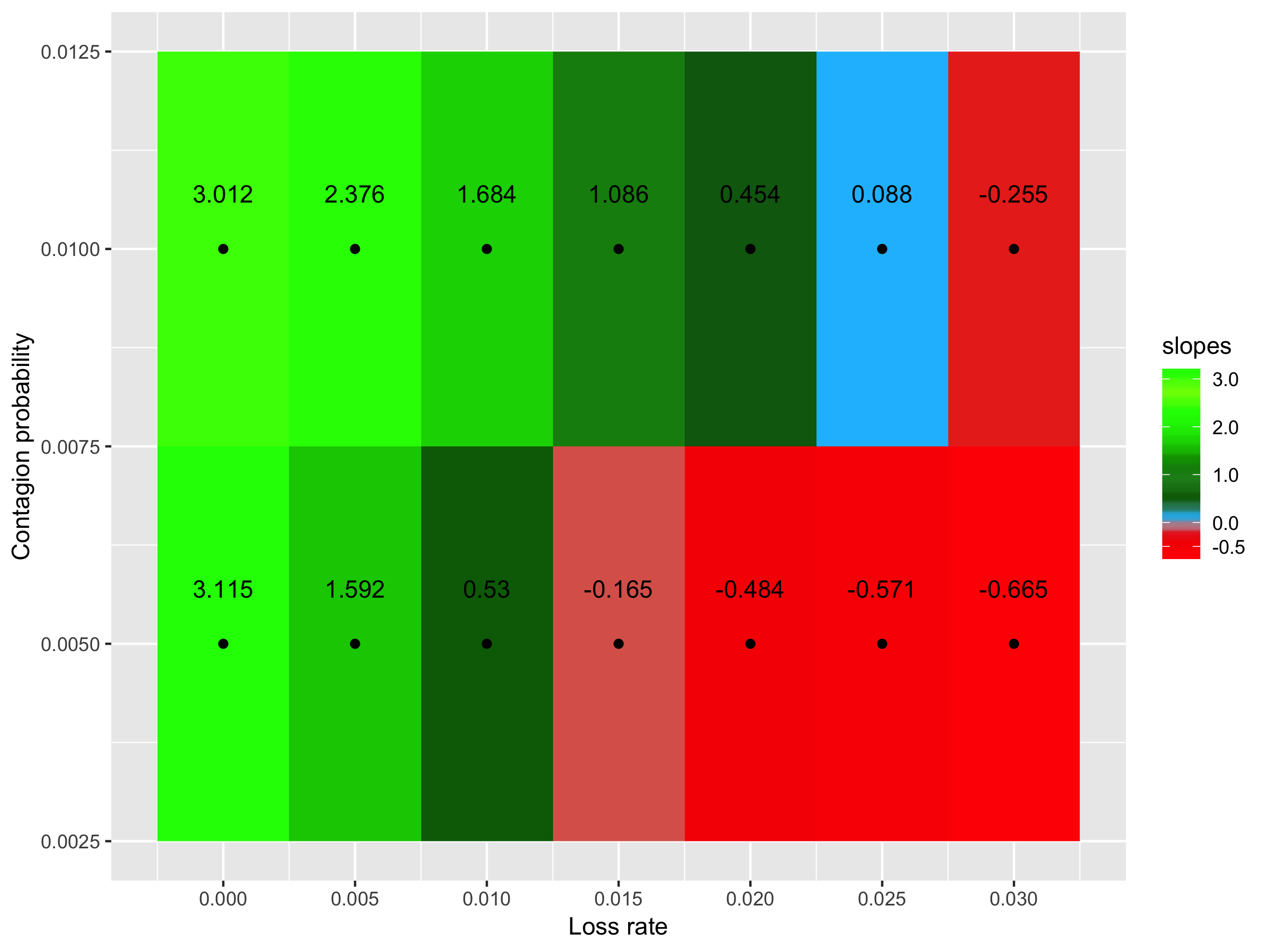

**Supp. Fig. 5.2 - Effect of considering a ratio of 1 virulence gene for 4 resistance genes.** Slope of the regression between the diversity of virulence genes and resistance genes according to the contagion probability (vertical axis) and the loss rate (horizontal axis). Green: positive slopes; and red: negative slopes.

| **Supp. Table 5.3 –** The impact of considering a ratio of 2 virulence genes for 1 resistance gene on the correlation between virulence and resistance genes. | | | | | |
| --- | --- | --- | --- | --- | --- |
| **Contagion probability (%)** | **Resistance genes loss rate (%)** | **Slope considering a ratio of 2:1** | **Slope considering a ratio of 1:1** | **P-value of T-test for differences of slopes** | **Change in the slope signal?** |
| **0.5** | 0 | 0.402 | 0.775 | 2.05x10^-208^ | No |
|  | 0.5 | 0.214 | 0.381 | 7.16x10^-70^ | No |
|  | 1 | 0.061 | 0.109 | 9.67x10^-9^ | No |
|  | 1.5 | -0.020 | -0.038 | 2.08x10^-2^ | No |
|  | 2 | -0.055 | -0.113 | 1.43x10^-14^ | No |
|  | 2.5 | -0.070 | -0.145 | 1.61x10^-25^ | No |
|  | 3 | -0.084 | -0.174 | 2.24x10^-40^ | No |
| **1** | 0 | 0.374 | 0.742 | 0 | No |
|  | 0.5 | 0.298 | 0.586 | 1.19x10^-239^ | No |
|  | 1 | 0.213 | 0.431 | 4.32x10^-104^ | No |
|  | 1.5 | 0.131 | 0.253 | 2.02x10^-33^ | No |
|  | 2 | 0.058 | 0.106 | 4.21x10^-7^ | No |
|  | 2.5 | 0.011 | -0.002 | 1.16x10^-1^ | Yes |
|  | 3 | -0.031 | -0.069 | 2.17x10^-7^ | No |

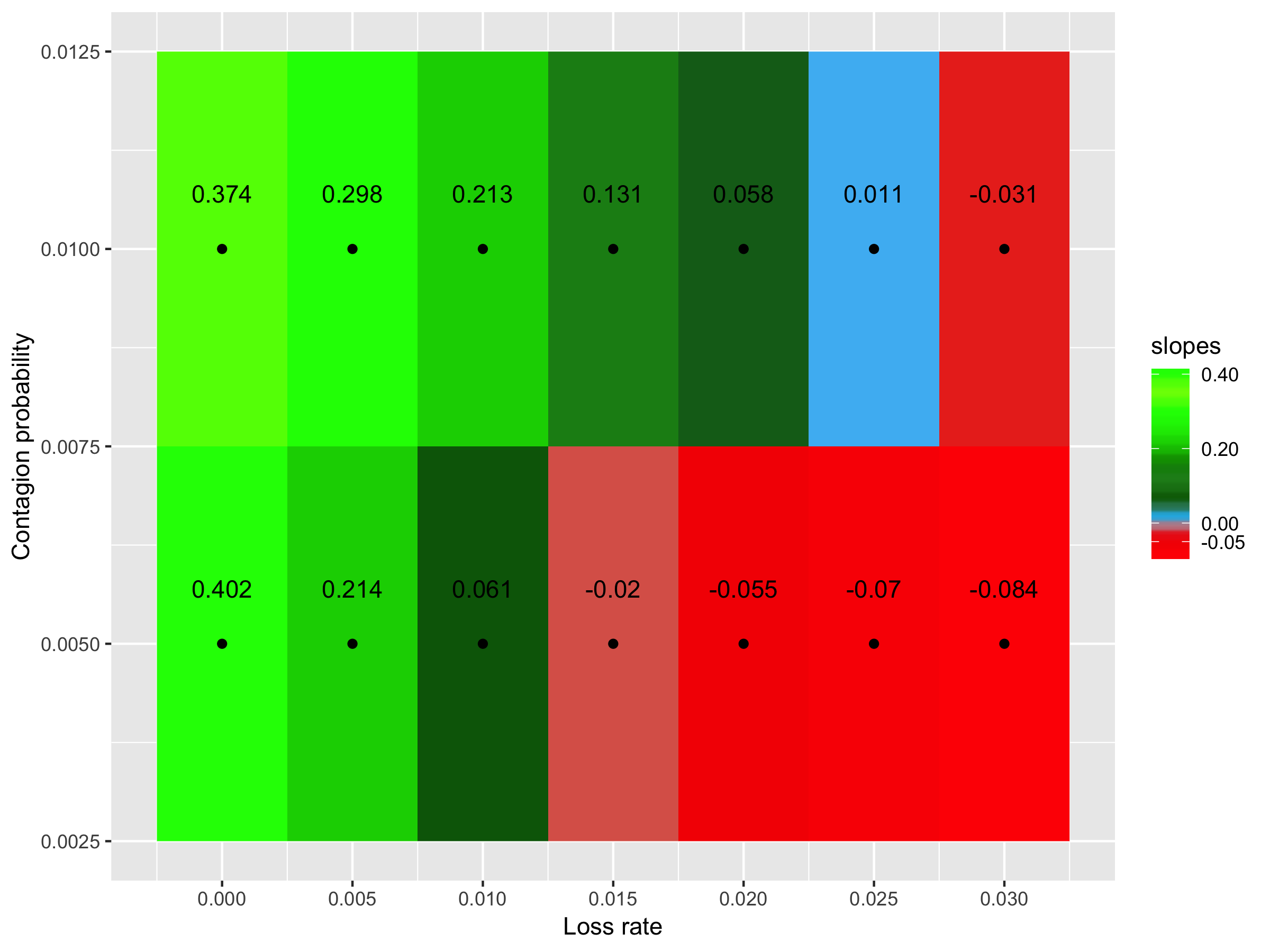

**Supp. Fig. 5.3 - Effect of considering a ratio of 2 virulence genes for 1 resistance gene.** Slope of the regression between the diversity of virulence genes and resistance genes according to the contagion probability (vertical axis) and the loss rate (horizontal axis). Green: positive slopes; and red: negative slopes.

| **Supp. Table 5.4 –** The impact of considering a ratio of 4 virulence genes for 1 resistance gene on the correlation between virulence and resistance genes. | | | | | |
| --- | --- | --- | --- | --- | --- |
| **Contagion probability (%)** | **Resistance genes loss rate (%)** | **Slope considering a ratio of 4:1** | **Slope considering a ratio of 1:1** | **P-value of T-test for differences of slopes** | **Change in the slope signal?** |
| **0.5** | 0 | 0.196 | 0.775 | 0 | No |
|  | 0.5 | 0.099 | 0.381 | 5.03x10^-192^ | No |
|  | 1 | 0.033 | 0.109 | 4.18x10^-22^ | No |
|  | 1.5 | -0.008 | -0.038 | 3.17x10^-5^ | No |
|  | 2 | -0.026 | -0.113 | 1.03x10^-35^ | No |
|  | 2.5 | -0.041 | -0.145 | 4.99x10^-54^ | No |
|  | 3 | -0.042 | -0.174 | 1.52x10^-93^ | No |
| **1** | 0 | 0.192 | 0.742 | 0 | No |
|  | 0.5 | 0.146 | 0.586 | 0 | No |
|  | 1 | 0.110 | 0.431 | 3.09x10^-224^ | No |
|  | 1.5 | 0.071 | 0.253 | 8.51x10^-80^ | No |
|  | 2 | 0.030 | 0.106 | 4.78x10^-17^ | No |
|  | 2.5 | -0.001 | -0.002 | 8.26x10^-1^ | No |
|  | 3 | -0.014 | -0.069 | 5.57x10^-16^ | No |

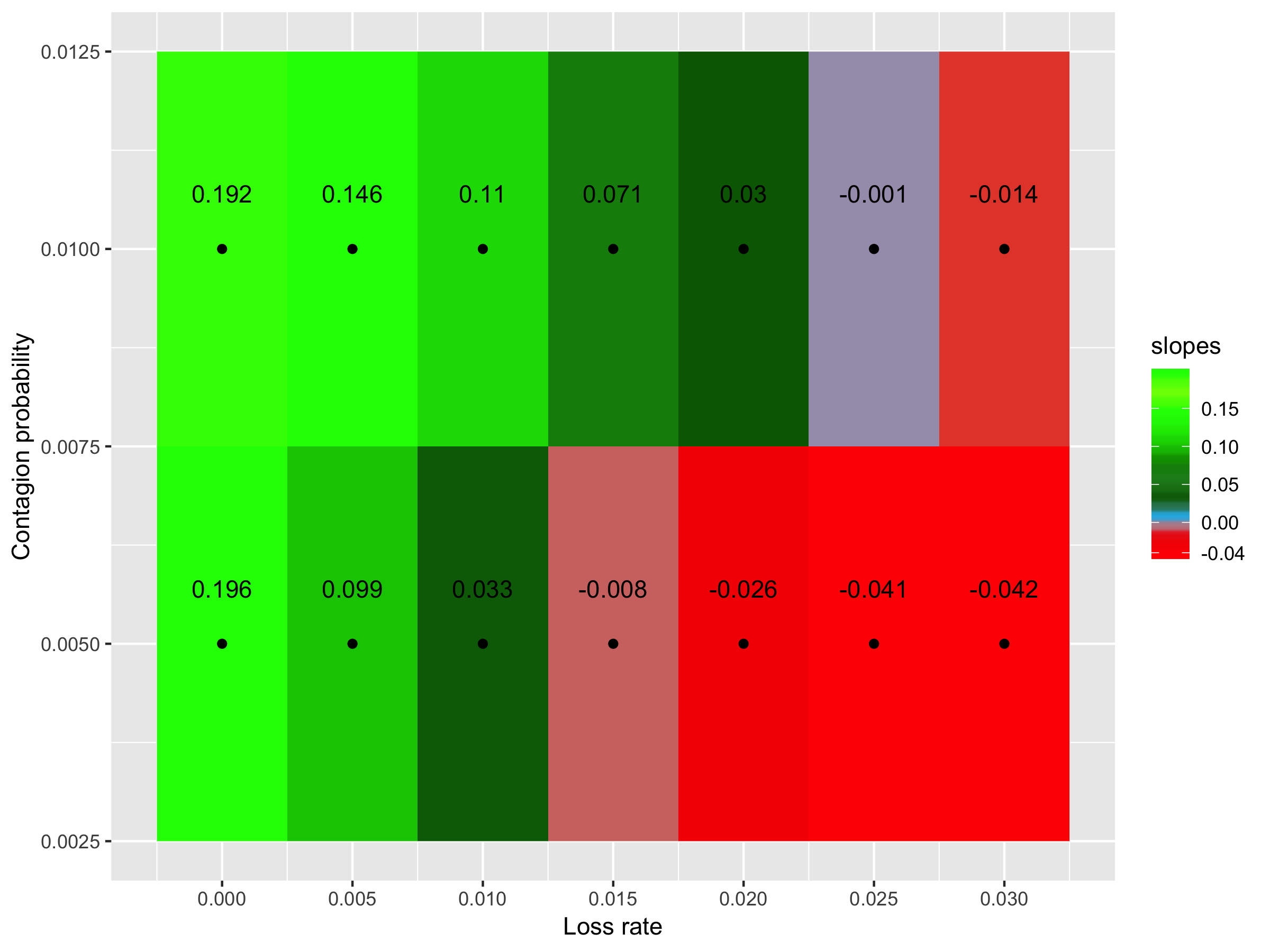

**Supp. Fig. 5.4 - Effect of considering a ratio of 4 virulence genes for 1 resistance gene.** Slope of the regression between the diversity of virulence genes and resistance genes according to the contagion probability (vertical axis) and the loss rate (horizontal axis). Green: positive slopes; and red: negative slopes.

**6 – The correlation’s signal is robust under changes in the gene elimination probability when people take antibiotics (considering equal probabilities to eliminate virulence and resistance genes)**

| **Supp. Table 6.1 –** The impact of considering a probability of eliminating genes under antibiotic intake of 30% on the correlation between virulence and resistance genes. | | | | | |
| --- | --- | --- | --- | --- | --- |
| **Contagion probability (%)** | **Resistance genes loss rate (%)** | **Slope considering a probability of 30%** | **Slope considering a probability of 70%** | **P-value of T-test for differences of slopes** | **Change in the slope signal?** |
| **0.5** | 0 | 0.663 | 0.775 | 3.16x10^-19^ | No |
|  | 0.5 | 0.303 | 0.381 | 4.48x10^-7^ | No |
|  | 1 | 0.030 | 0.109 | 1.97x10^-6^ | No |
|  | 1.5 | -0.138 | -0.038 | 6.21x10^-12^ | No |
|  | 2 | -0.245 | -0.113 | 7.68x10^-22^ | No |
|  | 2.5 | -0.288 | -0.145 | 3.29x10^-27^ | No |
|  | 3 | -0.314 | -0.174 | 1.40x10^-34^ | No |
| **1** | 0 | 0.712 | 0.742 | 2.66x10^-3^ | No |
|  | 0.5 | 0.540 | 0.586 | 6.46x10^-3^ | No |
|  | 1 | 0.354 | 0.431 | 3.44x10^-5^ | No |
|  | 1.5 | 0.250 | 0.253 | 8.89x10^-1^ | No |
|  | 2 | 0.011 | 0.106 | 1.60x10^-5^ | No |
|  | 2.5 | -0.128 | -0.002 | 2.86x10^-11^ | No |
|  | 3 | -0.227 | -0.069 | 2.20x10^-17^ | No |

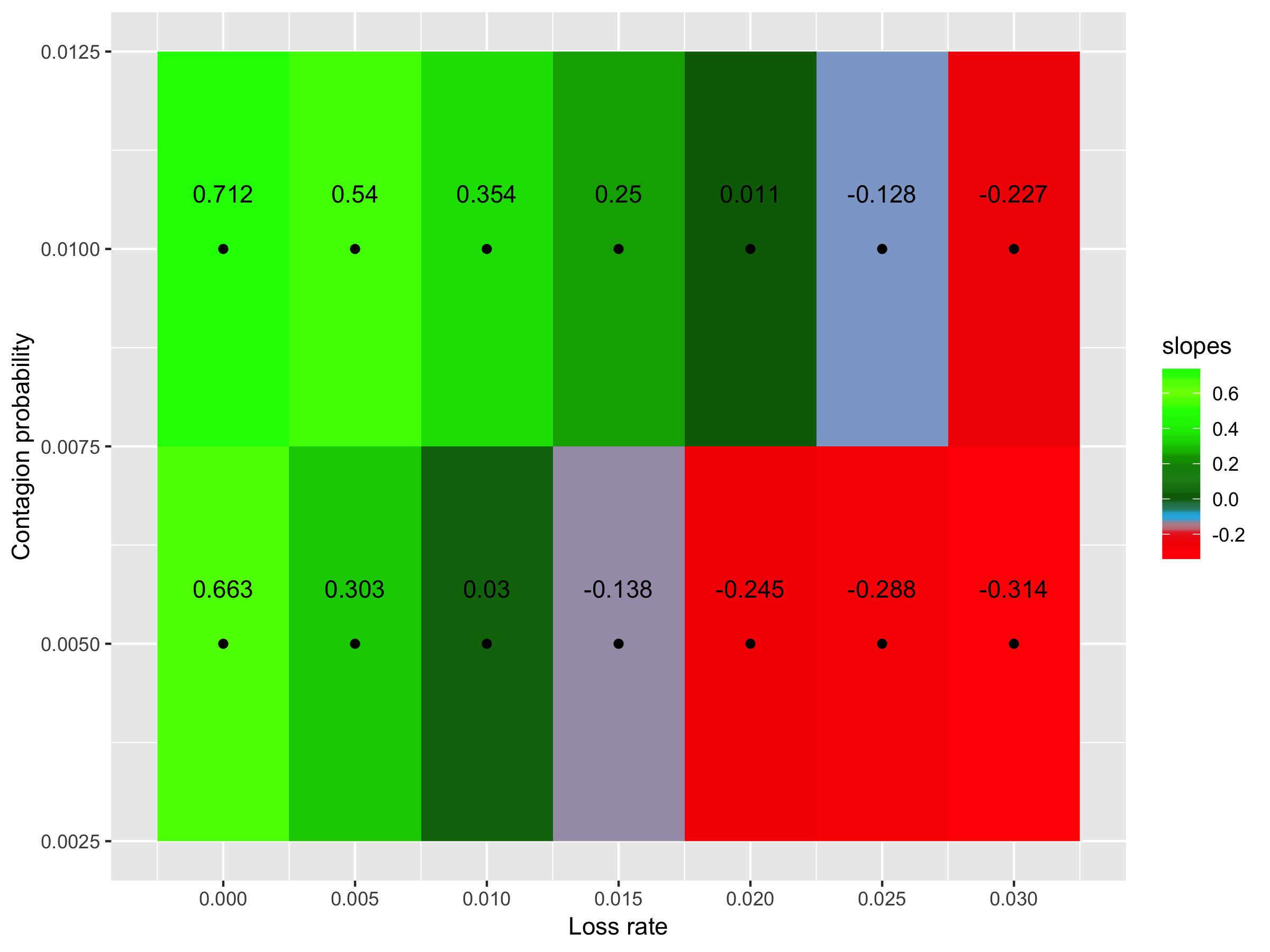

**Supp. Fig. 6.1 - Effect of considering a probability of eliminating genes under antibiotic intake of 30%.** Slope of the regression between the diversity of virulence genes and resistance genes according to the contagion probability (vertical axis) and the loss rate (horizontal axis). Green: positive slopes; and red: negative slopes.

| **Supp. Table 6.2 –** The impact of considering a probability of eliminating genes under antibiotic intake of 50% on the correlation between virulence and resistance genes. | | | | | |
| --- | --- | --- | --- | --- | --- |
| **Contagion probability (%)** | **Resistance genes loss rate (%)** | **Slope considering a probability of 50%** | **Slope considering a probability of 70%** | **P-value of T-test for differences of slopes** | **Change in the slope signal?** |
| **0.5** | 0 | 0.697 | 0.775 | 1.33x10^-9^ | No |
|  | 0.5 | 0.359 | 0.381 | 7.26x10^-2^ | No |
|  | 1 | 0.091 | 0.109 | 1.14x10^-1^ | No |
|  | 1.5 | -0.072 | -0.038 | 2.00x10^-3^ | No |
|  | 2 | -0.144 | -0.113 | 2.88x10^-3^ | No |
|  | 2.5 | -0.189 | -0.145 | 1.07x10^-5^ | No |
|  | 3 | -0.195 | -0.174 | 1.34x10^-2^ | No |
| **1** | 0 | 0.723 | 0.742 | 4.60x10^-2^ | No |
|  | 0.5 | 0.583 | 0.586 | 7.43x10^-1^ | No |
|  | 1 | 0.379 | 0.431 | 1.97x10^-4^ | No |
|  | 1.5 | 0.269 | 0.253 | 3.09x10^-1^ | No |
|  | 2 | 0.094 | 0.106 | 4.16x10^-1^ | No |
|  | 2.5 | -0.037 | -0.002 | 9.49x10^-3^ | No |
|  | 3 | -0.113 | -0.069 | 1.32x10^-4^ | No |

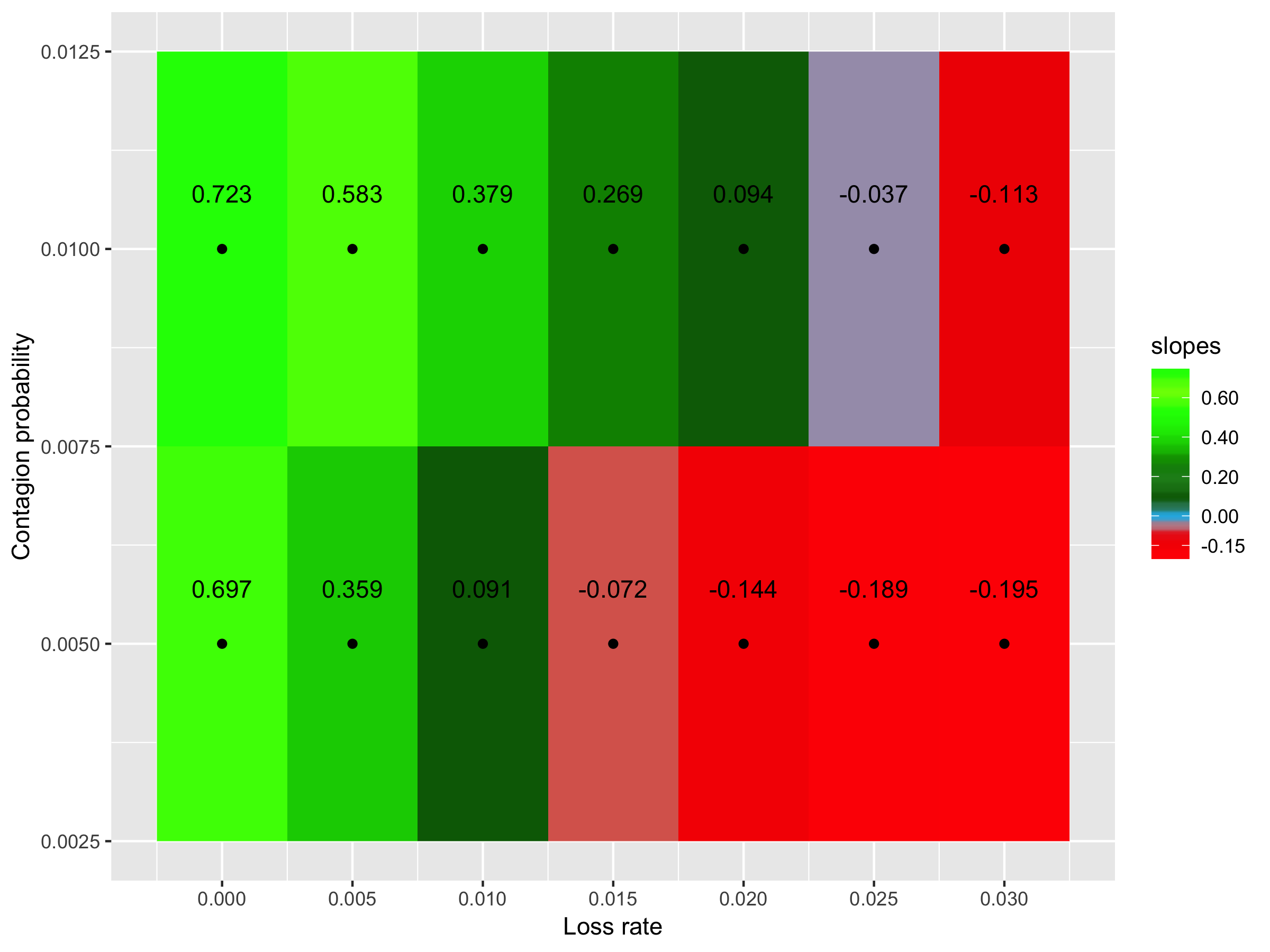

**Supp. Fig. 6.2 - Effect of considering a probability of eliminating genes under antibiotic intake of 50%.** Slope of the regression between the diversity of virulence genes and resistance genes according to the contagion probability (vertical axis) and the loss rate (horizontal axis). Green: positive slopes; and red: negative slopes.

**7 – The correlation’s signal is robust under changes in the gene elimination probability when people take antibiotics (considering different probabilities to eliminate virulence and resistance genes)**

| **Supp. Table 7.1 –** The impact of considering a probability of eliminating virulence genes under antibiotic intake of 30% and a probability of eliminating resistance genes under antibiotic intake of 50% on the correlation between virulence and resistance genes. | | | | | |
| --- | --- | --- | --- | --- | --- |
| **Contagion probability (%)** | **Resistance genes loss rate (%)** | **Slope considering a probability of 30% for virulence genes and 50% for resistance genes** | **Slope considering a probability of 70%** | **P-value of T-test for differences of slopes** | **Change in the slope signal?** |
| **0.5** | 0 | 0.943 | 0.775 | 4.23x10^-26^ | No |
|  | 0.5 | 0.500 | 0.381 | 6.64x10^-15^ | No |
|  | 1 | 0.135 | 0.109 | 1.05x10^-1^ | No |
|  | 1.5 | -0.100 | -0.038 | 1.06x10^-5^ | No |
|  | 2 | -0.210 | -0.113 | 6.56x10^-15^ | No |
|  | 2.5 | -0.244 | -0.145 | 7.54x10^-17^ | No |
|  | 3 | -0.272 | -0.174 | 1.05x10^-18^ | No |
| **1** | 0 | 1.110 | 0.742 | 8.33x10^-137^ | No |
|  | 0.5 | 0.852 | 0.586 | 4.45x10^-56^ | No |
|  | 1 | 0.645 | 0.431 | 3.50x10^-24^ | No |
|  | 1.5 | 0.355 | 0.253 | 2.67x10^-6^ | No |
|  | 2 | 0.155 | 0.106 | 1.85x10^-2^ | No |
|  | 2.5 | -0.048 | -0.002 | 9.15x10^-3^ | No |
|  | 3 | -0.183 | -0.069 | 6.14x10^-11^ | No |

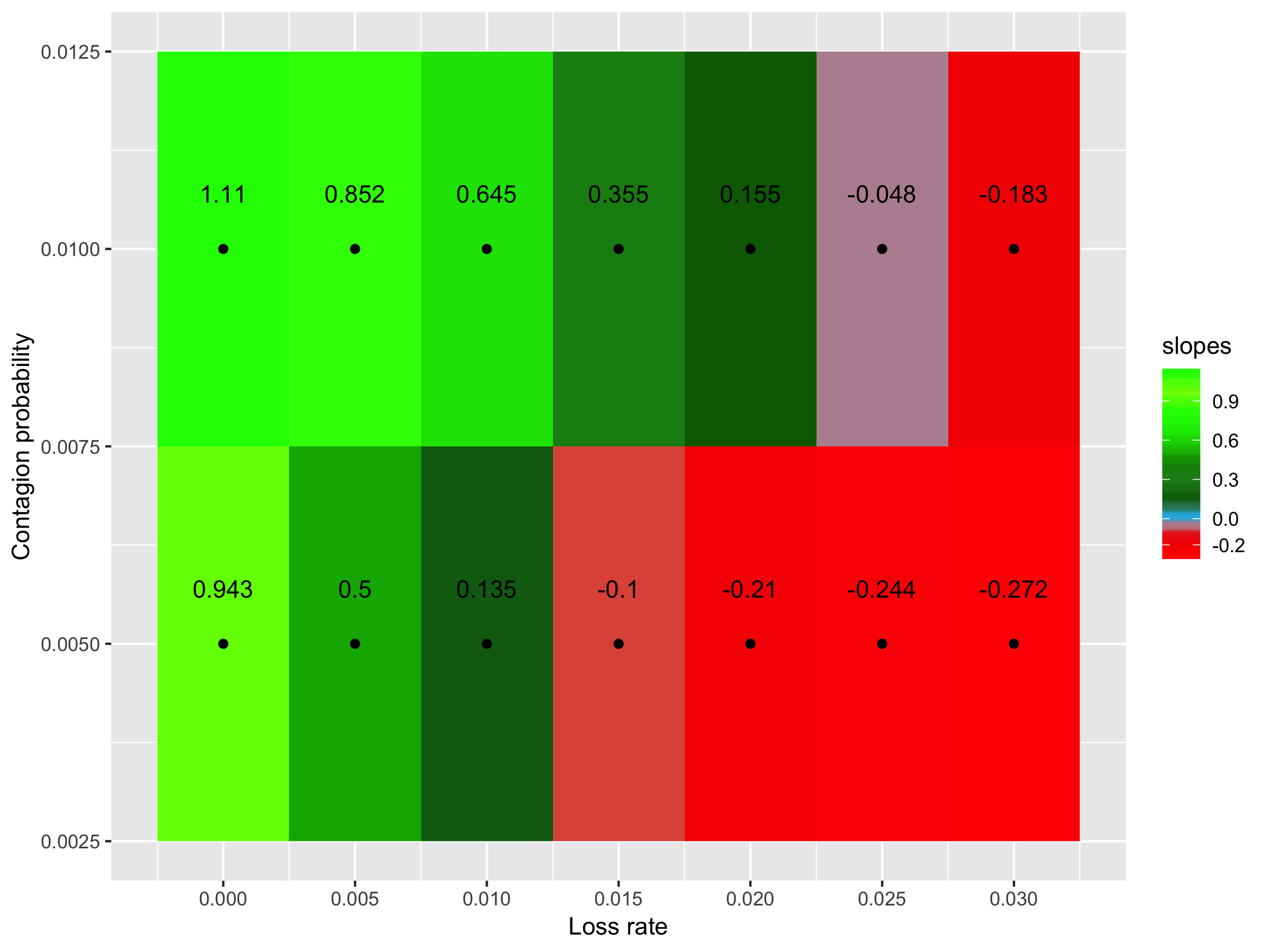

Supp. Fig. 7.1 - Effect of considering a probability of eliminating virulence genes under antibiotic intake of 30% and a probability of eliminating resistance genes under antibiotic intake of 50%. Slope of the regression between the diversity of virulence genes and resistance genes according to the contagion probability (vertical axis) and the loss rate (horizontal axis). Green: positive slopes; and red: negative slopes.

| **Supp. Table 7.2 –** The impact of considering a probability of eliminating virulence genes under antibiotic intake of 30% and a probability of eliminating resistance genes under antibiotic intake of 70% on the correlation between virulence and resistance genes. | | | | | |
| --- | --- | --- | --- | --- | --- |
| **Contagion probability (%)** | **Resistance genes loss rate (%)** | **Slope considering a probability of 30% for virulence genes and 70% for resistance genes** | **Slope considering a probability of 70%** | **P-value of T-test for differences of slopes** | **Change in the slope signal?** |
| **0.5** | 0 | 1.115 | 0.775 | 2.00x10^-65^ | No |
|  | 0.5 | 0.595 | 0.381 | 4.88x10^-39^ | No |
|  | 1 | 0.161 | 0.109 | 2.19x10^-4^ | No |
|  | 1.5 | -0.058 | -0.038 | 1.12x10^-1^ | No |
|  | 2 | -0.167 | -0.113 | 3.09x10^-6^ | No |
|  | 2.5 | -0.217 | -0.145 | 3.12x10^-11^ | No |
|  | 3 | -0.234 | -0.174 | 6.32x10^-9^ | No |
| **1** | 0 | 1.466 | 0.742 | 4.11x10^-271^ | No |
|  | 0.5 | 1.078 | 0.586 | 1.76x10^-141^ | No |
|  | 1 | 0.838 | 0.431 | 1.03x10^-68^ | No |
|  | 1.5 | 0.419 | 0.253 | 7.25x10^-17^ | No |
|  | 2 | 0.252 | 0.106 | 4.1110^-12^ | No |
|  | 2.5 | 0.016 | -0.002 | 3.00x10^-1^ | Yes |
|  | 3 | -0.100 | -0.069 | 3.11x10^-2^ | No |

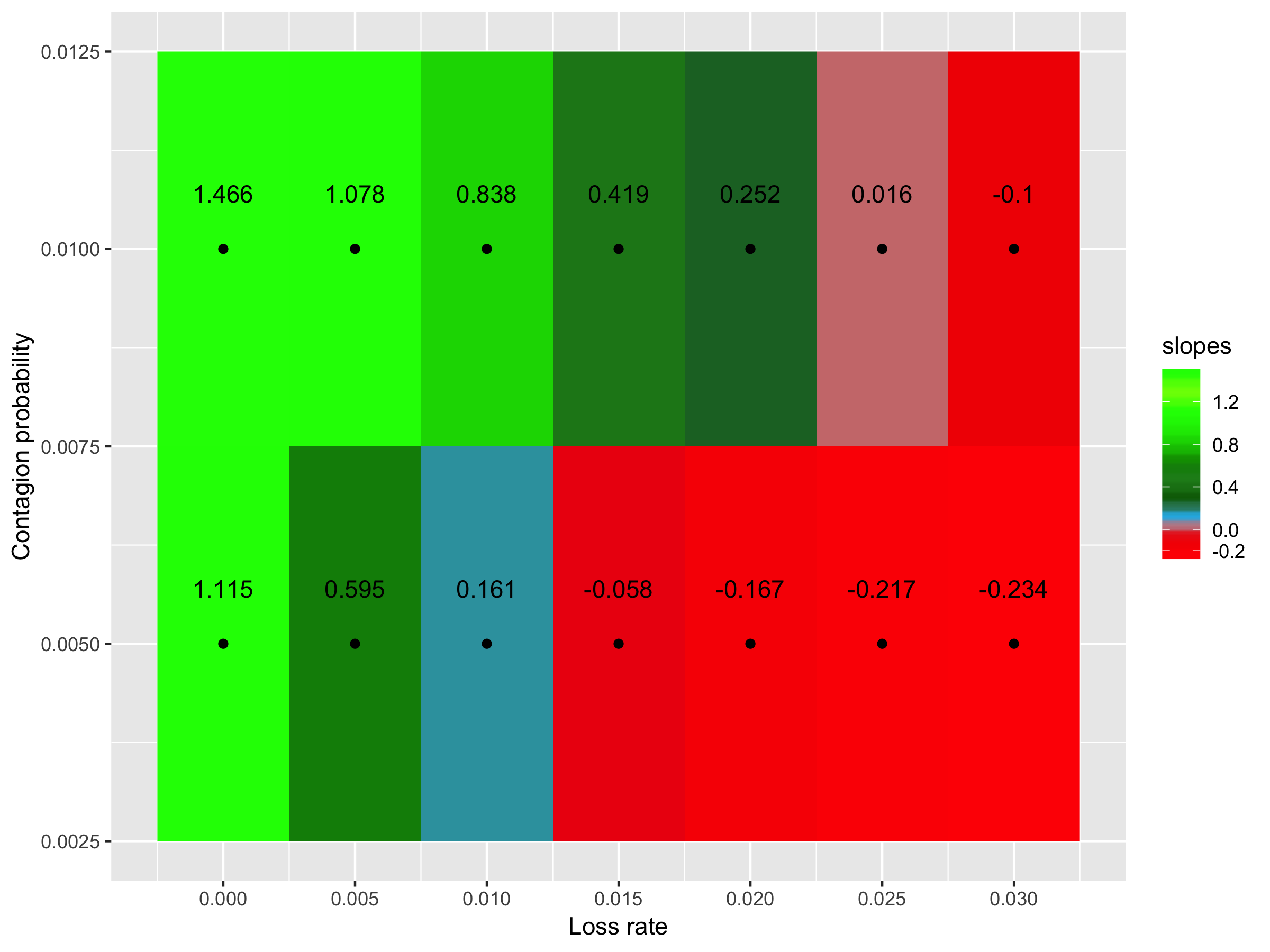

Supp. Fig. 7.2 - Effect of considering a probability of eliminating virulence genes under antibiotic intake of 30% and a probability of eliminating resistance genes under antibiotic intake of 70%. Slope of the regression between the diversity of virulence genes and resistance genes according to the contagion probability (vertical axis) and the loss rate (horizontal axis). Green: positive slopes; and red: negative slopes.

| **Supp. Table 7.3 –** The impact of considering a probability of eliminating virulence genes under antibiotic intake of 50% and a probability of eliminating resistance genes under antibiotic intake of 30% on the correlation between virulence and resistance genes. | | | | | |
| --- | --- | --- | --- | --- | --- |
| **Contagion probability (%)** | **Resistance genes loss rate (%)** | **Slope considering a probability of 50% for virulence genes and 30% for resistance genes** | **Slope considering a probability of 70%** | **P-value of T-test for differences of slopes** | **Change in the slope signal?** |
| **0.5** | 0 | 0.467 | 0.775 | 6.04x10^-138^ | No |
|  | 0.5 | 0.262 | 0.381 | 5.73x10^-22^ | No |
|  | 1 | 0.037 | 0.109 | 4.59x10^-8^ | No |
|  | 1.5 | -0.111 | -0.038 | 2.98x10^-10^ | No |
|  | 2 | -0.192 | -0.113 | 1.73x10^-13^ | No |
|  | 2.5 | -0.224 | -0.145 | 6.39x10^-16^ | No |
|  | 3 | -0.231 | -0.174 | 1.07x10^-9^ | No |
| **1** | 0 | 0.465 | 0.742 | 3.90x10^-199^ | No |
|  | 0.5 | 0.360 | 0.586 | 9.46x10^-81^ | No |
|  | 1 | 0.277 | 0.431 | 5.37x10^-25^ | No |
|  | 1.5 | 0.110 | 0.253 | 2.06x10^-20^ | No |
|  | 2 | 0.024 | 0.106 | 1.51x10^-7^ | No |
|  | 2.5 | -0.088 | -0.002 | 2.98x10^-9^ | No |
|  | 3 | -0.137 | -0.069 | 2.60x10^-8^ | No |

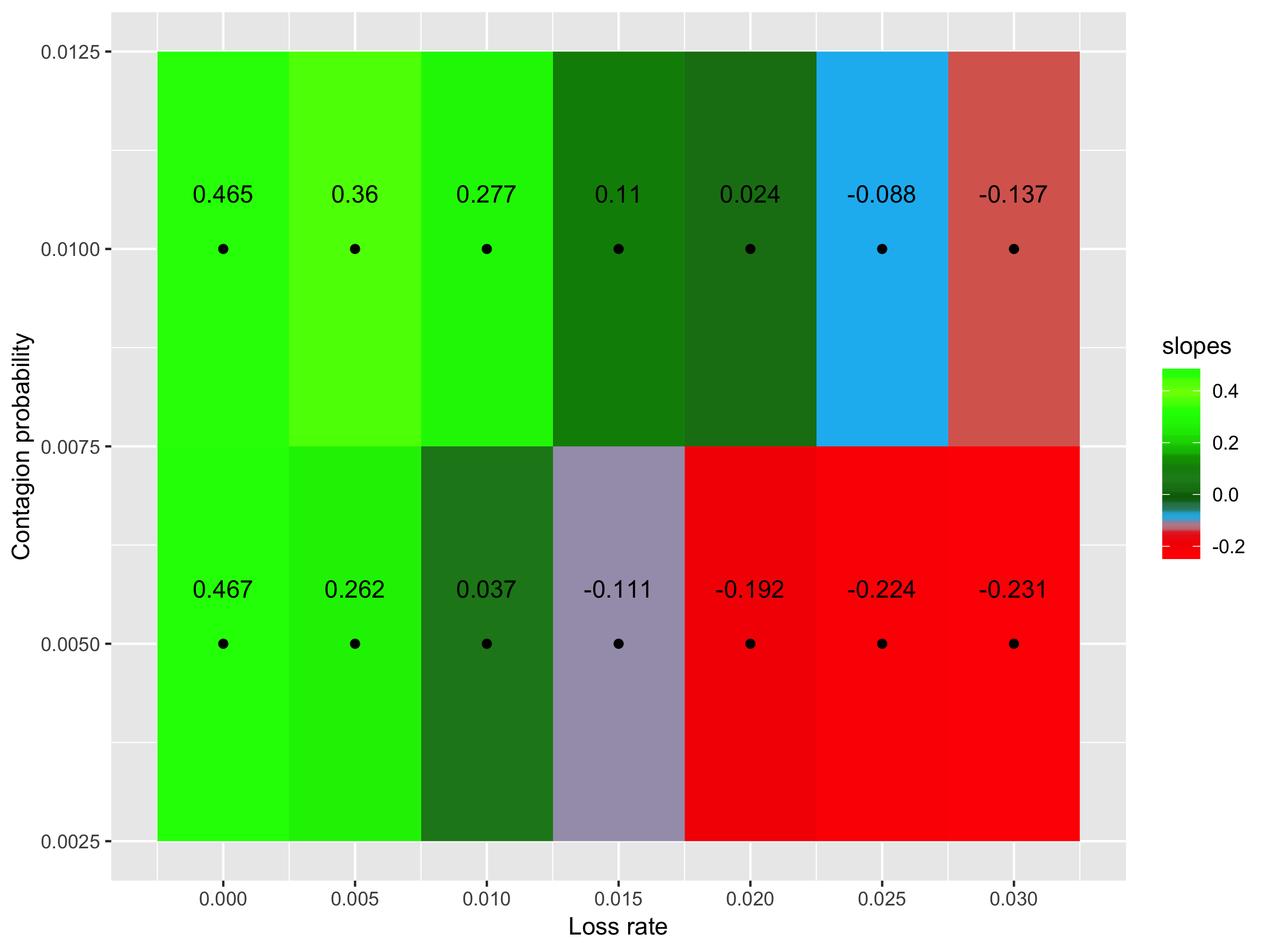

Supp. Fig. 7.3 - Effect of considering a probability of eliminating virulence genes under antibiotic intake of 50% and a probability of eliminating resistance genes under antibiotic intake of 30%. Slope of the regression between the diversity of virulence genes and resistance genes according to the contagion probability (vertical axis) and the loss rate (horizontal axis). Green: positive slopes; and red: negative slopes.

| **Supp. Table 7.4 –** The impact of considering a probability of eliminating virulence genes under antibiotic intake of 50% and a probability of eliminating resistance genes under antibiotic intake of 70% on the correlation between virulence and resistance genes. | | | | | |
| --- | --- | --- | --- | --- | --- |
| **Contagion probability (%)** | **Resistance genes loss rate (%)** | **Slope considering a probability of 50% for virulence genes and 70% for resistance genes** | **Slope considering a probability of 70%** | **P-value of T-test for differences of slopes** | **Change in the slope signal?** |
| **0.5** | 0 | 0.836 | 0.775 | 8.71x10^-6^ | No |
|  | 0.5 | 0.437 | 0.381 | 4.09x10^-6^ | No |
|  | 1 | 0.120 | 0.109 | 3.44x10^-1^ | No |
|  | 1.5 | -0.041 | -0.038 | 7.61x10^-1^ | No |
|  | 2 | -0.121 | -0.113 | 3.90x10^-1^ | No |
|  | 2.5 | -0.173 | -0.145 | 1.27x10^-3^ | No |
|  | 3 | -0.174 | -0.174 | 9.56x10^-1^ | No |
| **1** | 0 | 0.945 | 0.742 | 1.63x10^-79^ | No |
|  | 0.5 | 0.739 | 0.586 | 2.41x10^-35^ | No |
|  | 1 | 0.521 | 0.431 | 1.87x10^-10^ | No |
|  | 1.5 | 0.346 | 0.253 | 3.00x10^-9^ | No |
|  | 2 | 0.151 | 0.106 | 1.42x10^-3^ | No |
|  | 2.5 | 0.016 | -0.002 | 1.50x10^-1^ | Yes |
|  | 3 | -0.068 | -0.069 | 9.16x10^-1^ | No |

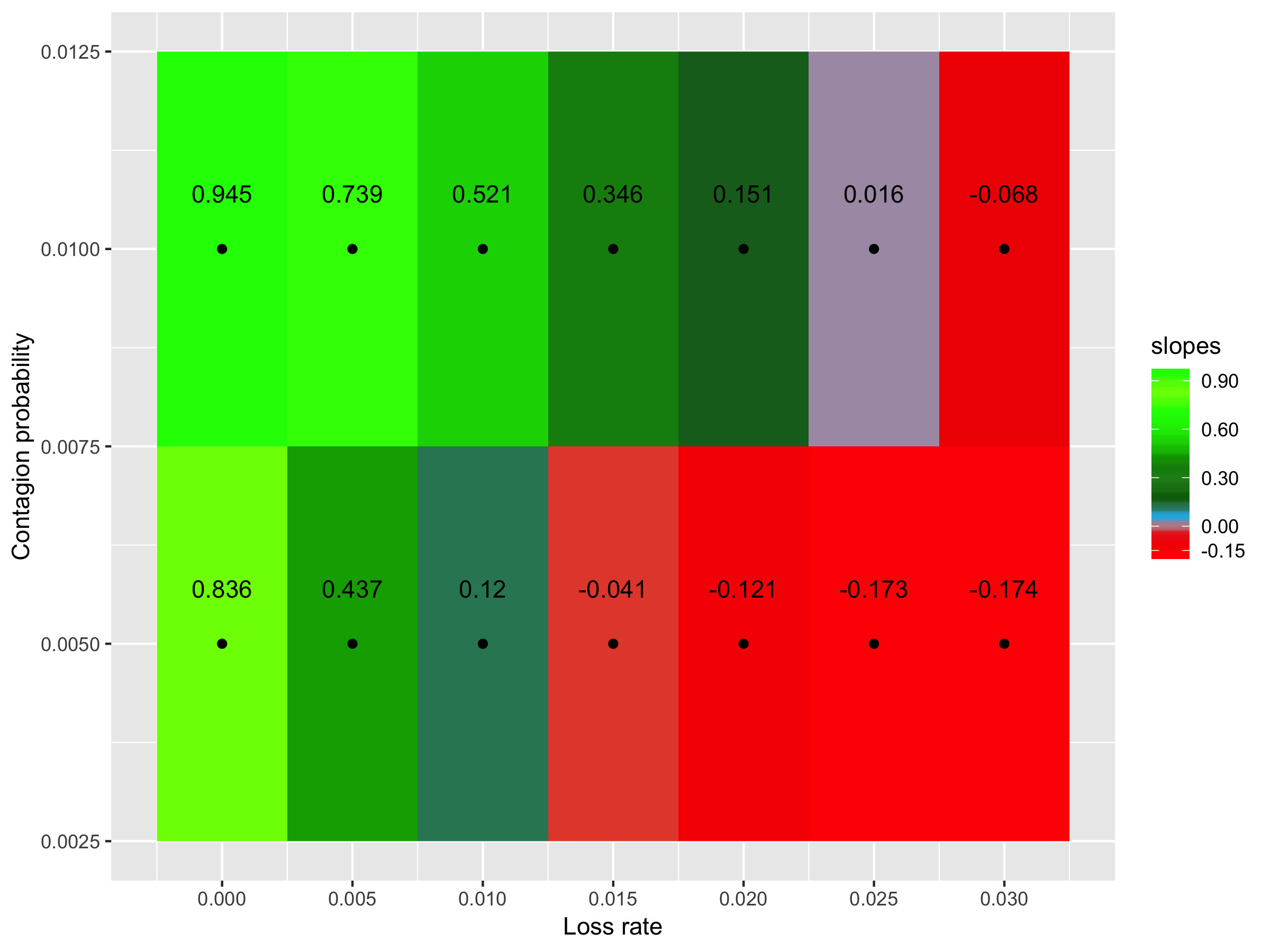

Supp. Fig. 7.4 - Effect of considering a probability of eliminating virulence genes under antibiotic intake of 50% and a probability of eliminating resistance genes under antibiotic intake of 70%. Slope of the regression between the diversity of virulence genes and resistance genes according to the contagion probability (vertical axis) and the loss rate (horizontal axis). Green: positive slopes; and red: negative slopes.

| **Supp. Table 7.5 –** The impact of considering a probability of eliminating virulence genes under antibiotic intake of 70% and a probability of eliminating resistance genes under antibiotic intake of 30% on the correlation between virulence and resistance genes. | | | | | |
| --- | --- | --- | --- | --- | --- |
| **Contagion probability (%)** | **Resistance genes loss rate (%)** | **Slope considering a probability of 70% for virulence genes and 30% for resistance genes** | **Slope considering a probability of 70%** | **P-value of T-test for differences of slopes** | **Change in the slope signal?** |
| **0.5** | 0 | 0.418 | 0.775 | 2.96x10^-153^ | No |
|  | 0.5 | 0.217 | 0.381 | 3.75x10^-40^ | No |
|  | 1 | 0.016 | 0.109 | 1.95x10^-13^ | No |
|  | 1.5 | -0.101 | -0.038 | 1.81x10^-7^ | No |
|  | 2 | -0.189 | -0.113 | 1.38x10^-11^ | No |
|  | 2.5 | -0.182 | -0.145 | 7.19x10^-5^ | No |
|  | 3 | -0.195 | -0.174 | 1.97x10^-2^ | No |
| **1** | 0 | 0.364 | 0.742 | 0 | No |
|  | 0.5 | 0.284 | 0.586 | 7.17x10^-179^ | No |
|  | 1 | 0.181 | 0.431 | 2.58x10^-82^ | No |
|  | 1.5 | 0.099 | 0.253 | 1.04x10^-29^ | No |
|  | 2 | 0.010 | 0.106 | 1.55x10^-13^ | No |
|  | 2.5 | -0.060 | -0.002 | 1.36x10^-16^ | No |
|  | 3 | -0.135 | -0.069 | 8.50x10^-11^ | No |

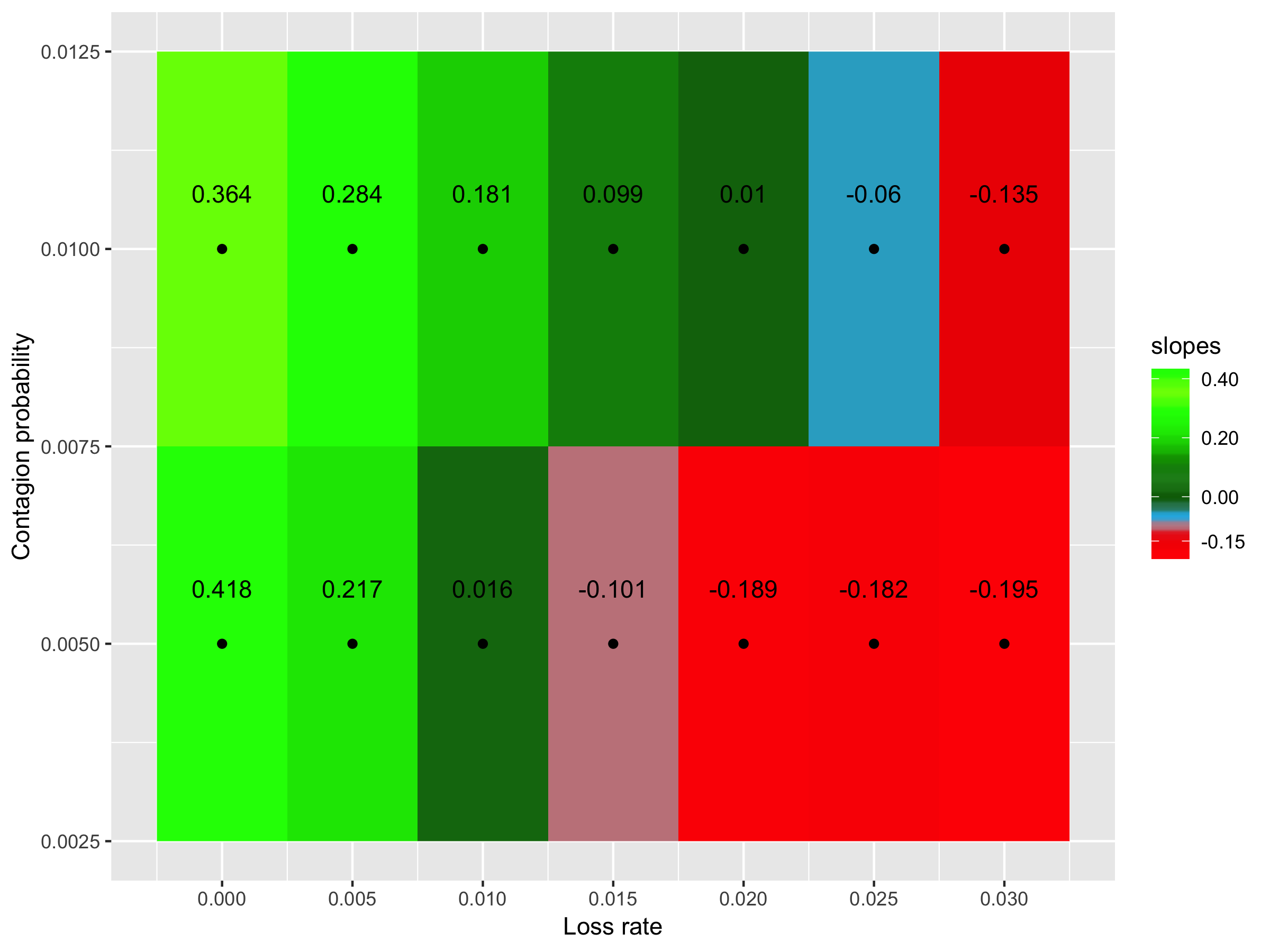

Supp. Fig. 7.5 - Effect of considering a probability of eliminating virulence genes under antibiotic intake of 70% and a probability of eliminating resistance genes under antibiotic intake of 30%. Slope of the regression between the diversity of virulence genes and resistance genes according to the contagion probability (vertical axis) and the loss rate (horizontal axis). Green: positive slopes; and red: negative slopes.

| **Supp. Table 7.6 –** The impact of considering a probability of eliminating virulence genes under antibiotic intake of 70% and a probability of eliminating resistance genes under antibiotic intake of 50% on the correlation between virulence and resistance genes. | | | | | |
| --- | --- | --- | --- | --- | --- |
| **Contagion probability (%)** | **Resistance genes loss rate (%)** | **Slope considering a probability of 70% for virulence genes and 50% for resistance genes** | **Slope considering a probability of 70%** | **P-value of T-test for differences of slopes** | **Change in the slope signal?** |
| **0.5** | 0 | 0.627 | 0.775 | 5.91x10^-29^ | No |
|  | 0.5 | 0.351 | 0.381 | 1.44x10^-2^ | No |
|  | 1 | 0.087 | 0.109 | 5.40x10^-2^ | No |
|  | 1.5 | -0.064 | -0.038 | 1.61x10^-2^ | No |
|  | 2 | -0.144 | -0.113 | 1.43x10^-3^ | No |
|  | 2.5 | -0.165 | -0.145 | 3.39x10^-2^ | No |
|  | 3 | -0.178 | -0.174 | 6.10x10^-1^ | No |
| **1** | 0 | 0.563 | 0.742 | 1.73x10^-99^ | No |
|  | 0.5 | 0.439 | 0.586 | 6.75x10^-48^ | No |
|  | 1 | 0.314 | 0.431 | 6.64x10^-21^ | No |
|  | 1.5 | 0.196 | 0.253 | 5.59x10^-6^ | No |
|  | 2 | 0.073 | 0.106 | 1.30x10^-2^ | No |
|  | 2.5 | -0.033 | -0.002 | 8.33x10^-3^ | No |
|  | 3 | -0.094 | -0.069 | 1.08x10^-2^ | No |

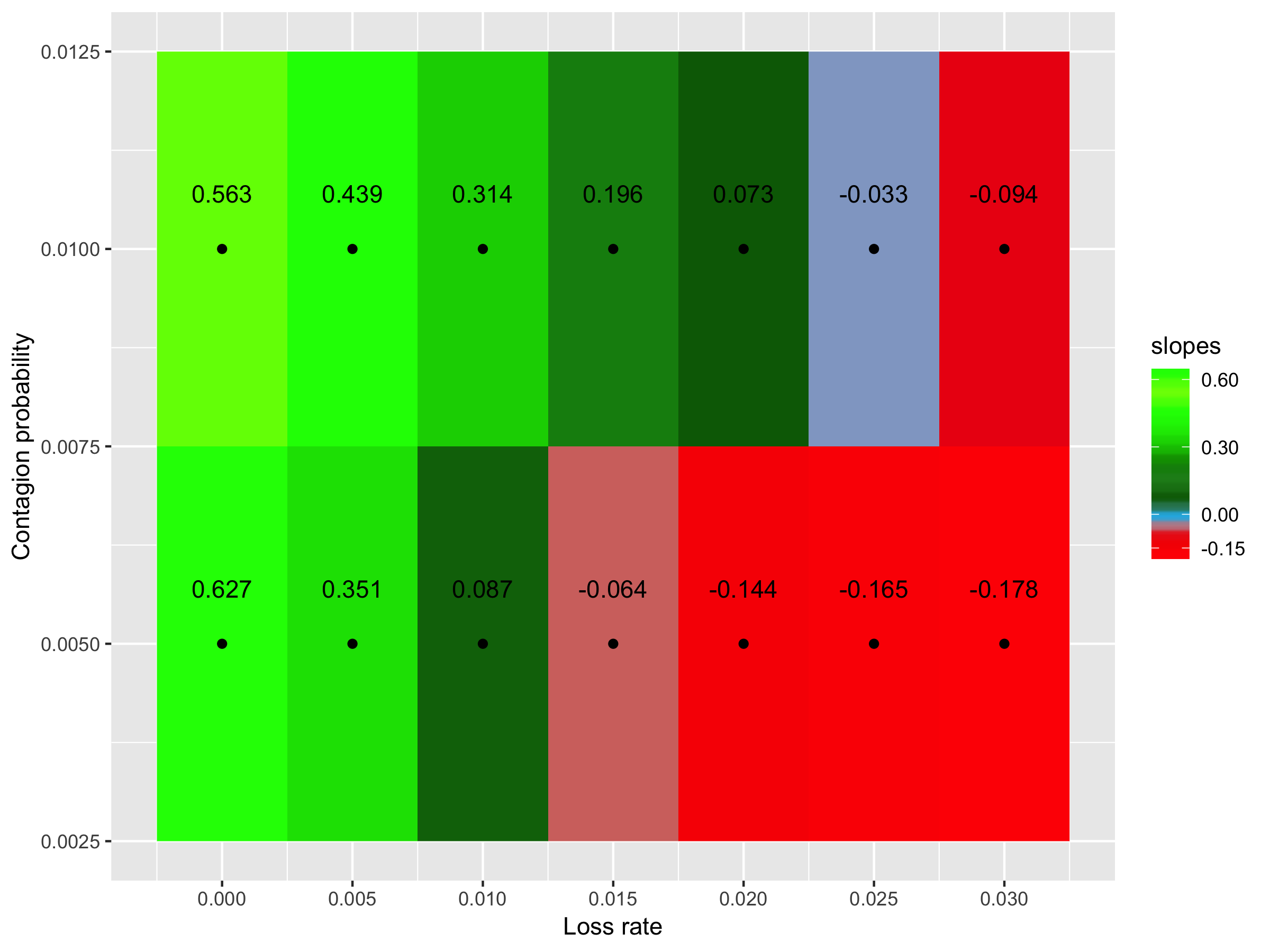

Supp. Fig. 7.6 - Effect of considering a probability of eliminating virulence genes under antibiotic intake of 70% and a probability of eliminating resistance genes under antibiotic intake of 50%. Slope of the regression between the diversity of virulence genes and resistance genes according to the contagion probability (vertical axis) and the loss rate (horizontal axis). Green: positive slopes; and red: negative slopes.

**8 – The initial proportion of metagenomes containing antibiotic resistance genes has no impact on correlations’ signal**

| **Supp. Table 8.1 –** The impact of considering that, initially, only 10% of metagenomes contain antibiotic resistance genes on the correlation between virulence and resistance genes. | | | | | |
| --- | --- | --- | --- | --- | --- |
| **Contagion probability (%)** | **Resistance genes loss rate (%)** | **Slope considering 10% metagenomes containing resistance genes initially** | **Slope considering all metagenomes containing resistance genes initially** | **P-value of T-test for differences of slopes** | **Change in the slope signal?** |
| **0.5** | 0 | 0.774 | 0.775 | 9.40x10^-1^ | No |
|  | 0.5 | 0.394 | 0.381 | 2.68x10^-1^ | No |
|  | 1 | 0.129 | 0.109 | 6.77x10^-2^ | No |
|  | 1.5 | -0.038 | -0.038 | 9.79x10^-1^ | No |
|  | 2 | -0.113 | -0.113 | 9.88x10^-1^ | No |
|  | 2.5 | -0.146 | -0.145 | 9.08x10^-1^ | No |
|  | 3 | -0.157 | -0.174 | 4.44x10^-2^ | No |
| **1** | 0 | 0.748 | 0.742 | 4.66x10^-1^ | No |
|  | 0.5 | 0.586 | 0.586 | 9.34x10^-1^ | No |
|  | 1 | 0.423 | 0.431 | 4.84x10^-1^ | No |
|  | 1.5 | 0.270 | 0.253 | 1.79x10^-1^ | No |
|  | 2 | 0.119 | 0.106 | 2.86x10^-1^ | No |
|  | 2.5 | 0.006 | -0.002 | 4.40x10^-1^ | Yes |
|  | 3 | -0.058 | -0.069 | 2.55 x10^-1^ | No |

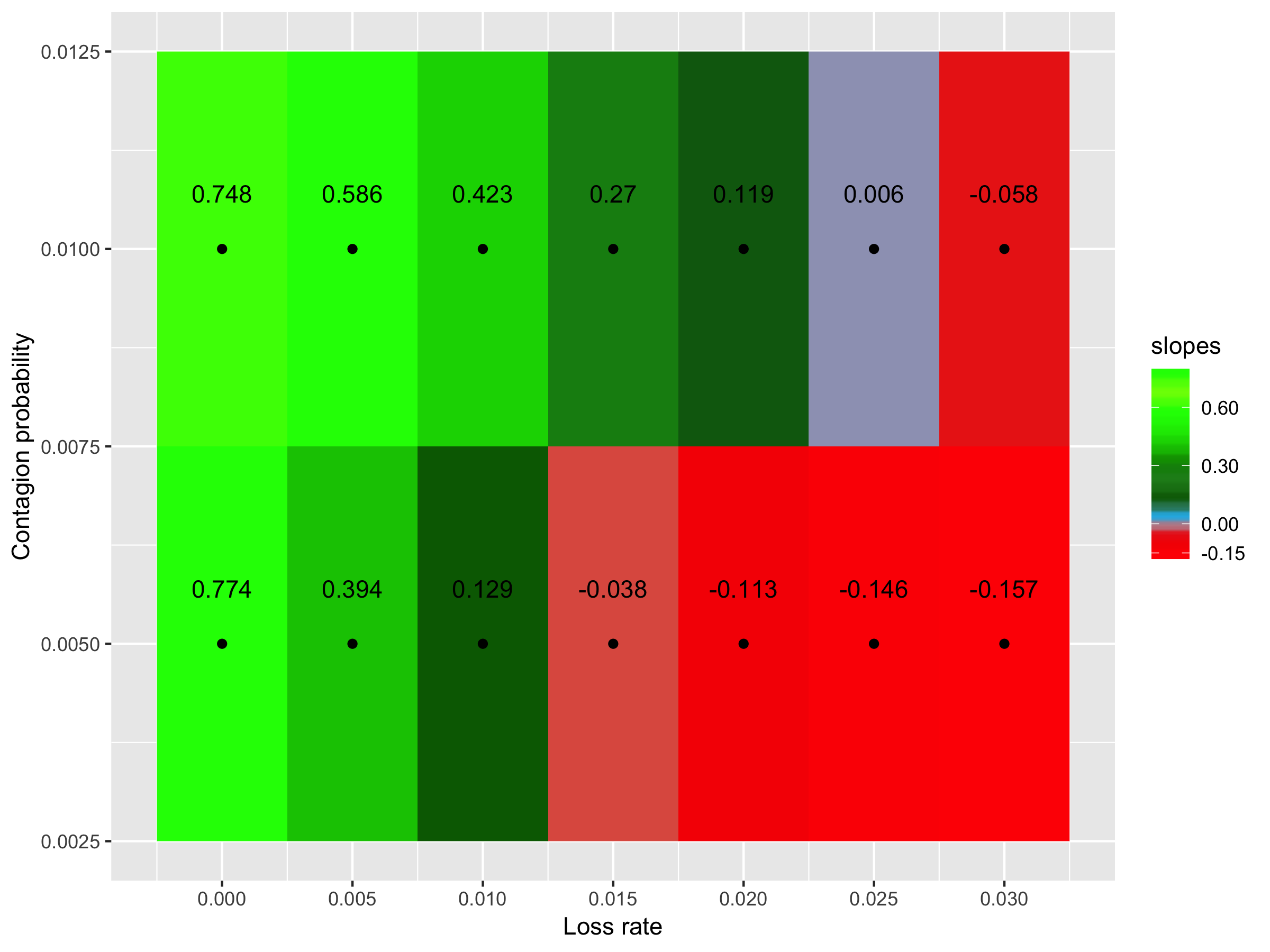

**Supp. Fig. 8.1 - Effect of considering that, initially, only 10% of metagenomes contain antibiotic resistance genes.** Slope of the regression between the diversity of virulence genes and resistance genes according to the contagion probability (vertical axis) and the loss rate (horizontal axis). Green: positive slopes; and red: negative slopes.

**9 – The network type has no impact on correlation’s signal**

| **Supp. Table 9.1 –** The impact of considering a random network (p=1) on the correlation between virulence and resistance genes. | | | | | |
| --- | --- | --- | --- | --- | --- |
| **Contagion probability (%)** | **Resistance genes loss rate (%)** | **Slope considering a random network (p=1)** | **Slope considering a small-world network (p=0.5)** | **P-value of T-test for differences of slopes** | **Change in the slope signal?** |
| **0.5** | 0 | 0.789 | 0.775 | 3.10x10^-1^ | No |
|  | 0.5 | 0.425 | 0.381 | 1.57x10^-4^ | No |
|  | 1 | 0.105 | 0.109 | 6.59x10^-1^ | No |
|  | 1.5 | -0.014 | -0.038 | 2.77x10^-2^ | No |
|  | 2 | -0.113 | -0.113 | 9.78x10^-1^ | No |
|  | 2.5 | -0.150 | -0.145 | 6.17x10^-1^ | No |
|  | 3 | -0.149 | -0.174 | 5.59x10^-3^ | No |
| **1** | 0 | 0.745 | 0.742 | 7.47x10^-1^ | No |
|  | 0.5 | 0.575 | 0.586 | 2.40x10^-1^ | No |
|  | 1 | 0.412 | 0.431 | 1.24x10^-1^ | No |
|  | 1.5 | 0.272 | 0.253 | 1.31x10^-1^ | No |
|  | 2 | 0.133 | 0.106 | 3.21x10^-2^ | No |
|  | 2.5 | 0.023 | -0.002 | 2.06x10^-2^ | Yes |
|  | 3 | -0.059 | -0.069 | 3.13x10^-1^ | No |

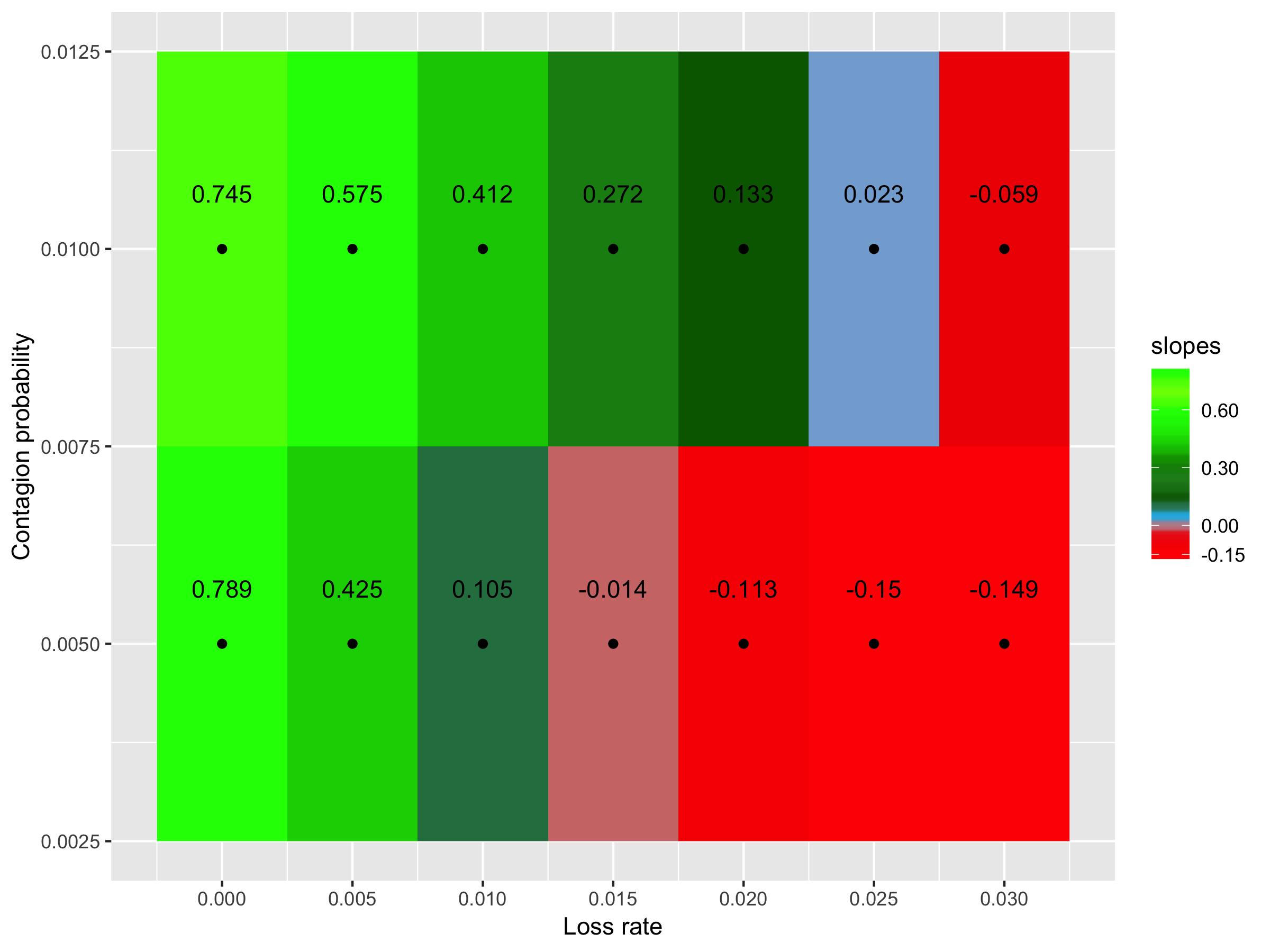

**Supp. Fig. 9.1 - Effect of considering a random network (p=1).** Slope of the regression between the diversity of virulence genes and resistance genes according to the contagion probability (vertical axis) and the loss rate (horizontal axis). Green: positive slopes; and red: negative slopes.

| **Supp. Table 9.2 –** The impact of considering a regular network (p=0) on the correlation between virulence and resistance genes. | | | | | |
| --- | --- | --- | --- | --- | --- |
| **Contagion probability (%)** | **Resistance genes loss rate (%)** | **Slope considering a regular network (p=0)** | **Slope considering a small-world network (p=0.5)** | **P-value of T-test for differences of slopes** | **Change in the slope signal?** |
| **0.5** | 0 | 7.062 | 0.775 | 7.48x10^-38^ | No |
|  | 0.005 | 2.589 | 0.381 | 3.31x10^-21^ | No |
|  | 0.01 | 0.569 | 0.109 | 1.23x10^-4^ | No |
|  | 0.015 | -0.93 | -0.038 | 1.46x10^-11^ | No |
|  | 0.02 | -1.135 | -0.113 | 1.23x10^-16^ | No |
|  | 0.025 | -1.304 | -0.145 | 4.17x10^-25^ | No |
|  | 0.03 | -1.076 | -0.174 | 1.81x10^-16^ | No |
| **1** | 0 | 0.729 | 0.742 | 1.39x10^-1^ | No |
|  | 0.005 | 0.602 | 0.586 | 8.97x10^-2^ | No |
|  | 0.01 | 0.394 | 0.431 | 6.77x10^-4^ | No |
|  | 0.015 | 0.193 | 0.253 | 3.66x10^-8^ | No |
|  | 0.02 | 0.02 | 0.106 | 5.90x10^-17^ | No |
|  | 0.025 | -0.08 | -0.002 | 8.76x10^-17^ | No |
|  | 0.03 | -0.12 | -0.069 | 1.86x10^-10^ | No |

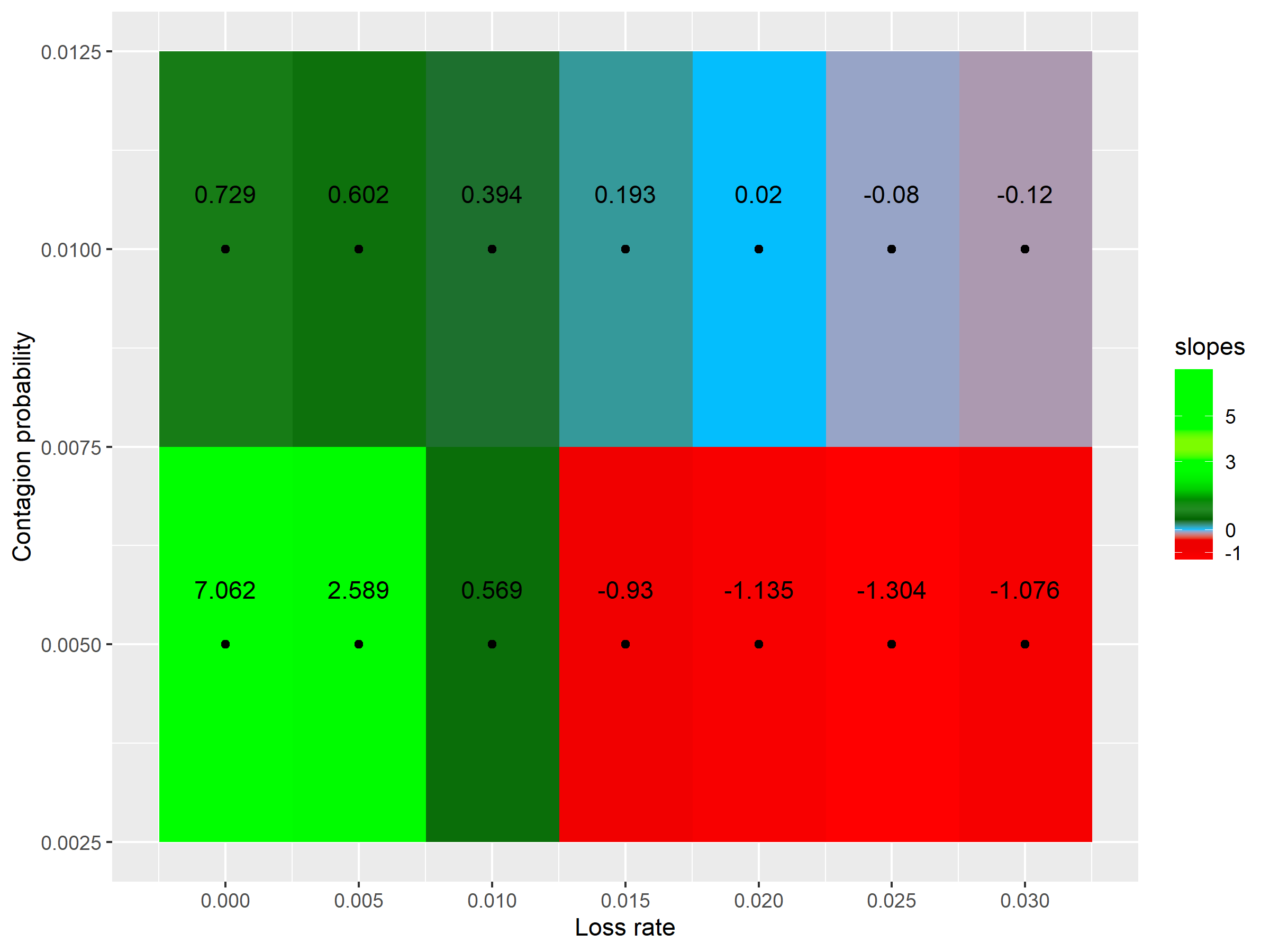

**Supp. Fig. 9.2 - Effect of considering a regular network (p=0).** Slope of the regression between the diversity of virulence genes and resistance genes according to the contagion probability (vertical axis) and the loss rate (horizontal axis). Green: positive slopes; and red: negative slopes.

|  | **500 cycles** | **2500 cycles** | **7500 cycles** | **10000 cycles** |
| --- | --- | --- | --- | --- |
| **Network p=0** | 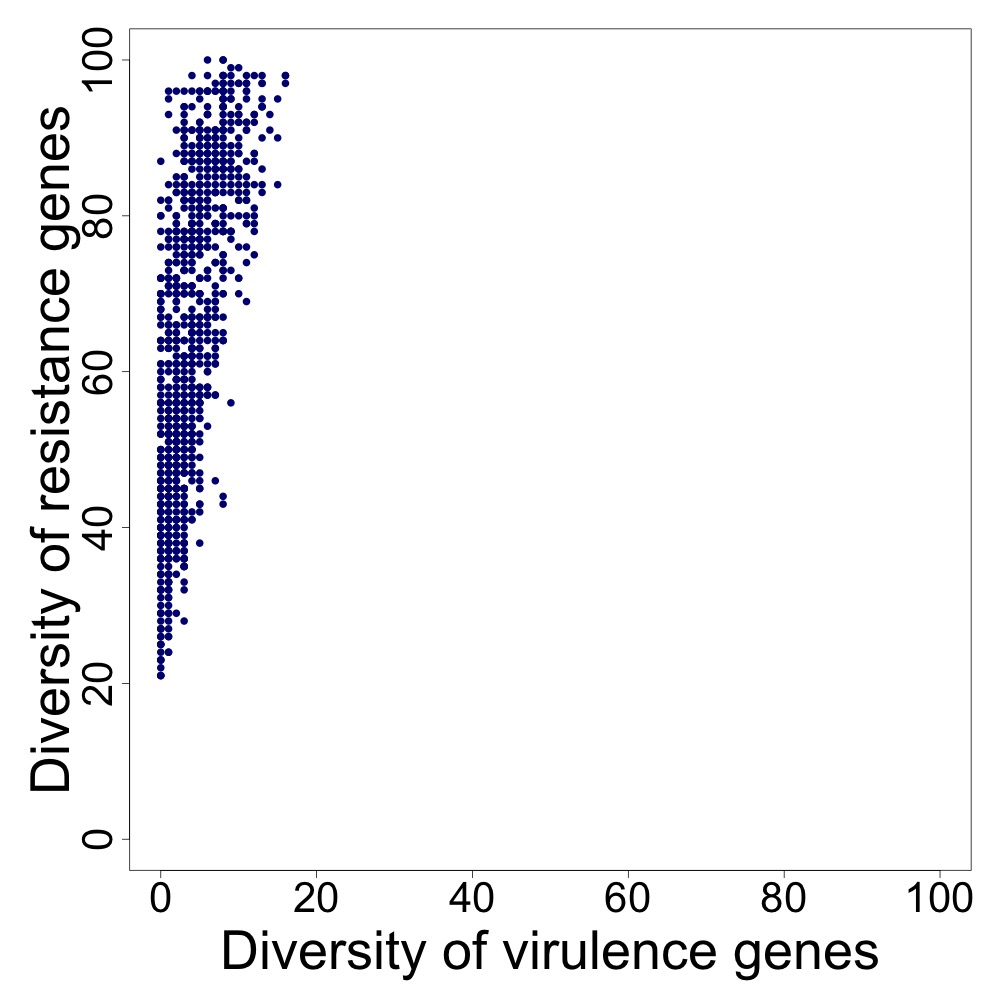 | 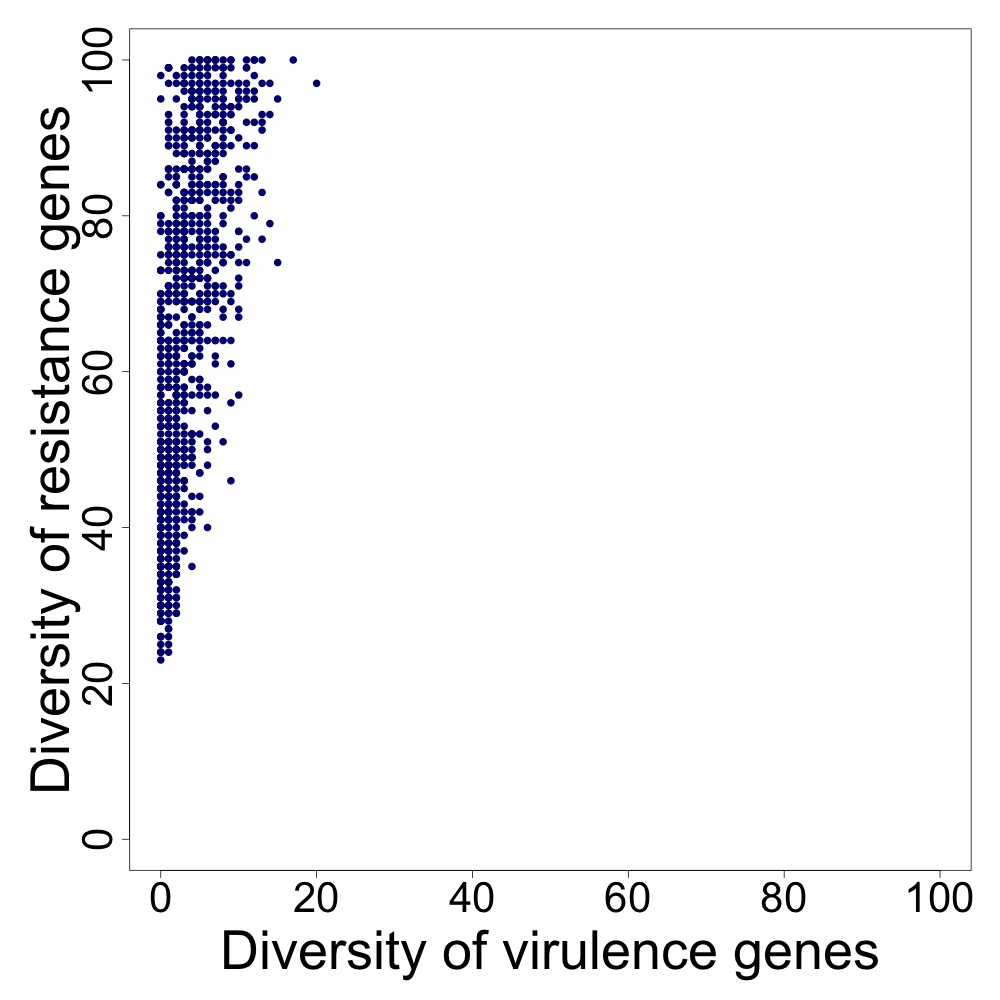 | 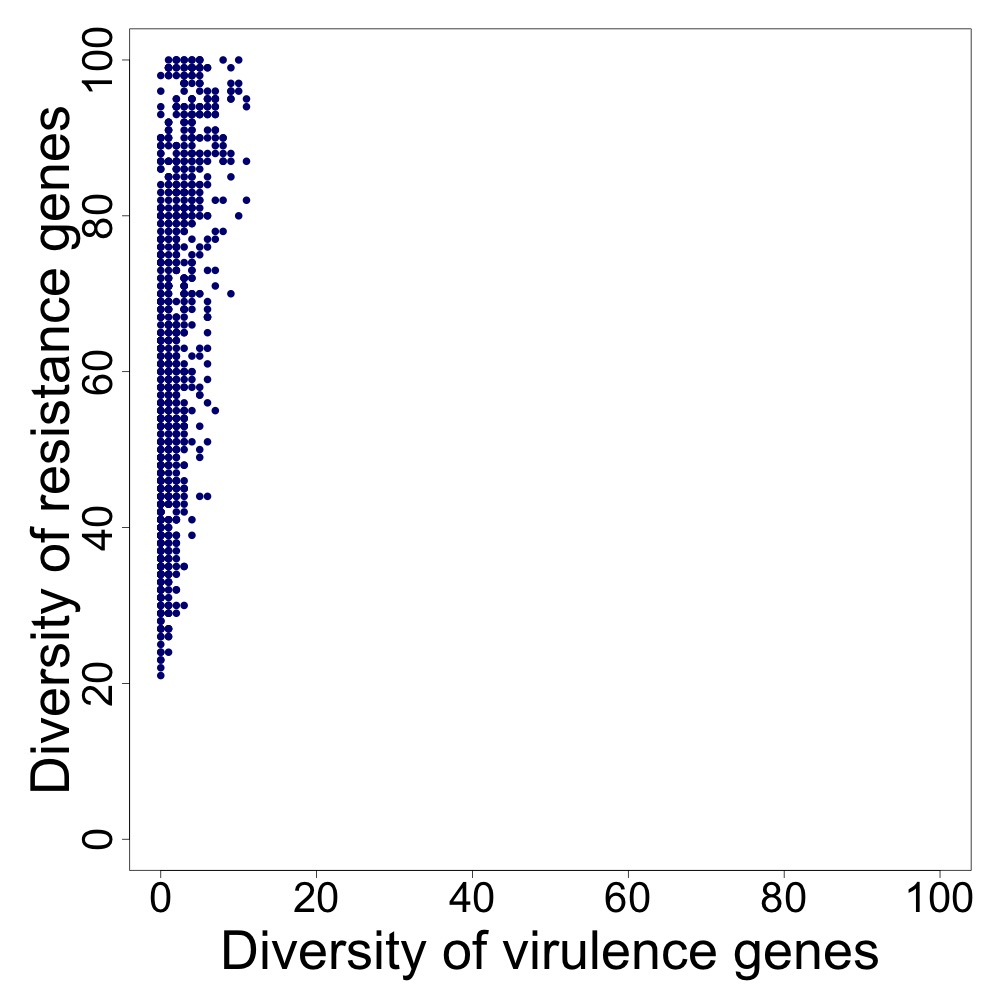 | 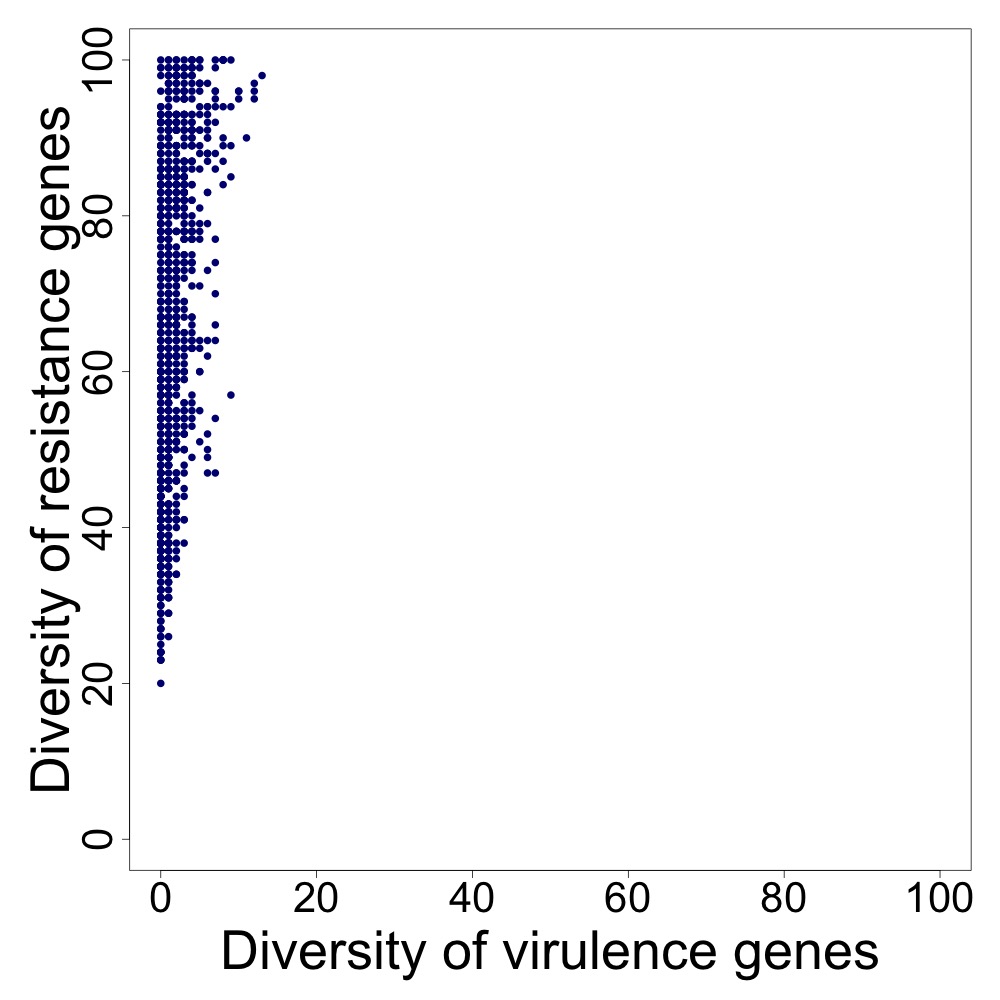 |
| **Network p=0.1** | 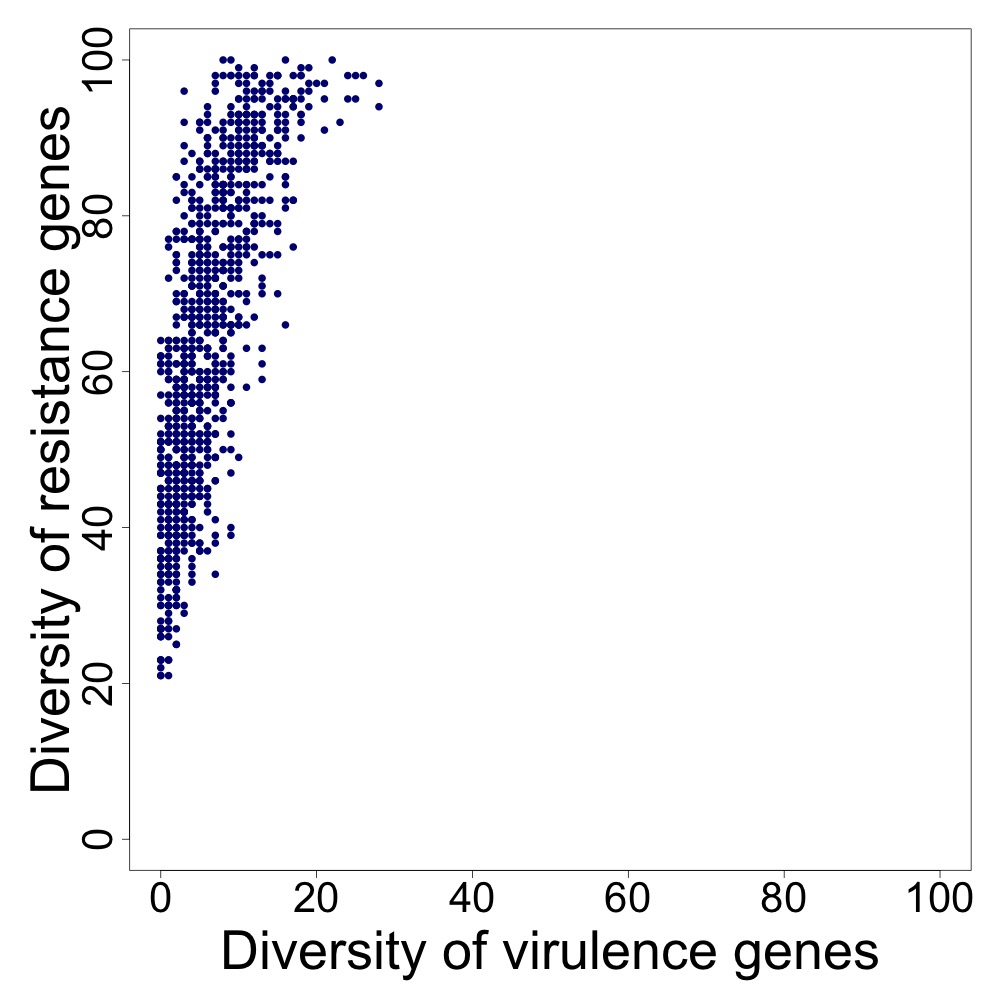 | 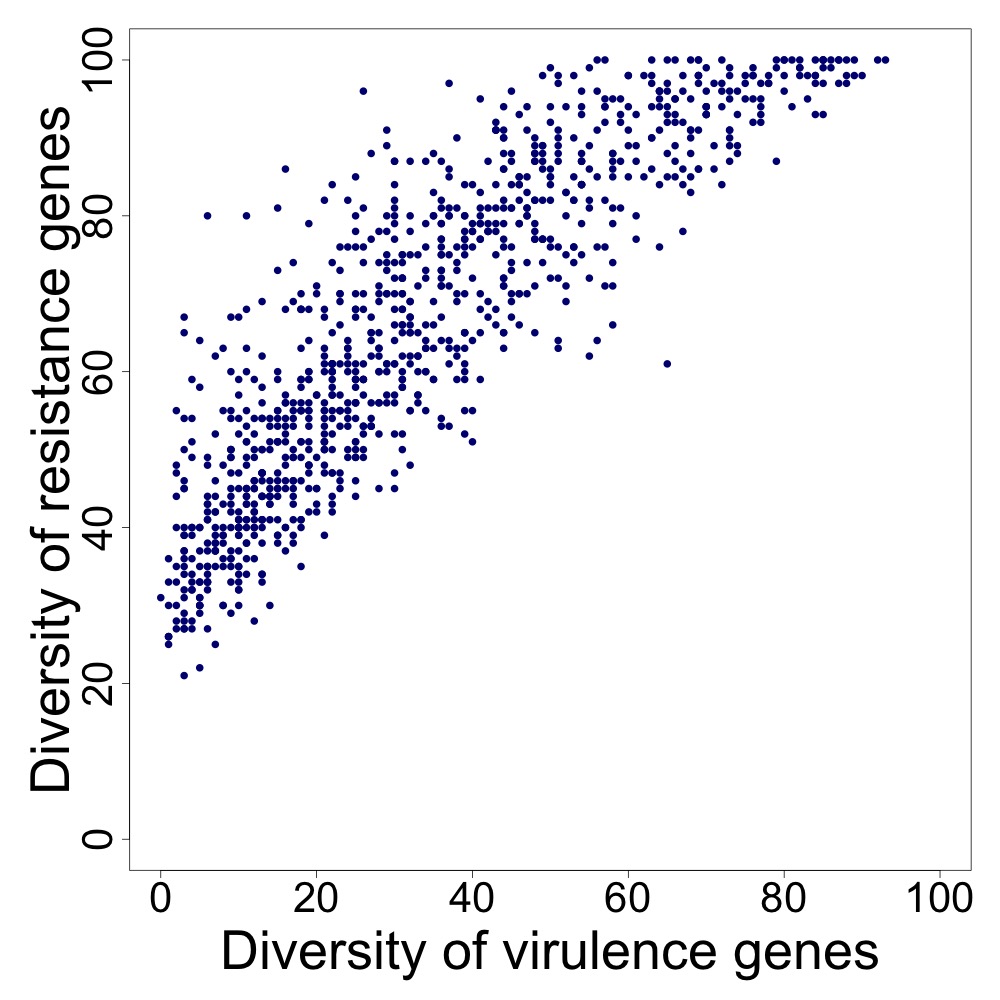 | 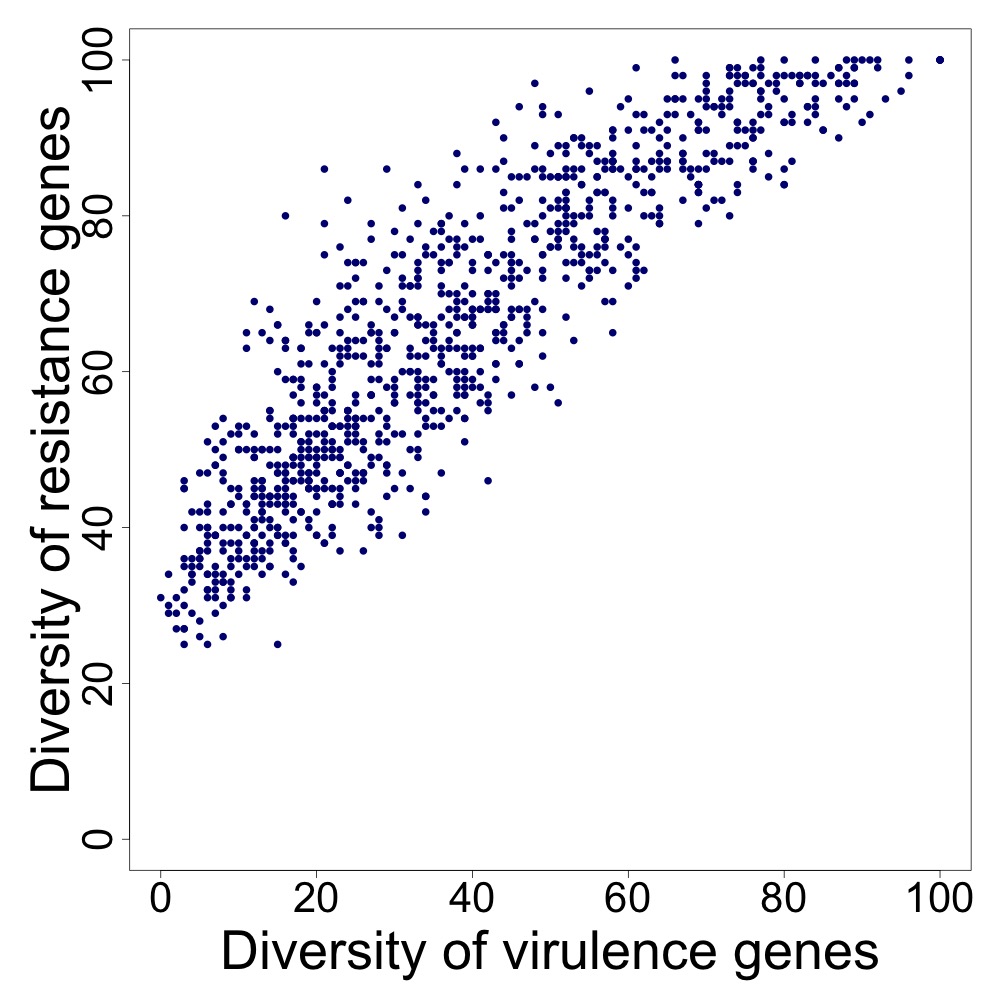 | 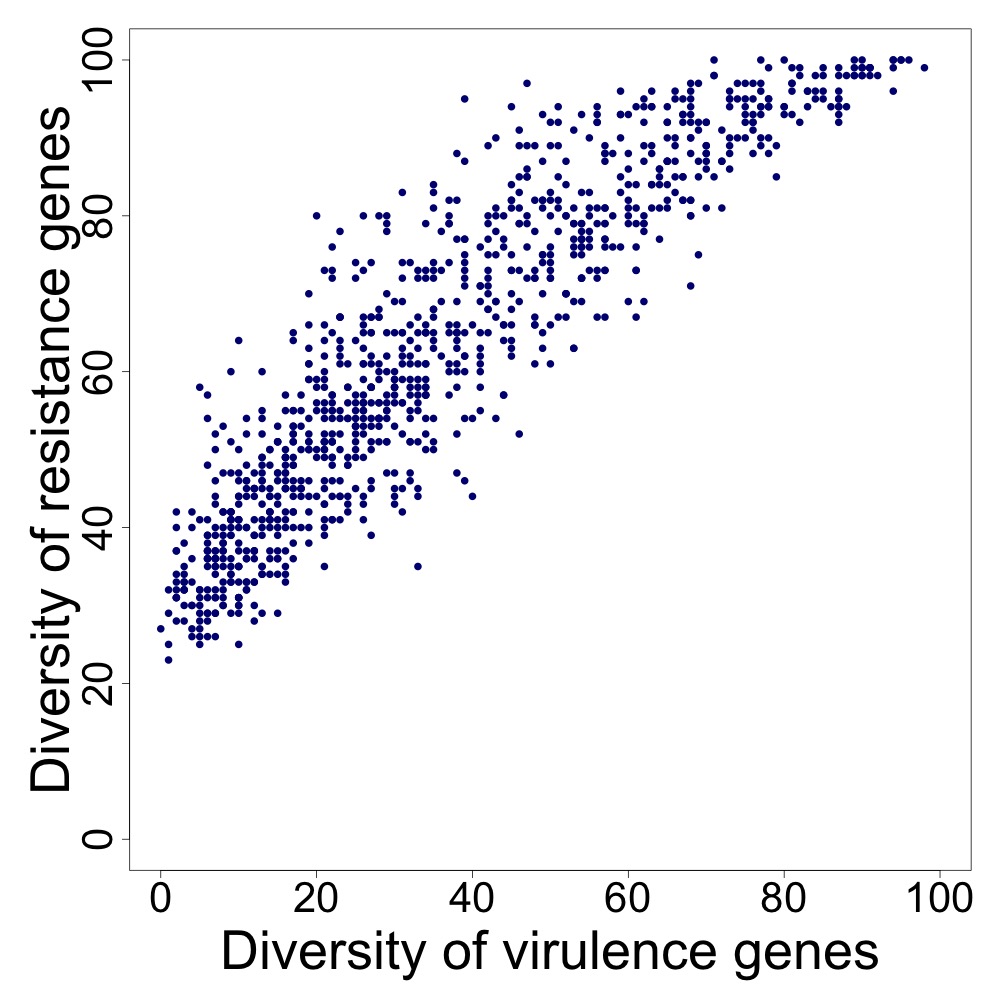 |
| **Network p=0.5** | 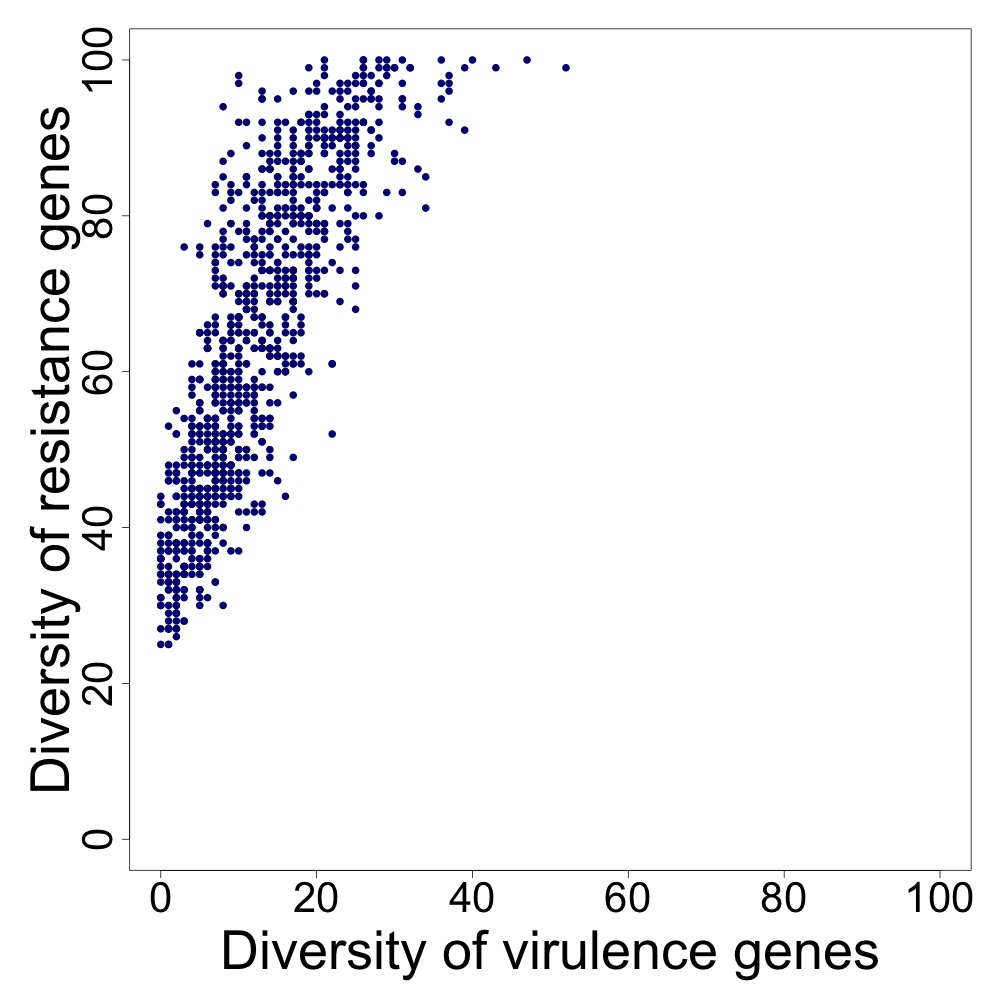 | 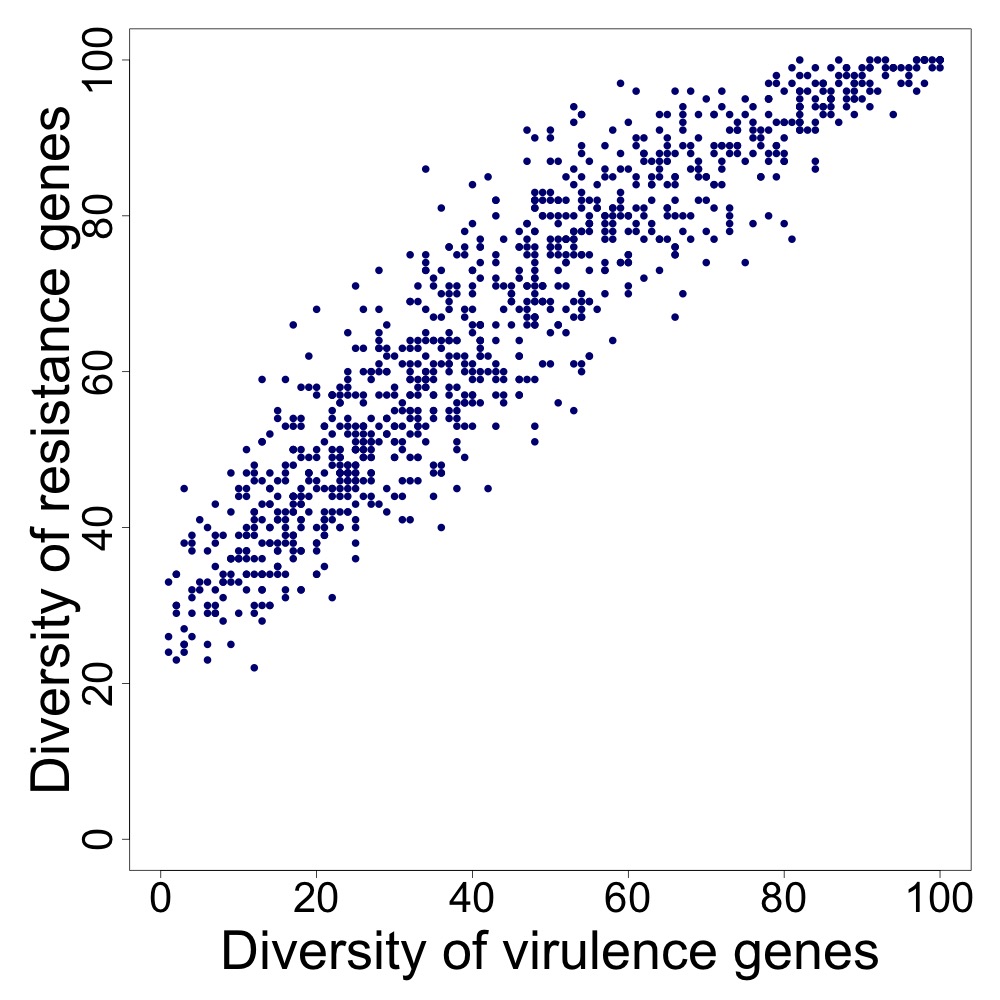 | 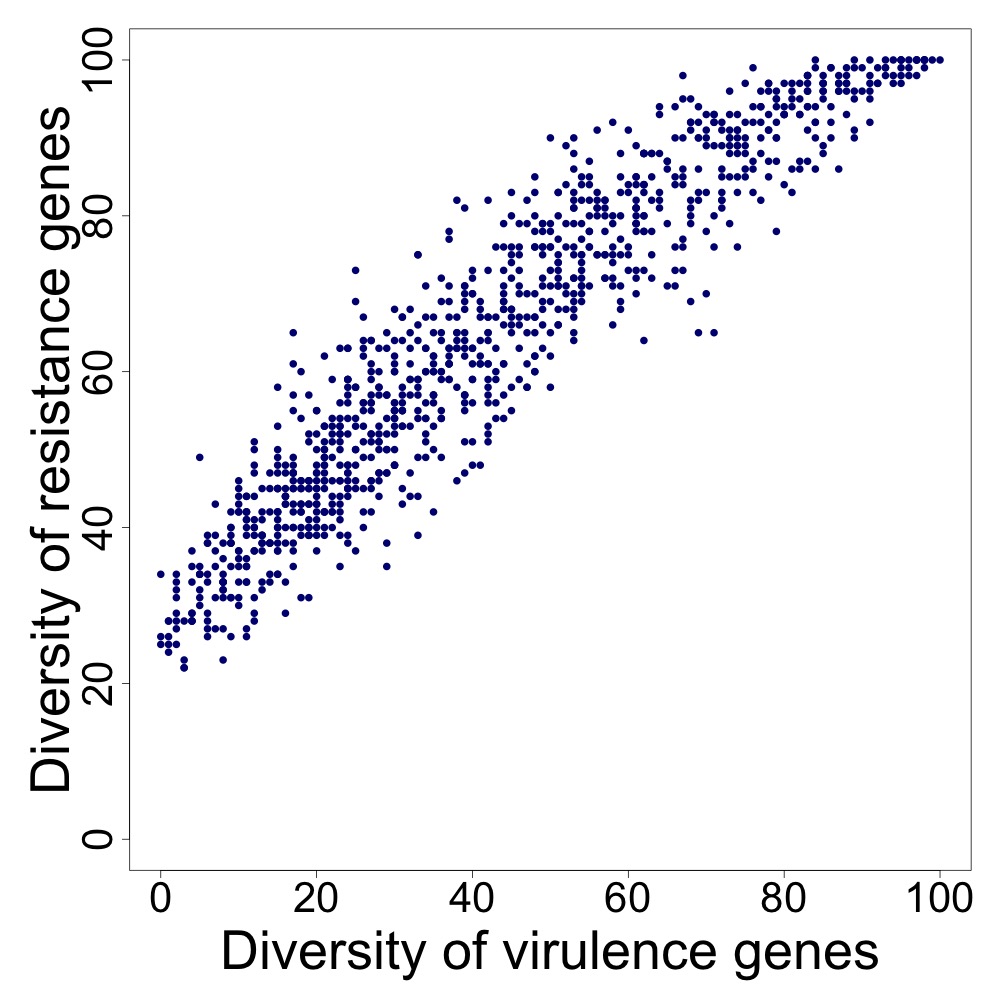 | 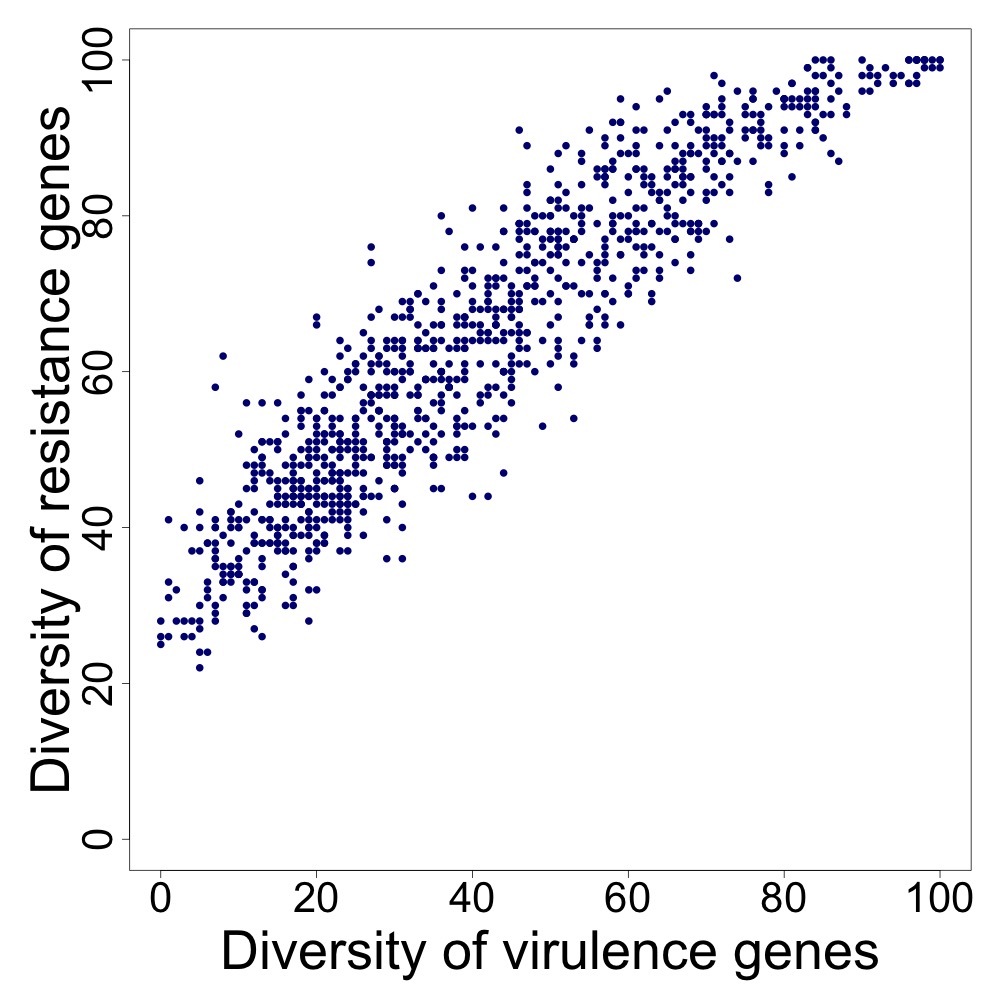 |
| **Network p=1** | 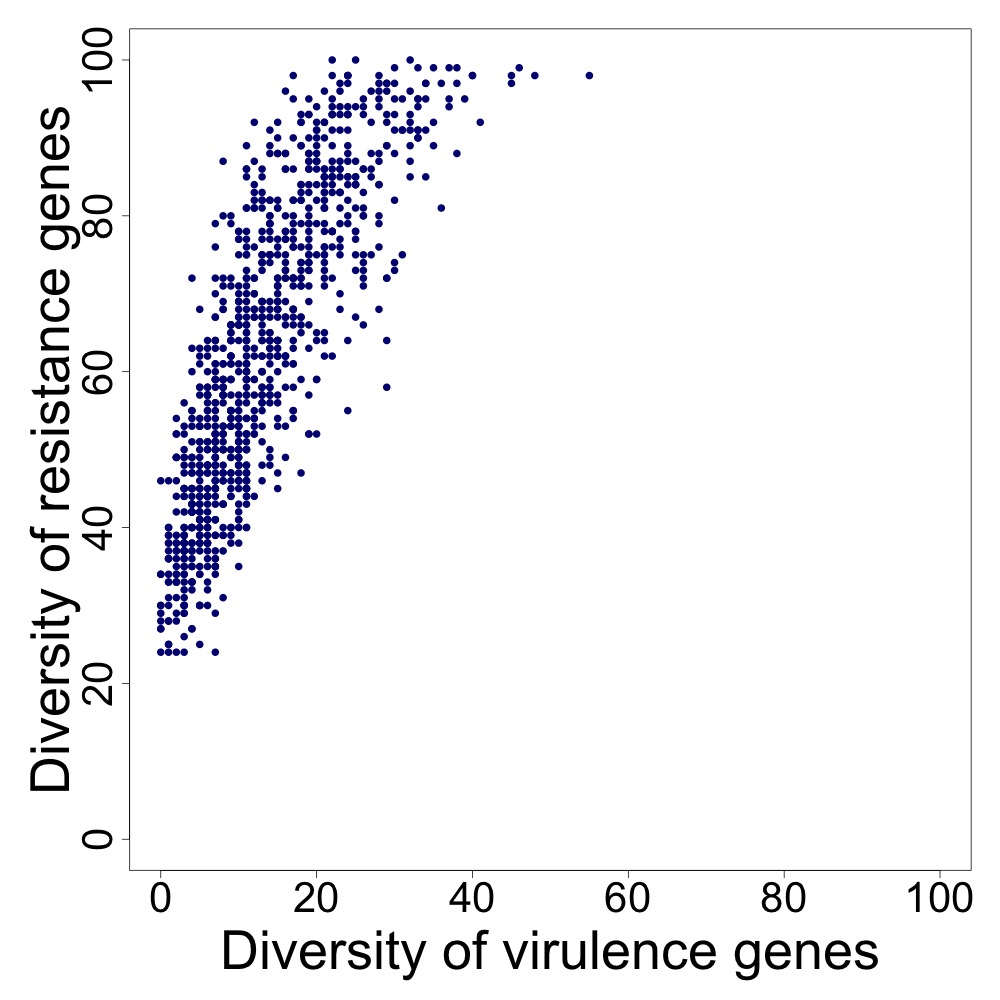 | 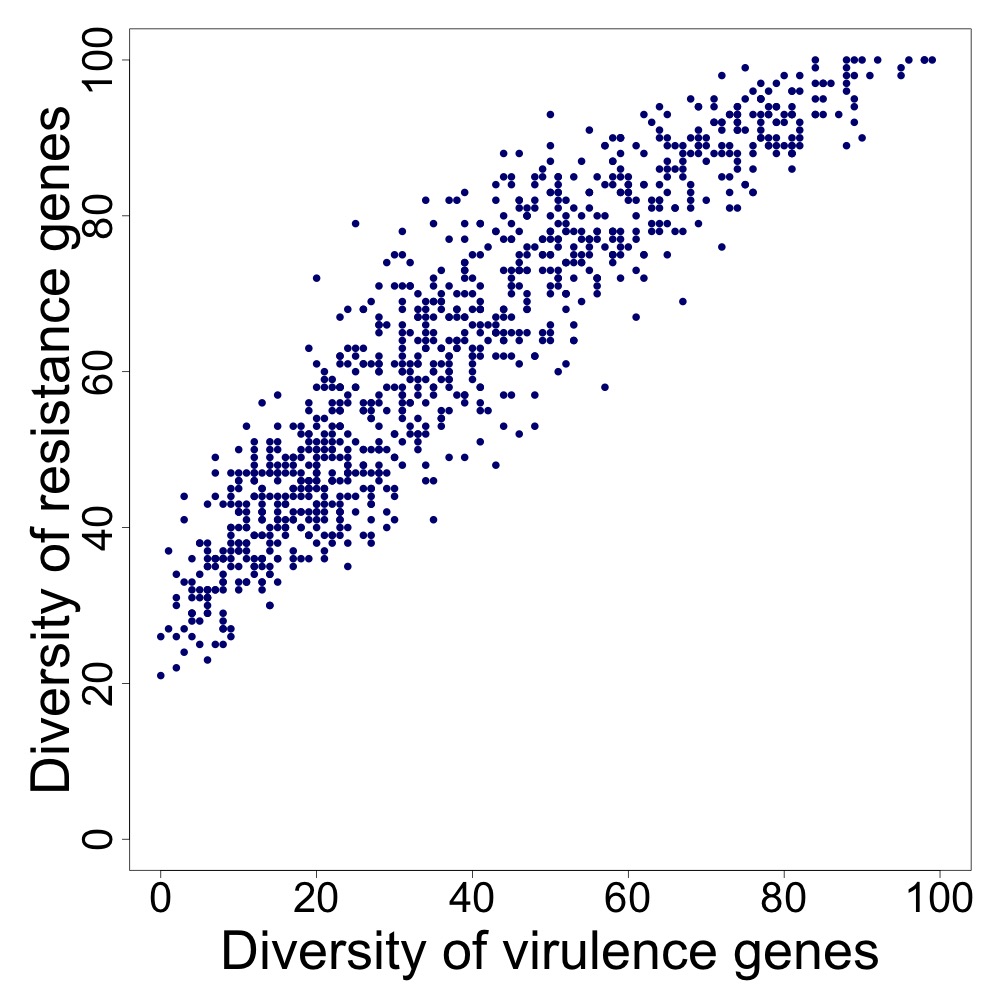 | 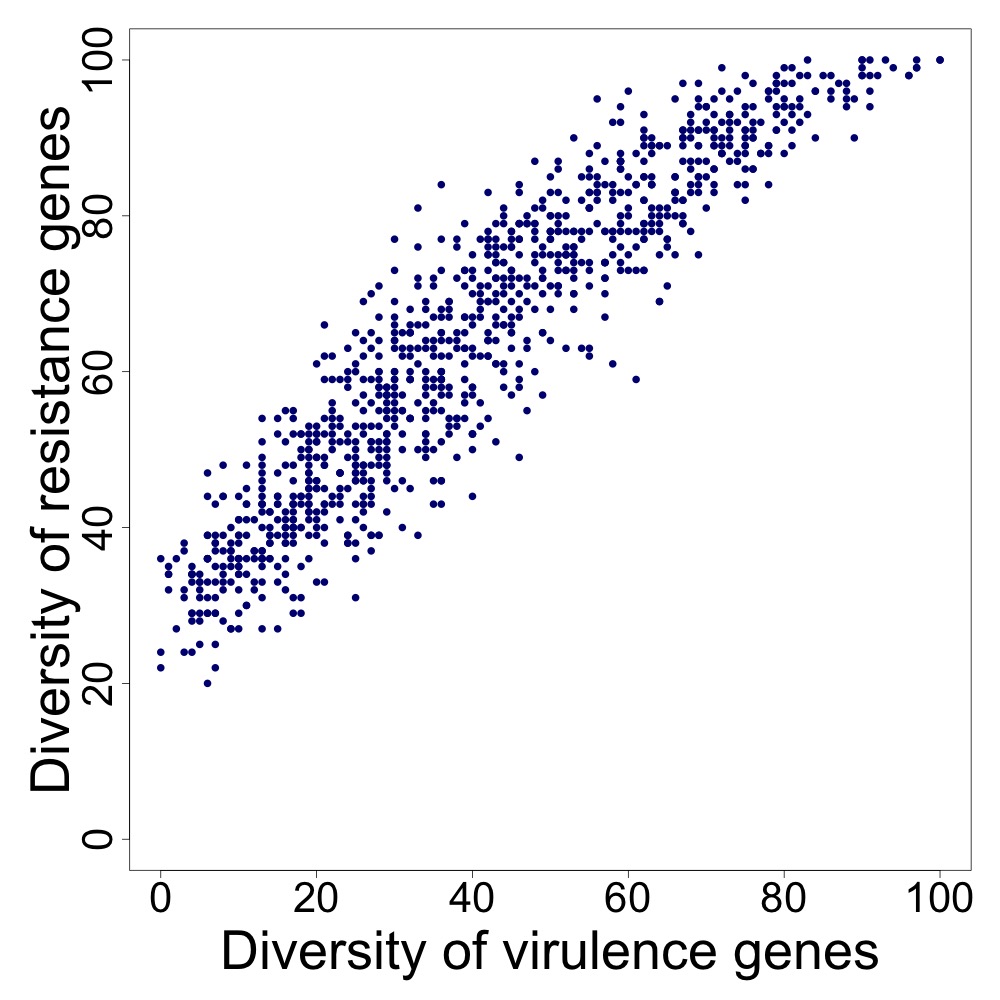 | 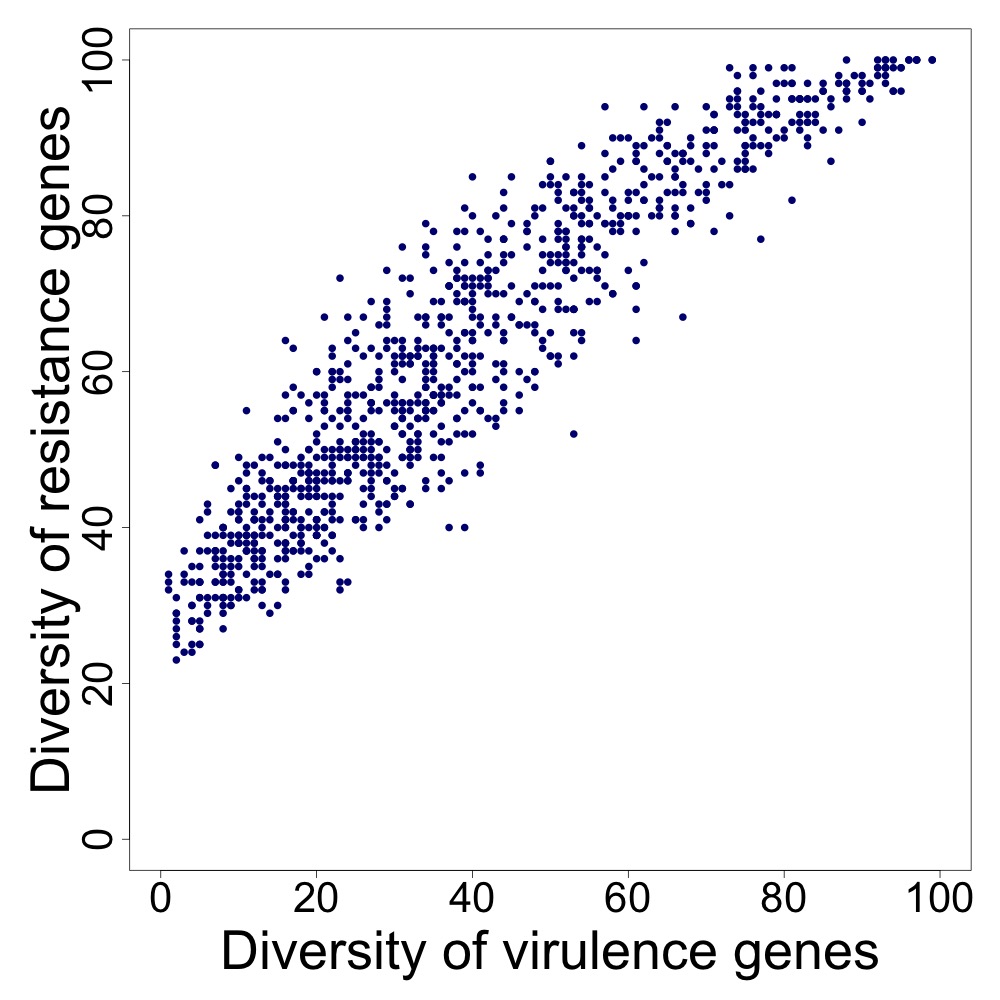 |
| **Supp. Fig. 9.3 - Effect of the networks.** The correlations obtained using a network with p = 0.5 or 1 are stable at cycle 2500, while the correlation obtained with network p = 0.1 is only stable at cycle 7500. Using a network with p = 0, there is no correlation because there is loss of diversity of virulence genes. Correlations obtained for contagion probability of 0.5% and loss rate of 0%. Columns: (1) Cycle 500; (2) Cycle 2500; (3) Cycle 7500; and (4) Cycle 10000. Rows: (1) p=0; (2) p=0.1; (3) p=0.5; and (4) p=1. The vertical axes represent the diversity of resistance genes. The horizontal axes represent the diversity of virulence genes. Each dot represents the diversity of virulence genes present in an individual metagenome. The results were obtained using the parameters described in Methods (Table 2). | | | | |
